# SpoVG and the Kre-ComK Regulatory Module Orchestrate Production of the EPE Toxin in *Bacillus subtilis*

**DOI:** 10.64898/2026.04.02.716078

**Authors:** Sarah Miercke, Kirsten Schaubruch, Sandra Maaß, Anne Katrin Rußeck, Andreas C. Lawaetz, Emma L. Denham, Ralf Heermann, Thorsten Mascher

**Author notes:** Corresponding author: Thorsten Mascher.

## Abstract

Survival of bacteria in their natural habitat requires dynamic responses and adaptation to environmental cues. In *Bacillus subtilis*, one adaptive strategy is cannibalism, a form of programmed cell death during post-exponential development. Cannibalism enhances multicellular differentiation by prolonging or preventing commitment to endospore formation under starvation conditions. *B. subtilis* produces three cannibalism toxins: the sporulation delay protein, the sporulation killing factor, and the epipeptide EPE. Production of the latter is encoded in the *epeXEPAB* operon. Expression of this operon is transcriptionally controlled by the stationary phase regulators Spo0A and AbrB. Here, we demonstrate that EPE production is also post-transcriptionally regulated by two RNA binding proteins, Kre and SpoVG. Deletion of *comK*, the master regulator of competence development, abolished EPE production. This defect was reversed by additionally deleting *kre*. The RNA-binding protein, Kre, binds the *epeX* transcript and acts as a bidirectional ComK repressor, indicating that ComK indirectly regulates EPE biosynthesis via Kre. A second RNA-binding protein, SpoVG, also binds to the *epeX* mRNA. While Kre acts as a negative regulator, SpoVG was essential for EPE production. These findings reveal a novel regulatory connection between competence and cannibalism, expanding our understanding of how programmed cell death is coordinated in *B. subtilis*.

**GRAPHICAL ABSTRACT:** 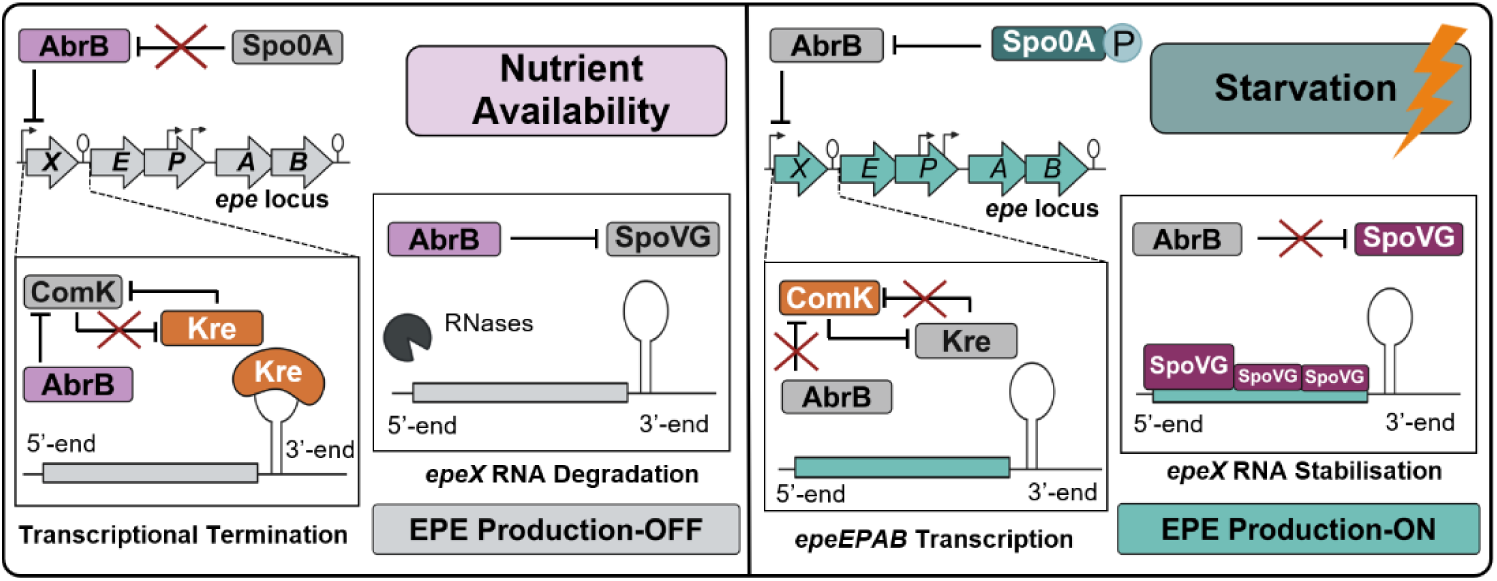

## INTRODUCTION

*Bacillus subtilis* is a Gram-positive, spore-forming bacterium that is continuously subjected to fluctuating and harsh conditions in the soil, which necessitates a dynamic response to ensure its establishment and resilience to competitive pressures (1). This adaptability is driven by sophisticated regulatory networks that control growth, multicellular differentiation and survival. One aspect of this struggle for survival is biological warfare, based on the production of molecules such as antimicrobial peptides, which act against competitors but may also be directed against self. In *B. subtilis*, this process is termed cannibalism, a bacterial form of programmed cell death (PCD) that occurs during post-exponential development (2). Cannibalism enables *B. subtilis* to self-regulate its own population in response to nutrient-limiting conditions, by sacrificing a subpopulation to postpone or even prevent the highly energy-demanding process of endospore formation (3). In addition to the cannibalism toxins sporulation delay protein (SDP) and the sporulation killing factor (SKF) (2,4), *B. subtilis* also produces the epipeptide EPE (5).

EPE is a ribosomally synthesised antimicrobial peptide, which is encoded in the *epeXEPAB* biosynthesis locus (Figure 1a) (6). The pre-pro-peptide EpeX (49 aa) is encoded by the *epeX* gene and post-translationally modified by the radical S-adenosyl-L-methionine (SAM) epimerase EpeE, resulting in the conversion of L-valine_4_ and L-isoleucine_12_ to their D-forms (Figure 1a) (7). The activity of EpeE relies on an iron-sulphur ([4Fe-4S]) cluster coordinated by three cysteines and an exchangeable SAM in its active site, which is essential for catalysing the epimerisation reaction (8,7). The resulting pre-peptide is subsequently processed and exported, presumably by the membrane-anchored signal peptidase EpeP, in order to secrete the 17 amino acids linear mature toxin EPE (Figure 1a) (7). The ATP-binding cassette (ABC) transporter EpeAB provides autoimmunity against intrinsically produced EPE, but does not protect cells from externally applied toxin (Figure 1a) (5). EPE primarily targets *B. subtilis*, although it can also affect closely related Gram-positive species (6,9). Its antimicrobial activity derives from its ability to severely disrupt cytoplasmic membrane integrity by dissipating the membrane potential through permeabilization, which is accompanied by a rapid decrease in membrane fluidity and the formation of lipid domains (5).

**Figure 1:**
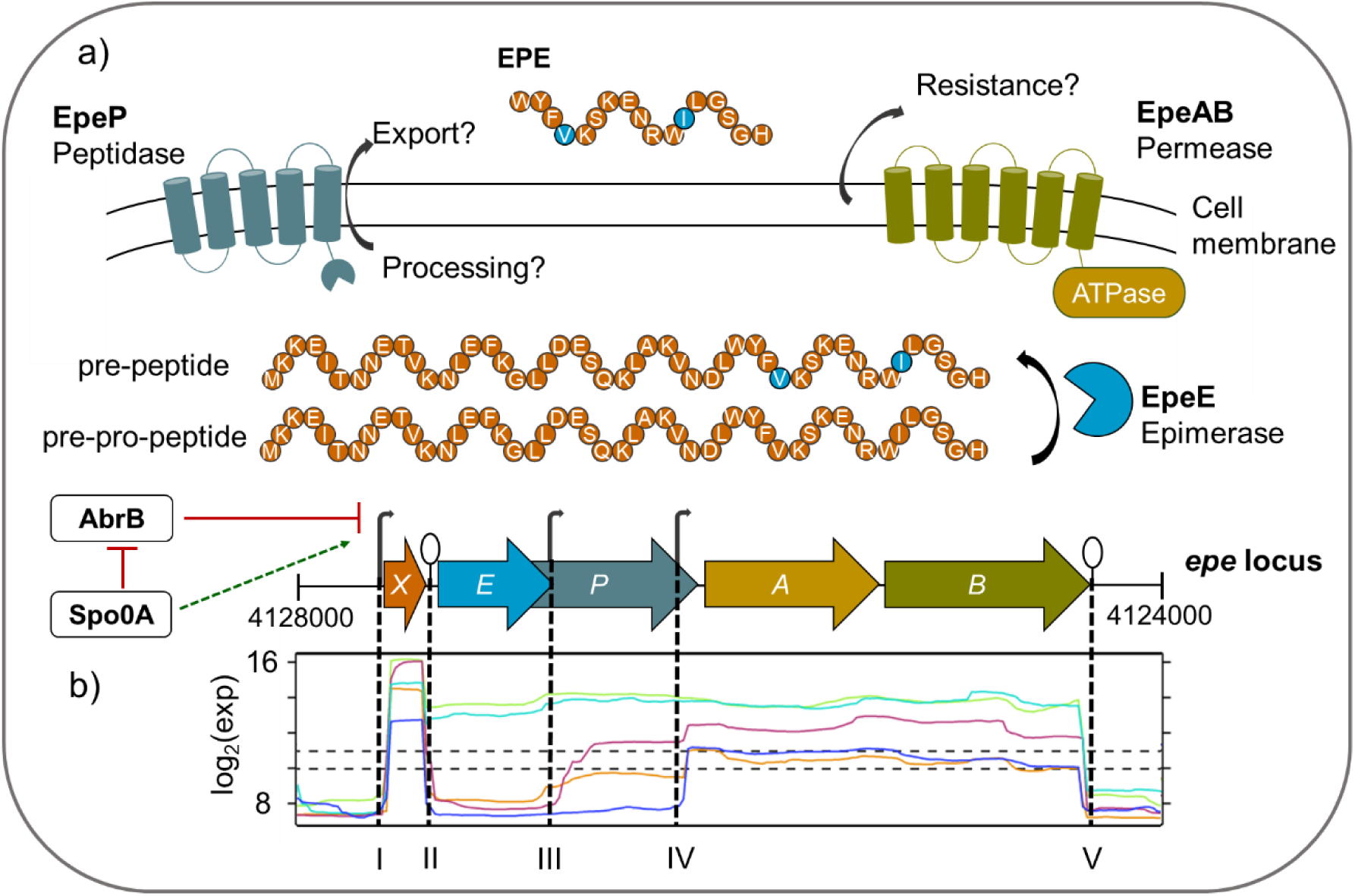
Schematic representation of the *epe* locus, EPE biosynthesis, and *epe* expression in *B. subtilis.* **a)** Expression of *epeXEP* is driven by P*_epeX_* and transcriptionally regulated by AbrB repression. Upon initiation of sporulation Spo0A indirectly activates P*_epeX_* due to repression of AbrB. Expression of *epeXEP* leads to the production of the ribosomally synthesised antimicrobial peptide EPE, whereby *epeX* encodes for a pre-pro-peptide of 49 aa (orange), which is post-translationally modified by the radical SAM epimerase EpeE (light blue), introducing to epimerisation’s at L-Val_4_ and L-Ile_12_ (highlighted in light blue). The resulting pre-peptide is subsequently processed and exported by the membrane anchored peptidase EpeP (dark blue) to release the 17 aa linear mature toxin, called EPE. Autoimmunity against intrinsic EPE is ensured by the expression of the ABC transporter EpeAB (yellow/green). The scheme was adapted from Popp et al., 2021 with modifications. **b)** Mapping of mRNA abundance based on comprehensive tiling array data indicates 3 upshifts of mRNA abundance. While the first coincides with the *epeX* promoter **(I)**, the following two correspond to the already mapped promoters located within *epeP* (P*_epeA1/2_*) **(III, IV)**. Moreover, a strong growth condition-dependent downshift at the 3’-end of *epeX* was reported by analysing over 200 different growth conditions **(II)**. Conditions: mid-log growth in M9 minimal medium with malate (turquois), shift from CH minimal medium to SM sporulation medium 2 h (light green) and 6 h (orange, blue) after shift, biofilm growth after 36 h in MSgg medium at 30°C (red). In contrast to the growth independent termination at the 3’-end of *epeB* **(V)**, growth-dependent changes in particular *epeEP* mRNA abundance cannot exclusively explained by the mapped terminator **(II)** and rises the hypothesis of post-transcriptional regulation at the 3’-end of *epeX*. The expression profile was derived from (10–12).

Damage to the membrane activates the cell envelope stress-sensing two-component system LiaRS, leading to strong induction of the *liaIH* operon, whose products confer resistance to extracellular EPE (5). The P*_liaI_*-*luxABCDE* (*lux*) reporter therefore serves as a sensitive and specific readout for the presence of active EPE (5). Importantly, additional deletion of the toxin autoimmunity system, encoded by *epeAB*, further increases cellular susceptibility to EPE and thereby enhances the sensitivity of the P*_liaI_*-*lux* reporter, making a strain containing Δ*epeAB* and the P*_liaI_*-*lux* reporter a powerful tool for assessing EPE production (Figure S1) (13).

EPE production is tightly regulated at the level of *epeXEP* expression. Transcription of the *epe* operon is driven by the *epeX* promoter, P*_epeX_*, which is indirectly activated by Spo0A via its repression of the global transition state regulator AbrB at the onset of stationary phase (Figure 1a) (6). In addition, two constitutive promoters (P*_epeA1;2_*) within *epeP* are postulated to facilitate EpeAB-mediated autoimmunity prior to the production of the toxin (10). Transcript quantification in the course of a comprehensive tiling array study indicated a strong upshift of mRNA abundance at the 5’-end of *epeX* (Figure 1b (I)) as well as two slighter upshifts corresponding to the location of P*_epe1;2_* (Figure 1b (III-IV)) (10). Notably, a strong growth condition-dependent downshift of mRNA abundance occurs at the 3’-end of *epeX* (Figure 1b (II)) (11,10). While growth under nutrient starvation and conditions promoting Spo0A activation maintain a relatively high level of transcription covering the entire *epe* locus (light green, and green in Figure 1b), biofilm formation and commitment to sporulation (pink, and purple in Figure 1b) resulted in a significant reduction of the *epeEP* transcript (Figure 1b (II)) (10,11). The strong downshift at the 3’-end of *epeB* is caused by a rho-independent terminator (Figure 1b (V)) (12). Together, these expression data suggest additional post-transcriptional regulation at the 3’-end of *epeX* under nutrient-limiting conditions, as exemplified by our recent study showing that the Fur regulated small regulatory RNA FsrA positively influences EPE production by base-pairing with the intergenic region between *epeX* and *epeE* under iron starvation (13).

In fluctuating and nutrient-limited environments, division-of-labour is a means to diversify an isogenic population by initiating multiple differentiation strategies at the same time (14). In *B. subtilis*, these adaptive responses are primarily governed by the master regulator of sporulation, Spo0A, and quorum sensing-dependent mechanisms that assess population density. Together, they orchestrate cell fate decisions in response to nutrient limitations (Figure 2) (15,14). Upon sensing nutrient depletion, the phosphorelay, a signal-transduction cascade, is initiated by the histidine kinases KinA-E (16). This pathway sequentially phosphorylates Spo0F and Spo0B, ultimately resulting in the phosphorylation of the response regulator Spo0A (Figure 2 (I)) (17,18). Activated Spo0A (Spo0A∼P) represses the transition state regulator AbrB, thereby initiating subsequent post-exponential differentiation processes (Figure 2 (I)) (19). At low levels of Spo0A∼P, individual cells can differentiate for example into competent cells capable of DNA uptake, or engage in cannibalistic behaviour to delay sporulation (Figure 2 (II-III)) (20).

**Figure 2:**
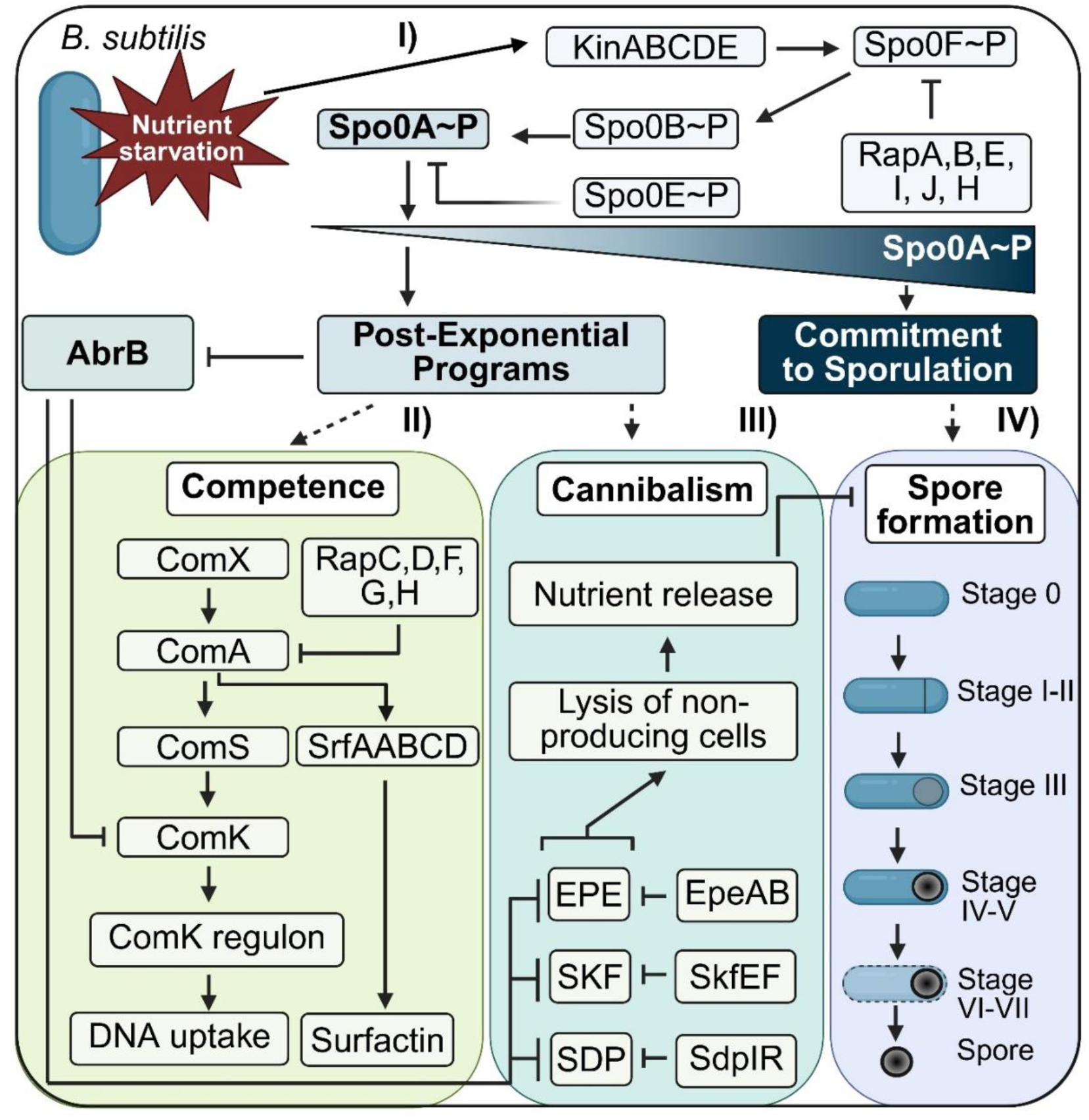
Nutrient starvation induced differentiation strategies in *B. subtilis*. **I)** In response to nutrient limitation, *B. subtilis* initiates a phosphorylation cascade involving the kinases KinA-E, which sequentially activate Spo0F, Spo0B, and ultimately Spo0A. This phosphorelay system serves as a master regulatory mechanism coordinating alternative cell fates. Activation of this pathway leads to the repression of the transition state regulator AbrB and initiates post-exponential programs. **II)** At low levels of phosphorylated Spo0A (Spo0A∼P), cell may differentiate into the competence state, becoming capable of DNA uptake. **III)** Similarly, low Spo0A∼P levels can trigger production of cannibalism toxins that lyse non-resistant sibling cells to delay sporulation and recycle nutrients. **IV)** High levels of Spo0A∼P drive the irreversible commitment to endospore formation, ensuring survival under prolonged stress.

In contrast, a high level of Spo0A∼P results in the final commitment to sporulation, resulting in the formation of highly resilient endospores. Sporulation is a complex, tightly regulated process involving sequential stages and compartment-specific sigma factors that ensure precise temporal and spatial coordination between the forespore and mother cell (Stage 0 to Stage VII, Figure 2 (IV)) (21–23). Although endospores can survive extreme conditions for long periods, they are metabolically dormant and thus incapable of growth, replication, or direct competition with actively dividing bacteria for nutrients and position within their ecological niche (24,25). Moreover, endospore formation is highly energy-consuming and irreversible, making it a survival strategy of last-resort, when all other options for adaptation have been exhausted (26).

Together, these alternative cell fates - competence, cannibalism, and sporulation underscore the capacity of *B. subtilis* to balance survival, competition, and population heterogeneity in fluctuating environments (27,20). Reflecting this regulatory complexity, although EPE production is transcriptionally controlled by AbrB and Spo0A (6), expression data suggested the presence of additional post-transcriptional control (Figure 1b). We therefore, investigated the regulation of EPE biosynthesis and uncovered two RNA-binding proteins that fine-tune its production, with one acting as a regulatory link between competence and cannibalism.

## MATERIAL AND METHODS

### Bacterial Strains and Growth Conditions

*Bacillus subtilis* and *Escherichia coli* were cultivated at 37 °C and 220 rpm agitation in lysogeny broth (LB) [1% tryptone (w/v), 0.5% (w/v) yeast extract, and 1% (sodium chloride], Difco sporulation media (DSM) [0.8% (w/v) nutrient broth, 0.1% (w/v) potassium chloride, 1mM magnesium sulphate, 10 µM manganese chloride, 1 µM iron(II) sulphate x 7 H_2_O, and 0.5 mM calcium chloride], or chemical defined MNGE [88.2% 1X MN medium (1.36% (w/v) dipotassium phosphate × 3 H_2_O, 0.6% (w/v) monopotassium phosphate, 0.1% (w/v) sodium citrate × H_2_O), 1.9% (w/v) glucose, 0.19% (w/v) potassium glutamate, 0.001% (w/v) ammonium ferric citrate, and 0.035% (w/v) magnesium sulphate]. For solidification of agar plates 1.5% (w/v) agar-agar were applied. All *B. subtilis* and *E. coli* strains used in this study were listed in (Table S1). Selection of *B. subtilis* strains harbouring a resistance cassette was carried out by applying chloramphenicol (5 µg mL^-1^), kanamycin (10 µg mL^-1^), spectinomycin (100 µg mL^-1^), phleomycin (2 µg/mL), or for macrolide-lincomycin-streptogramin B (MLS) resistance 1 µg mL^-1^ erythromycin combined with 25 µg/mL lincomycin. *E. coli* strains carrying plasmids were selected by using kanamycin (50 µg mL^-1^), chloramphenicol (35 µg mL^-1^), or ampicillin (100 µg mL^-1^). Transformation of *B. subtilis* and *E. coli* was performed according to (28,29).

### DNA Manipulation

The construction of plasmids was based on standard cloning procedures, as described previously (28). Enzymes used in this study were purchased from New England Biolabs and the manufacturer’s instructions have been followed (NEB, Ipswich, MA, USA). DNA fragment purification and plasmid isolation was performed with the HiYield PCR Gel Extraction/PCR Clean-up Kit, or HiYield Plasmid Mini-Kit (Süd-Laborbedarf GmbH, Gauting, Germany), respectively, according to the corresponding protocols, and genomic DNA was isolated by alkaline lysis (29). Incorporation of luciferase based reporter constructs into the genome was performed by applying pBS3C-*lux*, or pBS3C*αlux* based plasmids (30,31), and marker less genomic modifications were achieved by applying pJOE8999 derived plasmids (15). The commercially available pET28a expression vector was used for generation of heterologous protein overexpression plasmids. Verification of all constructs and genomic modifications were carried out by colony PCR and sequencing. Plasmids and primers applied in this study are listed in Table S2 and Table S3 respectively.

### Reporter Gene Assay

Luciferase activity of strains harbouring respective reporter fusions was assayed as previously detailed described (31). Briefly, cultures were grown in LB media with appropriate antibiotics overnight, and day cultures without antibiotics were inoculated 1:100 in DSM medium until and OD_600_ of 0.2-0.4. Subsequently, the cell suspension was diluted with fresh DSM media to an OD_600_ of 0.05, and 100 µL were transferred into a 96-well plate (black wells, clear bottom, Greiner Bio-One, Frickenhausen, Germany), and placed in BioSpa 8 automated incubator (BioTek, Winooski, USA), which is under control of the software Gen5^TM^ (BioTek, Winooski, USA). Incubation of samples occurred at 37 °C and OD_600_ as well as relative luminescence units (RLU) were monitored every 9 min for 18 h. Assays were performed at least as biological and technical triplicate.

### *In vitro* transcription of RNA fragments

*In vitro* transcription was carried out using a PCR-amplified template derived from *B. subtilis* DK1042 wild-type chromosomal DNA. The forward primer used for amplification included the T7 RNA polymerase promoter sequence (5’-CTAATACGACTCACTATAGGGAGA-3’) at its 5’-end to enable transcription. Approximately 1 µg of the purified PCR product was used for in vitro transcription in a 20 µl reaction containing 2 µL 10X ribonucleotide solution mix, 2 µL 10X T7 Polymerase transcription buffer, 2 µL T7 polymerase (N0466 and M0251, New England Biolabs, Ipswich, Massachusetts, USA, respectively) and 1 µL RNase inhibitor (N2511, Promega Corporation, Madison, Wisconsin, USA). The reaction was incubated at 37 °C for 2 hours. Subsequently, 2 µL of DNase I was added, and the reaction was incubated for an additional 30 minutes at 37 °C to remove the DNA template. The transcription reaction was terminated by the addition of 2.5 µL of 0.2 M EDTA (pH 8.0). For RNA precipitation, 2.5 µL of 4 M LiCl and 75 µL of cold ethanol were added to the reaction mixture, followed by incubation at -20 °C overnight. The precipitated RNA was collected by centrifugation (15 minutes at 13,000 rpm, 4 °C), and the supernatant was carefully removed. The RNA pellet was washed with 75 µL of 70% (v/v) ethanol and centrifuged again under the same conditions. After discarding the supernatant, the pellet was air-dried and resuspended in 100 µL of RNase-free H_2_O containing 1 µL of RNase inhibitor. The RNA probe was incubated at 37 °C for 30 minutes and stored at -80 °C.

### Protein Production and Purification

Overexpression of Kre and SpoVG was conducted by applying the expression strain *E. coli* BL21 harbouring pET28b(+) derived plasmids with the respective protein sequence. An overnight culture was used to inoculate 1.8 L LB broth supplemented with 50 µg mL^-1^ kanamycin to an OD_600_ 0.1. The culture was grown at 37 °C and aeration until OD_600_ 1.0 before induction with IPTG to a final concentration of 1 mM. Subsequently, expression was performed at 30°C and 150 rpm for 20 h and cells were harvested by centrifugation. The cell pellet was then stored at -20 °C. Protein purification was performed according to (32) with small changes. In brief, the cell pellet was resuspended in 20 mL lysis buffer [20 mM Tris-HCl, 5 mg mL^-1^ lysozyme, 0.01 mg mL^-1^ DNase I, and 200 µL Halt^TM^ Protease Inhibitor Cocktail (Thermo Fisher Scientific, Massachusetts, USA)] and incubated for 45 min at 4 °C, before 3 times sonication for 1 min with the Sonopuls GM 70 [amplitude: 100%, Cyc: 0.6, (Bandelin, Berlin, Germany)]. Following centrifugation at 4 °C and 20,000 rpm for 45 min the protein lysate was loaded on HisTrap^TM^ FF column (Cytiva, Uppsala, Sweden) for immobilized metal affinity chromatography (IMAC). After washing with His-wash buffer [50 mM Tris-HCl (pH 8.0), 300 mM NaCl, 10 mM imidazol], the proteins were eluted by applying His-elution buffer [50 mM Tris-HCl (pH 8.0), 300 mM NaCl, 500 mM imidazol]. Moreover, this was followed by size exclusion chromatography (SEC) using Superdex^TM^ 200 Increase 10/300 GL column (GE Healthcare, Chicago, USA) and SEC buffer [50 mM Tris-HCl (pH 8.0), 300 mM NaCl]. For long time storage samples were immediately supplemented with 19% (v/v) glycerol, frozen in liquid nitrogen and stored at - 80 °C. Protein concentration was determined by Bradford assay using 5X Bradford reagent (SERVA Electrophoresis GmbH, Heidelberg, Germany) according to the manufacturer’s recommendation (33). Protein separation was performed by sodium dodecyl sulphate polyacrylamide gel electrophoresis (SDS-Page) using a discontinuous gel system (Bio-Rad, Hercules, CA, USA) with a 12.5% (w/v) polyacrylamide separating gel and 4% (w/v) polyacrylamide stacking gel of 1 mm thickness. For the separating gel a 10 mL stock solution (4.2 mL 30 % (v/v) acrylamide and bis-acrylamide solution [37.5:1], 2.5 mL 1.5 M Tris-HCl [pH 8.8], 0.1 mL 10% (w/v) SDS, and 3.2 mL deionised H_2_O) was prepared. Polymerisation was carried out by the addition of 50 µL 10% (w/v) ammonium peroxide sulphate (APS), and 10 µL N,N,N’,N’-tetramethlyethylenediamine (TEMED). The gel solution was poured to fill approximately two-thirds of the casting chamber. The gel was then overlaid with water and allowed to polymerise for ∼30 min. Subsequently, the stacking gel was prepared from 0.67 mL 30% (v/v) acrylamide and bis-acrylamide solution [37.5:1], 0.4 mL 1.5 M Tris-HCl [pH 6.8], 50 µL 10% (w/v) SDS, and 3.85 mL deionised H_2_O). After addition of 50 µL 10% (w/v) APS and 8 µL TEMED, the stacking gel was poured onto the polymerised separating gel and a comb inserted for well formation. Electrophoresis was carried out in SDS running buffer (25 mM Tris-HCl [pH 8.0], 192 mM glycine, 0.1% SDS) at a constant voltage of 100 V for 1.5 hours. Following electrophoresis, proteins were fixed in 50% (v/v) acetone for 10 minutes and proteins were visualised by staining with Coomassie solution (45% (v/v) methanol, 10% (v/v) acetic acid, 2.5 g L^-1^ Coomassie G-250) for 30 minutes. Destaining was achieved by washing the gels with deionised water 3-5 times, followed by overnight washing on a rocking platform. A paper towel was placed in the container to efficiently absorb excess dye.

### Blue Native Polyacrylamide Gel Electrophoresis

Blue native PAGE was performed using the SERVA native PAGE system (SERVA Electrophoresis GmbH, Heidelberg, Germany) according to the manufacturer’s protocol. In brief, samples were prepared 1:2 in 2X sample buffer [1 M 6-amino-capronacid, 100 mM BisTris (pH 7.0), 100 mM NaCl, 20% (w/v) glycerine, and 0.1% (w/v) SERVA Blue G250] and loaded on a SERVAGel^TM^ N 3-12% (w/v) gradient gel. Gel electrophoresis was performed by applying 1X native cathode buffer [50 mM tricine, 15 mM BisTris, and 0.002% (w/v) SERVA Blue G-solution] and 1X native anode buffer (50 mM BisTris-HCl, pH 7.0) at initially 50 V (constant) for 10 min, followed by 200 V (constant) for approximately 90 min. Subsequently, the gel was fixated in 20% (v/v) trichloroacetic acid for 30 min. Staining was carried out in 50% (v/v) staining solution I (0.2% (v/v) SERVA Blue R and 90% (v/v) ethanol) and 50% (v/v) staining solution II (20% (v/v) acetic acid) for 30 minutes. Following, the gel was washed in destaining solution [20% (v/v) ethanol, 5% (v/v) acetic acid, and 1% (w/v) glycerine] for 2 times 60 minutes and pictures were taken by FUSION FX Imager (Vilber Lourmat Deutschland GmbH, Eberhardzell, Germany).

### Electro Mobility Shift Assay

Electro mobility shift assays (EMSA) were performed to analyse protein RNA interactions. First, RNA was denaturised at 70°C for 10 min, followed by 5 min cooling down on ice. 5 pmol of *in vitro* transcribed *epeX* RNA was mixed with increasing concentrations of purified SpoVG in 5 x RNA binding buffer [50 mM Tris-HCl (pH 8.0), 50 mM NaCl, 5 mM MgCl_2,_ 5 mM DTT, 250 µg/mL BSA]. Subsequently, the RNA-protein mix was incubated for 30 min at 37°C. Following 6 x RNA loading dye [30% (v/v) glycerol, 0.25% (w/v) xylene blue, and 0.25% (v/v) bromophenol blue] was added to the reaction and 20 µL of the sample were loaded on non-denaturing polyacrylamide gels [6% (v/v) acrylamide and bis-acrylamide solution (37.5:1), 1x TBE [(89 mM Tris-HCl, pH 8.0), 89 mM boric acid, and 2 mM EDTA (pH 8.0)], 0.06% (w/v) APS, and 0.12% TEMED] and gel electrophoresis was conducted in 1 x TBE buffer. Subsequently, the gel was stained for 30 min in ethidium bromide water bath and then RNA was visualised by FUSION FX Imager (Vilber Lourmat Deutschland GmbH, Eberhardzell, Germany).

### Surface Plasmon Resonance (SPR) Spectroscopy

SPR assays were performed using a Biacore T200 system using CM5 carboxymethyldextran sensor chips (Cytiva, USA). The chips were pre-coated with anti-His antibodies (Biacore His-Capture Kit, Cytiva, USA), enabling complete regeneration of His_6_-tagged molecules from the sensor surface. Initially, the chips were equilibrated with HBS-EP buffer [10 mM HEPES (pH 7.4), 150 mM NaCl, 3 mM EDTA, 0.005% (v/v) surfactant P20] until the dextran matrix was fully swollen. Subsequently, the flow cells of each sensor chip were activated by injecting a 1:1 mixture of N-ethyl-N′-(3-dimethylaminopropyl)carbodiimide hydrochloride and N-hydroxysuccinimide, using the standard amine coupling protocol. All flow cells were loaded with anti-His_6_ antibody at a final concentration of 50 μg mL^-1^ in 10 mM sodium acetate (pH 4.5) for a contact time of 420 seconds, resulting in antibody surface densities of approximately 9000 to 12,000 response units (RU). Unoccupied binding sites were blocked by injection of 1 M ethanolamine/HCl (pH 8.0). Throughout each experiment, a reference curve was generated by injecting the same buffer used for protein dilution (0.1 M potassium phosphate buffer, pH 7.0). To eliminate bulk refractive index effects, experimental sensorgrams were corrected by subtraction of the reference curve. The His_6_-tagged pure proteins SpoVG (50 nM) and Kre (20 nM) were captured onto the chip in HBS-EP buffer at a flow rate of 10 μL min^-1^ for 60 seconds, yielding a final response of approximately 100 RU, respectively. RNA was injected over the chip using a multi-cycle kinetics protocol at a flow rate of 30 μL/min. Increasing concentrations (10 nM, 25 nM, 50 nM, 100 nM, 250 nM, 500 nM, 1,000 nM, 2,500 nM, and 5,000 nM) were sequentially injected, with intermediate regeneration steps, with a contact time of 180 seconds and a dissociation phase of 800 seconds. The 250 nM concentrations were injected as duplicate in the end of each experiment to check surface intactness throughout the run. Regeneration was achieved by injecting 10 mM glycine (pH 1.5) for 60 seconds at a flow rate of 30 μL min^-1^ across all flow cells, effectively removing both Kre and SpoVG protein and RNA from the chip surface. Additionally, blank cycle kinetics were recorded by injecting two 0 nM concentrations (= buffer) following protein capture, as well as a 0 nM concentration at the end of each measurement series. Each multi-cycle kinetics run was conducted 3-4 times at 25 °C. Sensorgrams were recorded using Biacore T200 Control Software 3.2 and analysed with Biacore T200 Evaluation Software 3.2. Flow cell 1 was used to record blank sensorgrams for subtraction of background bulk refractive index effects. Buffer controls from flow cells 2, 3, and 4 were subtracted from their respective sample sensorgrams to normalise for baseline drift. The resulting reference-subtracted sensorgrams were then normalised to a baseline of zero. Spikes observed at the beginning and end of injections were due to differences in flow path lengths between the flow cells on individual chips. R_max_ was calculated by measuring the maximum binding response for a 1:1 interaction and applying the formula 1 and binding stoichiometry (n) was calculated using the formular 2.

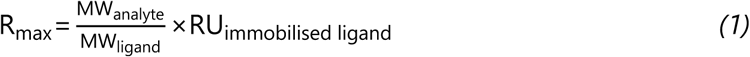

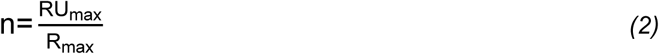

### Interaction Map Analysis

Interaction Map (IM) calculations were performed on the Ridgeview Diagnostic Server (Ridgeview Diagnostics, Uppsala, Sweden). For this purpose, the SPR sensorgrams were exported from the Biacore T200 Evaluation Software 3.2.1 as *.txt files and imported into TraceDrawer Software 1.10.1 (Ridgeview Instruments, Uppsala, Sweden). IM files were created using the IM tool within the software, generating files that were sent via e-mail to the server, where the IM calculations were performed (34). The resulting files were then evaluated for spots in the TraceDrawer 1.10.1 Software, and the IM spots were quantified.

### Mass Spectrometry

The wild type *Bacillus subtilis* DK1042, as well as *spoVG* and *kre* deletion mutants, were cultivated overnight in LB medium supplemented with the appropriate antibiotics. Overnight cultures were used to inoculate 50 mL DSM day cultures to an initial OD_600_ of 0.1, and incubated at 37 °C with shaking at 150 rpm. Samples were collected after 3, 4, 5, and 6 h of growth. Biological and technical triplicates were prepared. For protein extraction, 50 mL of culture were harvested by centrifugation at 10,000 rpm for 10 min at 4 °C. The supernatant was discarded, and cell pellets were resuspended in 20 mM Tris-HCl (pH 8.0). Cells were washed twice by centrifugation (10,000 rpm, 15 min) and resuspension in fresh Tris-HCl buffer. Pellets were subsequently resuspended in 2 mL lysis buffer (20 mM Tris-HCl, 5 mg mL^-1^ lysozyme, and 0.01 mg mL^-1^ DNase I) and incubated for 45 min at 4 °C. Cell disruption was performed by sonication (3 × 30 s) using a Sonopuls GM 70 (Bandelin, Berlin, Germany) at 100% amplitude and a cycle of 0.6. Following lysis, samples were centrifuged at 10,000 rpm, and the supernatant was transferred to a fresh tube and stored at -20 °C. For mass spectrometry (MS) analysis, proteins were digested in solution with either, LysC or trypsin, using RapiGest surfactant (Waters) to enhance digestion efficiency. Briefly, RapiGest was solved in 50 mM triethylammonium bicarbonate (TEAB, Sigma-Aldrich (St- Louis, MO, USA)) and added to 100 µg protein sample obtain a final concentration of 0.2% (w/v) RapiGest. Proteins were reduced with 500 mM Tris-(2-carboxyethyl)-phosphin (TCEP, Carl Roth GmbH and Co. KG (Karlsruhe, Germany)) at 80 °C for 20 min and alkylated with 500 mM iodoacetamide (IAA, Sigma-Aldrich (St- Louis, MO, USA)) for 15 min in the dark at room temperature. Afterwards samples were split into two aliquots of 50 µg protein content and 250 ng trypsin (Promega Corporation, Madison, Wisconsin, USA) was added to one aliquot, whereas 500 ng LysC (Promega Corporation, Madison, Wisconsin, USA) was added to the other. Both aliquots were digested for 5 hours at 37 °C. Upon completion of the digest, samples were acidified by the addition of 6 M HCl to obtain a pH <2. RapiGest was hydrolysed for 30 min at 4 °C and removed by two consecutive centrifugation steps. Obtained crude peptide mixture was purified using Pierce C18 tips (Thermo Fisher Scientific, Waltham, MA, USA) according to the manufacturer’s protocol and dried peptides were stored until measurement for which they were dissolved in 20 µL 0.1% (v/v) acetic acid.

Prior to liquid chromatography-MS (LC-MS) analysis, a transition list of EpeX, EpeE, SpoVG, and Kre was generated and optimised in order to keep the peptides with the highest response factors. Correctness of transitions was additionally verified by a supplemental LC-MS/MS run comprising product ion scans of target peptides, by checking their retention times, and by overlaying transition profiles. Peptides were separated by reversed phase column chromatography using an EASY nLC II (Thermo Fisher Scientific, Waltham, MA, USA) with self-packed columns (outer diamter 360 μm, inner diameter 100 μm, length 20 cm) filled with 3 µm diameter C18 particles (Dr. Maisch, Ammerbuch-Entringen, Germany). Following loading/ desalting in 0.1% acetic acid in water, the peptides were separated by applying a binary non-linear gradient from 1-99% acetonitrile in 0.1% acetic acid over 80 min. The LC was coupled online to a triple quadrupole (Q1-3) mass spectrometer (TSQ Vantage, Thermo Fisher Scientific, Waltham, MA, USA) operated in nano-electrospray mode. For ionisation 2400 V of spray voltage and 240 °C capillary temperature was used. The selectivity for both Q1 and Q3 were set to 0.7 Da (full width at half maximum (FWHM)). The collision gas pressure of Q2 was set at 1.2 mTorr. TSQ Vantage was operated in selected reaction monitoring (SRM) mode applying the transition list provided in Table S4. Raw files were processed using Skyline (version 23.1.0.455) (35). Normalised peak areas for each peptide were exported from the software. A signal to noise (S/N) ratio was calculated by dividing the peak height by the background value (both exported from Skyline) for each peptide in each sample. Peptide peak areas were weighted based on their (S/N) ratios before being averaged. Information of at least 2 unique peptides were combined to yield the final protein abundance.

## RESULTS

### EPE Production is Tightly Linked to Spo0A-Activating Phosphorelay

Due to the tight integration of differentiation pathways outlined above, we reasoned that the regulatory network governing EPE production is likely influenced by multiple cellular processes. We therefore performed a comprehensive screen of a large panel of single-gene deletion mutants of stationary phase functions involved in quorum sensing, competence and the Spo0A-controlling phosphorelay, using the highly sensitive P*_liaI_*-*lux* reporter, to systematically uncover regulatory interdependencies and identify regulatory circuits and developmental pathways that modulate EPE production.

Among the tested gene deletions, several mutants associated with the activation of Spo0A led to altered EPE production, as inferred by the induction of the EPE-specific P*_liaI_*-*lux* reporter, suggesting a functional link between the phosphorelay and toxin production (Figure 3). As controls to calibrate our screen, we included *spo0A* and *abrB* deletions into our screen: Deletion of *spo0A* led to an approx. 100-fold reduction in EPE-mediated stress, while absence of *abrB* increased EPE stress levels about two-fold (Figure 3b), in agreement with the Spo0A-dependent de-repression of AbrB in controlling *epeXEPAB* expression, as described previously (6).Deletion of *spo0F*, which is essential for initiating the phosphorelay that ultimately phosphorylates Spo0A (Figure 3a), resulted in a 100-fold reduction of EPE-mediated stress, comparable to the behaviour of a *spo0A* deletion (Figure 3b). Remarkably, P*_liaI_*-*lux* activity in both *spo0F* and *spo0A* mutants was even lower than that observed in the *epeX* deletion strain, suggesting that EPE-mediated stress is entirely absent in these backgrounds. However, sporulation-deficient mutants, like Δ*spo0A*, often suffer from severe growth defects, which can affect the relative luminescence units (RLU). These physiological effects may partially account for the observed differences (Figure S2).

**Figure 3:**
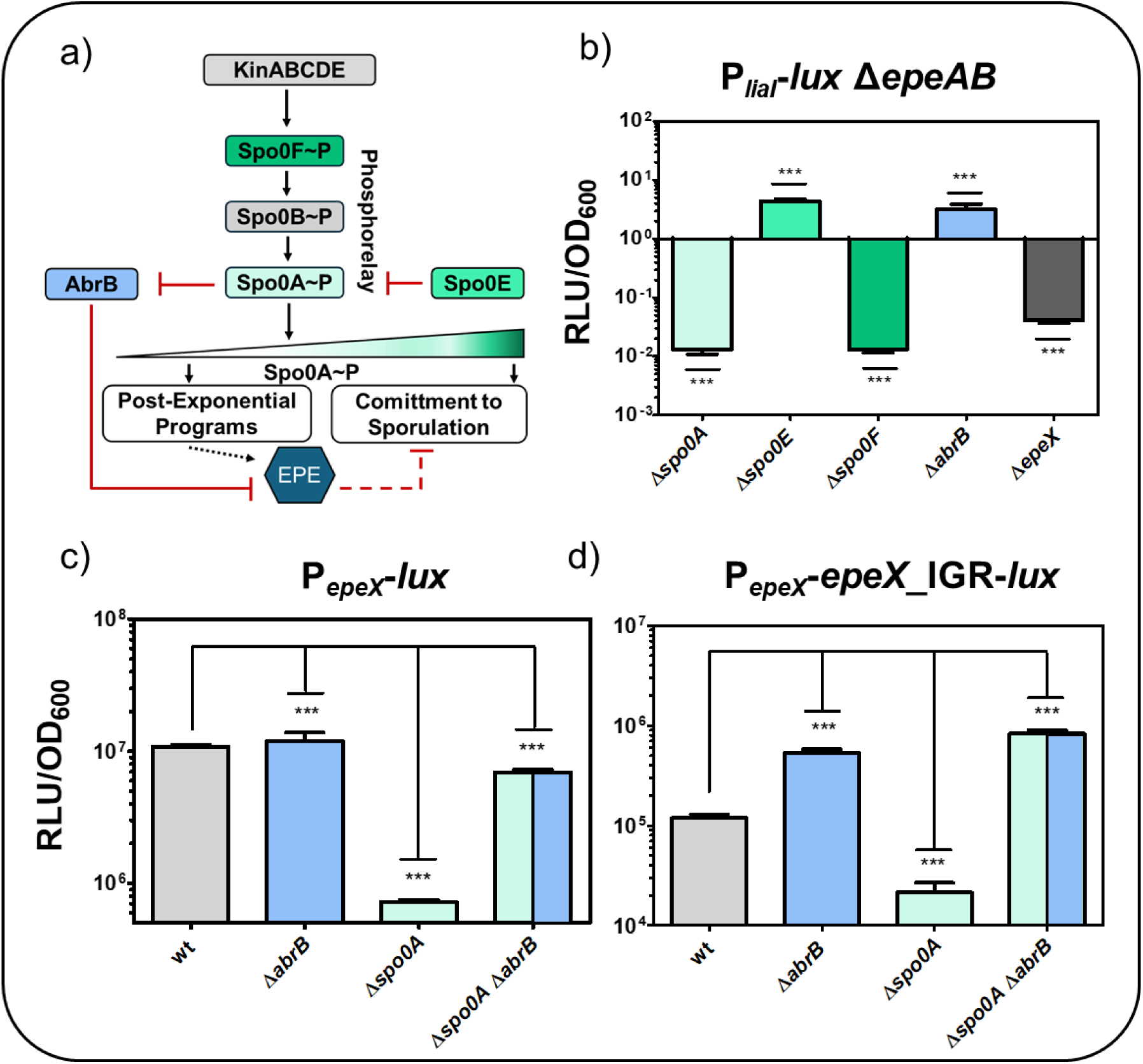
EPE production is under the direct control of Spo0A activation. **a)** Schematic overview of Spo0A activation, impacting *epe* expression. In response to environmental stress like nutrient starvation, the sensor kinases KinABCDE activate the phosphorelay of Spo0F and Spo0B, ultimately resulting in phosphorylation of Spo0A, the master regulator of sporulation. The phosphorylated state of Spo0A is under control of Spo0E. While Spo0A inhibits the transition state regulator AbrB, AbrB itself acts as transcriptional repressor of *epe* expression. Hence, activation of Spo0A (Spo0A∼P) leads to the production of EPE, causing cell lysis and subsequently release of nutrients, from which the toxin producer can fees. Thus, EPE production can indirectly prolong or even inhibit sporulation. However, prolonged nutrient starvation and concomitant accumulation of Spo0A∼P leads to commitment to sporulation. **b)** Effect of Spo0A-activating gene deletions on EPE-mediated stress response, depicted as fold change of maximum activities extracted from time-resolved data normalised to the reporter background strain P*_liaI_*-*lux epeAB*::*spec*. Deletion of *epeX* shows the background stress response, independently of EPE production. **c-d)** Maximal RLU/OD_600_ values over time were shown, demonstrating the impact of AbrB and Spo0A on **c)** *epeX* expression and **d)** *epeXEP* expression. **b-d)** Statistical significance was assessed using a one-way ANOVA followed by Dunnett’s post-hoc test comparing each mutant to the wild type. Significance is indicated as follows: ns = not significant; * = p<0.05; ** = p<0.01; *** = p<0.001). For simplification, gene deletions are indicated with the delta symbol (Δ) throughout the figure, although this does not necessarily imply clean deletion strains. The full genotypes and respective resistance cassettes are provided in Table S1.

Thus, the dramatically reduced EPE production observed in *spo0F* and *spo0A* mutants is caused by the absence of Spo0A∼P-mediated repression of *abrB*. Under such conditions, AbrB remains active, maintaining transcriptional repression of P*_epeX_*, and hence preventing EPE biosynthesis. Consequently, the EPE-mediated cell envelope stress response, as quantified by the P*_liaI_-lux*-dependent luminescence is not triggered in these mutants, as P*_epeX_* is never relieved from AbrB repression.

Deletion of *spo0E*, encoding the phosphatase responsible for dephosphorylating Spo0A∼P, resulted in a two-fold increase in EPE-mediated stress comparable to Δ*abrB*. This is caused by reduced dephosphorylation of Spo0A∼P, leading to enhanced repression of AbrB and therefore increased EPE production (Figure 3b).

In addition to the transcriptional P*_epeX_*-*lux* reporter, which allowed monitoring *epeX* expression, we also generated a translational P*_epeX_*-*epeX*-IGR-*lux* fusion, which includes *epeX*, the intergenic region (IGR) between *epeX* and *epeE*, as well as the native ribosome binding site of *epeE* to control *lux* expression. This construct served to assess the translation of *epeE* and *epeP* (Figure S1) (13). Consistent with the role of AbrB in repressing P*_epeX_* (6), deletion of *abrB* in both P*_epeX_*-*lux* and P*_epeX_*_*epeX*_IGR-*lux* reporter strains resulted in a one- to four-fold increase in luminescence, confirming its function as a transcriptional repressor of the *epeXEP* operon (Figure 3b & c). In contrast, deletion of *spo0A* led to a nine- to thirteen-fold reduction in reporter activity (Figure 3c & d). This strong repression was largely reversed by simultaneous deletion of *abrB*, supporting the conclusion that Spo0A indirectly activates P*_epeX_* by repressing *abrB* expression as previously demonstrated (6).

### Competence Development Regulates EPE production

AbrB acts as a global transition state repressor that controls the expression of hundreds of genes (36). In addition to its role in repressing the *epe* locus, AbrB also inhibits *comK* expression, and therefore activation of the master regulator of competence, ComK (37). Consequently, low levels of Spo0A∼P not only trigger *epe* expression but also relieve AbrB-mediated repression of *comK*, thereby promoting competence development. Competence is a transient physiological state that enables uptake of extracellular DNA from the environment. It is adopted by a small subpopulation of cells under conditions of nutrient limitation and high cell density (Figure 2 (II)) (38). Beyond promoting genetic diversity and DNA repair, competence can also contribute to nutrient acquisition through uptake and degradation of extracellular DNA, which can serve as a source of carbon-, nitrogen-, and phosphate-containing compounds (39). Since EPE production ultimately leads to lysis of sibling cells and the release of valuable nutrients, which could be taken up by the competent cells, we next examined if competence-associated genes similarly contribute to EPE regulation (Figure 4).

**Figure 4:**
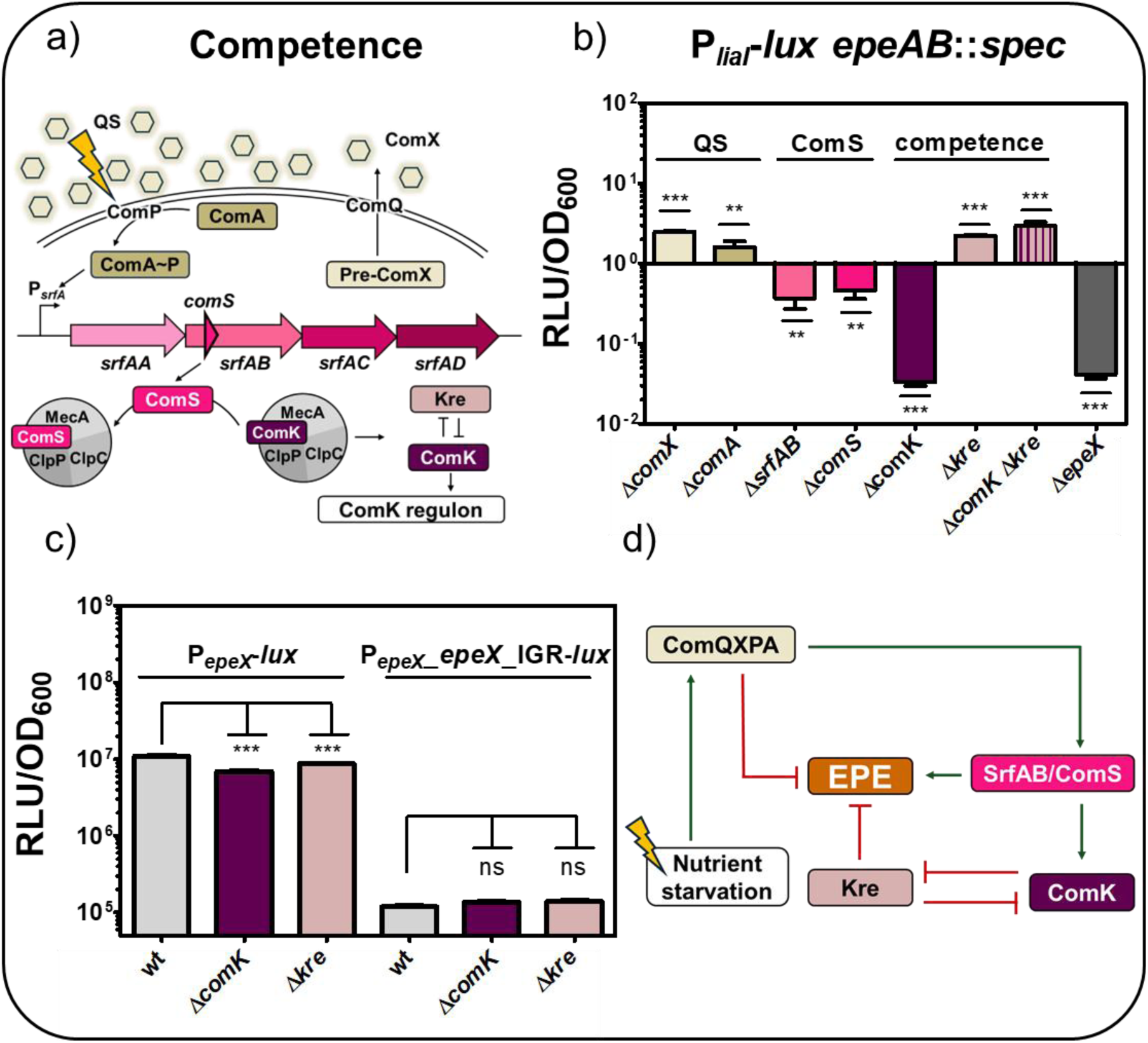
Effect of competence associated genes on EPE production and stress. **a)** Schematic representation of the competence pathway, involving quorum sensing (QS) dependent phosphorylation of ComA, allowing ComA to bind to P*_srfA_*. Thus, the *srf* operon is expressed, leading to production of surfactin and the small competence protein ComS, which prevents ComK for degradation due to higher binding affinity to the MecA-ClpC-ClpP proteolysis complex. Accumulation of ComK initiates expression of the ComK regulon and subsequently competence. Further, a bidirectional feedback loop between Kre and ComK is known. **b)** Effect of competence genes on EPE-induced stress response is shown as fold change of maximum RLU/OD_600_ values in comparison to the background P*_liaI_*-*lux epeAB*::*spec* reporter strain. **c)** Impact of ComK and Kre on *epeX* expression (left) and *epeXEP* expression (right) in comparison to the wild type (wt). Maximum RLU values normalised to the corresponding OD_600_ were shown. **b-c)** Statistical significance was assessed using a one-way ANOVA followed by Dunnett’s post-hoc test comparing each mutant to the wild type. Significance is indicated as follows: ns = not significant; * = p<0.05; ** = p<0.01; *** = p<0.001). For simplification, gene deletions are indicated with the delta symbol (Δ) throughout the figure, although this does not necessarily imply clean deletion strains. The full genotypes and respective resistance cassettes are provided in Table S1. **d)** Regulatory link between competence genes and EPE production derived from this analysis.

Competence development depends on quorum sensing, specifically the ComQXPA signalling pathway (Figure 4a). The peptide pheromone ComX is post-translationally modified by the prenyltransferase ComQ and sensed by the two-component system ComPA, comprised of the membrane-anchored sensor kinase ComP and the response regulator ComA (Figure 4a) (40).

As ComX accumulates extracellularly, ComP is activated and phosphorylates ComA (ComA∼P) (Figure 4a) (41). ComA∼P binds to the P*_srfAA_* promoter (42), leading to transcription of the *srfAABCD* (*srf*) operon (Figure 4a) (43,44). Embedded within this operon is the gene *comS*, a distinct open reading frame that overlaps with *srfAB* and encodes the small competence protein ComS. ComS prevents the proteolytic degradation of ComK by competitive binding to the MecA/ClpCP protease complex (Figure 4a). ComK expression is further auto-regulated by ComK itself and refined by Kre, which generates a bi-directional negative feedback loop with ComK (Figure 4a) (45,46). Once ComK reaches a critical threshold concentration, it not only activates its own expression but also induces its regulon, leading to expression of late competence genes essential for DNA uptake (46).

The influence of competence-associated genes on EPE production was assessed by measuring P*_liaI_*-*lux* activity as a measure of EPE production. Single gene deletion of *comX* and *comA*, which are essential for quorum sensing-mediated control of competence and surfactin production, resulted in a to two-fold increase in P*_liaI_* activity (Figure 4b). Deletion of *comS* alone or *srfAB*, which also removes the overlapping *comS* gene, led to an approximately seven-fold reduction in EPE-mediated stress. Remarkably, deletion of *comK*, the master regulator of late competence genes, resulted in a complete loss of EPE-induced stress, comparable to that observed in an *epeX* mutant (Figure 4b). These results indicate that ComK is essential for EPE production and highlight a novel regulatory interdependency between competence regulation and EPE production in *B. subtilis*.

While a *comK* deletion abolished EPE production entirely, the *epe* locus is not a member of the well-established ComK regulon (47). This suggested an indirect regulatory mechanism, by which ComK exerted its regulatory role on EPE. A potential candidate was Kre, which forms a bidirectional negative feedback loop with ComK (45). Initially described as a modulator of competence development, Kre has been proposed to regulate ComK by affecting the stability of its mRNA (45). Findings in *Bacillus anthracis* on the homologue KrrA show its function as an RNA-binding protein involved in post-transcriptional regulation (48).

We therefore determined the impact of Kre on the EPE-mediated stress response. While a two-fold increase of P*_liaI_* activity could be observed in a *kre* mutant, a double gene deletion of *comK* and *kre* fully restored EPE-dependent cell envelope stress response to the level of the *kre* mutant (Figure 4b). This suggests that the observed ComK effect is mediated indirectly through Kre, based on the bidirectional repression of ComK and Kre (Figure 4d).

Next, we analysed the impact of *comK* and *kre* deletions on both the transcriptional P*_epeX_*-*lux* and the translational P*_epeX_*_*epeX*_IGR-*lux* reporter (13) (Figure 4c). Despite the clear differences on EPE-mediated stress observed in the P*_liaI_-lux* reporter strains (Figure 4b), neither *comK* nor *kre* deletion affected *epeXEP* expression (Figure 4c). This is in contrast to the strong transcriptional repression of P*_epeX_* observed in sporulation-deficient mutants such as Δ*spo0A* (Figure 3b), *epeXEP* expression maintained wild type levels in *comK* and *kre* mutants (Figure 4c). These findings suggest that ComS-mediated ComK accumulation is essential for EPE production, presumably through its repression of Kre. The absence of any direct effect of Kre on *epeXEP* expression implies that Kre acts post-transcriptionally, potentially by altering mRNA stability or interfering with transcriptional termination (Figure 4c & d).

### The ComK-Repressor Kre Binds to the Intrinsic Structured IGR*_epeXE_*

We therefore investigated the potential post-transcriptional activity of Kre and its interaction with the *epeX* RNA to elucidate how Kre controls EPE production. Specifically, the RNA-binding properties of Kre and potential molecular targets at the *epe* locus were analysed.

Kre, an approximately 18 kDa protein, was heterologously overproduced in *E. coli* as a C-terminal histidine-tagged allele and purified by immobilised metal affinity chromatography (IMAC), followed by size exclusion chromatography. In SDS-PAGE, Kre-His_6_ showed a strong band of the expected molecular weight of 17.75 kDa (Figure S5a). While SDS-PAGE was performed under denaturation conditions, resulting in the monomeric state of a protein, attempts to characterise potential oligomeric states of Kre by native PAGE were hindered by its high isoelectric point (pI) of 9.8 (predicted by ProtParam), which is caused by the high number of positively charged residues, particularly 21 lysine residues, distributed throughout the protein. Consequently, it remains unclear whether Kre functions as a monomer, dimer, or higher-order oligomer in solution. Structural predictions using AlphaFold suggest that Kre comprises 4 *α*-helices, including a potential helix-turn-helix motif typically associated with nucleic acid binding (49). This predicted architecture is in agreement with its proposed function as an RNA-binding protein (Figure S5b) (48).

Since EMSA assays could also not be performed due to the high pI value of Kre, we employed surface plasmon resonance (SPR) spectroscopy to investigate the RNA-binding affinity of Kre to the *epeX* transcript. First, purified His_6_-tagged Kre protein was captured on the sensor chip that was previously immobilised with anti-His antibodies. Then, increasing concentrations, ranging from 10 nM to 5000 nM, of *epeX* RNA were injected over the chip surface. A panel of truncated *epeX* RNA fragments and the full-length *epeX* RNA were synthesised by *in vitro* transcription and applied in binding assays to explore potential binding sites across the *epeX* transcript (Figure 5a). The full-length transcript, designated *epeX* 1-11, spans from the transcriptional start site of *epeX* to the start codon of *epeE* (232 bp). Three partial fragments (*epeX* 1-3, *epeX* 5-7, and *epeX* 9-11 (80 bp each)) and a set of eight shorter 40 bp fragments (*epeX* 1 - *epeX* 11) were also generated to narrow down the binding region (Figure 5a). These fragments were designed with 20 bp overlaps between adjacent segments to ensure complete coverage of the transcript and minimise the risk of missing potential binding sites (Figure 5a).

**Figure 5:**
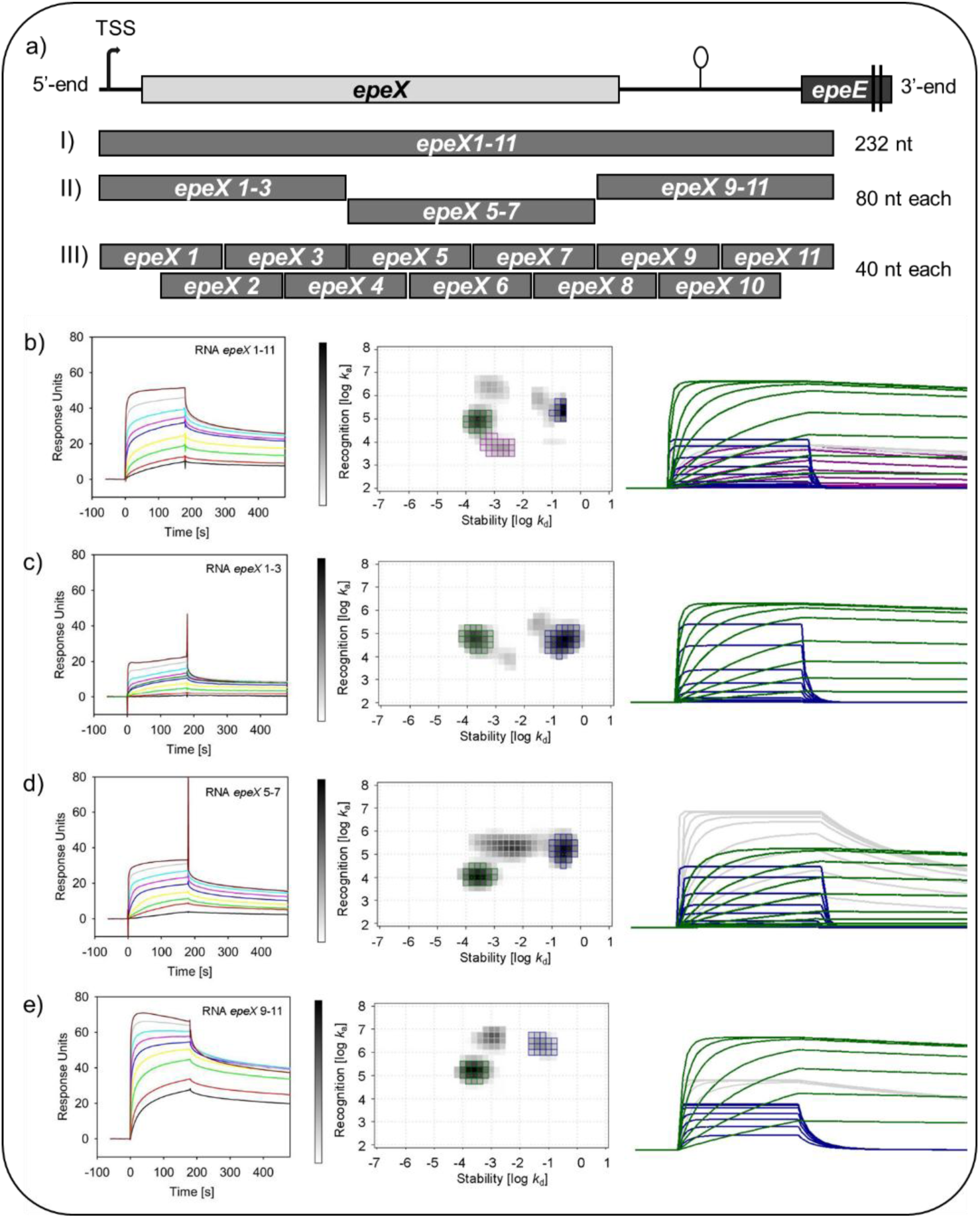
SPR analysis of Kre binding to full-length and truncated *epeX* RNA transcripts. **a)** Schematic overview of truncated *epeX* RNA fragments. **I**) The full-length RNA construct (*epeX* 1–11) spans from the +1 transcriptional start site of *epeX* to the start codon of *epeE* (232 nt). **II**) To identify specific binding regions, three mid-sized fragments (∼80 nt each) were generated: *epeX* 1–3, *epeX* 5–7, and *epeX* 9–11. **III**) For higher-resolution mapping, a tiling strategy was applied using eleven 40 bp fragments (*epeX* 1 through *epe*X 11), each overlapping its neighbouring fragment by 20 bp. **b-e)** Sensorgrams (left panels) depict response units of Kre to increasing concentrations of *epeX* RNA fragments (black: 10 nM; red: 25 nM; light green: 50 nM; yellow: 100 nM; blue: 250 nM; turquoise: 500 nM; light blue: 1000 nM; grey: 2500 nM; brown: 5000 nM). Interaction Maps (central panels) show the distribution of association and dissociation rate constants as stability (dissociation rate; log *k*_d_) and recognition (association rate, log *k*_a_). The corresponding calculated sensorgrams from the IM peaks (right panels) are presented on the right, the colours of the curved correspond to the respective peaks from the IM analyses. The data were quantified and the respective overall affinities (K_D_) calculated from the respective association (*k_a_*) and dissociation rates (*k_d_*) are shown, as well as the peak weights (PW) showing the overall contribution of the respective interaction towards the total sensorgrams given in (%). (see Table 1 for details). Since bulk binding are not included in the calculations, 100% were not reached in total. The full-length *epeX* RNA **b)** *epeX* 1-11, as well as truncated versions of the *epeX* RNA **c)** *epeX* 1-3, **d)** *epeX* 5-7, and **e)** *epeX* 9-11, were applied for SPR as well as IM analysis.

**Table 1:** Kinetic values derived from IM analysis for Kre binding to the full-length transcript *epeX* and truncated versions of *epeX* RNA. The table presents distinct kinetic populations along with their corresponding peak weights (PW), overall affinity (K_D_), dissociation rates (*k*_d_), and association rates (*k*_a_). Note that due to exclusion of nonspecific and bulk binding events for the analysis, the total population does not reach 100%.

| RNA fragment | Binding population | Peak weights [%] | $K_D$ | $k_d [\text{s}^{-1}]$ | $k_a [\text{M}^{-1} \text{s}^{-1}]$ |
| --- | --- | --- | --- | --- | --- |
| *epeX* 1-11 | Green | 32 | 4 nM | $2.8 \times 10^{-4}$ | $7.5 \times 10^4$ |
| | Blue | 17 | 0.9 $\mu\text{M}$ | $2.0 \times 10^{-1}$ | $2.3 \times 10^5$ |
| | Grey | 13 | 0.4 nM | $7.5 \times 10^{-4}$ | $2.2 \times 10^6$ |
| | Purple | 14 | 0.2 $\mu\text{M}$ | $1.3 \times 10^{-3}$ | $6.3 \times 10^3$ |
| *epeX* 1-3 | Green | 30 | 4 nM | $2.0 \times 10^{-4}$ | $5.7 \times 10^4$ |
| | Blue | 44 | 4 $\mu\text{M}$ | $2.0 \times 10^{-1}$ | $5.0 \times 10^4$ |
| *epeX* 5-7 | Green | 24 | 24 nM | $2.8 \times 10^{-4}$ | $1.2 \times 10^4$ |
| | Blue | 25 | 2 $\mu\text{M}$ | $2.5 \times 10^{-1}$ | $1.6 \times 10^5$ |
| | Grey | 35 | 13 nM | $3.1 \times 10^{-3}$ | $2.4 \times 10^5$ |
| *epeX* 9-11 | Green | 41 | 2 nM | $2.5 \times 10^{-4}$ | $1.6 \times 10^5$ |
| | Blue | 18 | 26 nM | $5.6 \times 10^{-2}$ | $2.2 \times 10^6$ |
| | Grey | 26 | 0.3 nM | $1.1 \times 10^{-3}$ | $4.5 \times 10^6$ |

Binding kinetics of Kre to distinct *epeX* RNA transcripts were evaluated by SPR analysis. Since the resulting sensorgrams (Figure 5b, left panels) did not follow a simple 1:1 binding stoichiometry, Interaction Map (IM) analyses were performed, in which the interactions are split into the respective binding events that contributes to the overall sensorgrams. Heatmaps of association (*k*_a_) and dissociation (*k*_d_) rates are displayed, in which peaks correspond to individual 1:1 binding event (Figure 5b, middle panel). The kinetics, affinities and prevalence of each binding mode is presented, together with its corresponding binding curve in Figure 5 and Table 1.

Initial experiments utilised the full-length *epeX* transcript (*epeX* 1-11) to assess binding potential (Figure 5a). The titration reached ∼50 response units (RU), indicating specific interactions and revealing a complex binding behaviour characterised by four distinct kinetic populations (Figure 5b).

The IM analysis suggested a primary Kre binding population (green), comprising 32% of the total binding response, exhibiting high binding affinity with an overall affinity (K_D_) of 4 nM, based on rapid association rate (*k*_a_=7.5 × 10^4^ M^-1^ s^-1^), and slow dissociation rate (*k*_d_=2.8 × 10^-4^ s^-1^). A smaller Kre binding population (grey; 13%) displayed an even faster association (*k*_a_=2.2 × 10^6^ M^-1^ s^-1^) and a relatively slow dissociation rate (*k*_d_=7.5 × 10^-4^ s^-1^), resulting in a lower K_D_ of 0.4 nM. Additional Kre populations were identified with reduced binding affinity [K_D_=0.9 µM (blue) and K_D_=0.2 µM (purple)], indicating heterogeneity and suggesting a multivalent interaction mode (Figure 5b & Table 1).

Subsequently, three truncated fragments of *epeX* RNA (*epeX* 1-3, *epeX* 5-7, and *epeX* 9-11) were tested to localise the interaction (Figure 5a). The 5’-fragment of the *epeX* RNA, (*epeX* 1-3) yielded two kinetic populations and reduced response units of ∼20 RU, representing an approximately 2.5-fold decrease relative to full-length *epeX* RNA. One population (green; 30%) exhibited high stability (K_D_=4 nM, *k*_a_=5.7 × 10^4^ M^-1^ s^-1^, and *k*_d_=2.0 × 10^-4^ s^-1^), though overall response was lower than for the full-length transcript. The second population (blue; 44%) was less stable with K_D_=4µM (Figure 5c & Table 1).

The central region of the *epeX* RNA (*epeX* 5-7), resulted in an intermediate RU of ∼30, and revealed three kinetic populations (Figure 5a). The most stable population (green; 24%) had a K_D_=24 nM, *k*_a_=1.2 × 10^4^ M^-1^ s^-1^, and *k*_d_=2.8 × 10^-4^ s^-1^. The remaining populations (grey, and blue) showed faster association, and higher dissociation rates (*k*_a_=2.4 × 10^5^ M^-1^ s^-1^, *k*_d_=3.1 × 10^-3^ s^-1^ and *k*_a_=1.6 × 10^5^ M^-1^ s^-1^; *k*_d_=2.5 × 10^-1^ s^-1^, respectively), consistent with weaker and more transient interactions (Figure 5d & Table 1).

The 3’-fragment of the *epeX* RNA (*epeX* 9-11), encompassing the IGR*_epeXE_* (Figure 5a), led to the highest binding response (∼70 RU), and showed strong affinity across three populations. Two populations (green, and grey) displayed particularly high stability with K_D_ values of 2 nM (*k*_a_=1.6 × 10^5^ M^-1^ s^-1^, and *k*_d_=2.5 × 10^-4^ s^-1^), and 0.3 nM (*k*_a_=4.5 × 10^6^ M^-1^ s^-1^, and *k*_d_=1.1 × 10^-3^ s^-1^), respectively (Figure 5e & Table 1).

Additionally, the set of 11 short fragments of the *epeX* transcript (*epeX* 1 - *epeX* 11) were analysed to distinguish interactions sites (Figure 5a). No detectable binding was observed for the short 40 bp fragments spanning *epeX* 1 to 8. In contrast, measurable binding occurred in fragments overlapping the terminator structure within the IGR*_epeXE_*. The strongest binding among these was observed for *epeX* 10, yielding a K_D_ of 81 nM, and comprising 57% of the overall response (Figure S6b). This interaction was characterised by a particular slow dissociation rate (*k*_d_=7.6 × 10^4^ s^-1^), despite a relatively slow association rate (*k*_a_=9.4 × 10^2^ M^-1^ s^-1^), suggesting a highly stable complex (Figure S6b). Notably *epeX* 10 contains the terminator hairpin, implicating this structure as a key binding site of Kre (Figure 5a). The fragments *epeX* 9 and 11 also showed binding with reduced affinity (K_D_=4 µM, and 0.9 µM, respectively), consistent with partial overlap with the terminator (Figure S6a & c).

Taken together, the SPR data revealed that Kre binds the *epeX* RNA with multiple kinetic populations, consistent with a complex, multivalent interaction landscape. The highest-affinity binding was consistently observed at the 3’-end of the transcript (*epeX* 9-11), encompassing the IGR*_epeXE_*. The Kre-*epeX* complex may be further stabilised by base-pairing interactions with other parts of the *epeX* RNA.

These findings support a model in which Kre preferentially engages the terminator hairpin structure, and presumably contributes to post-transcriptional regulation of EPE production. The modular and differential binding affinities across the RNA suggest as multivalent recognition mechanism, in which Kre interacts with multiple regions of the RNA, potentially by both sequence elements and structural motifs like helices and bulges. The consistent association of slow dissociation rates and high binding affinity with the IGR*_epeXE_*, indicates this as a key regulatory target for the RNA-binding protein Kre.

### The RNA-Binding Protein SpoVG Binds to the *epeX* RNA

While Kre binds to the terminator structures within the IGR*_epeXE_* and potentially promotes transcription termination and subsequent RNA decay, the exceptional stability of the *epeX* mRNA, with a half-life exceeding 15 min (50), indicates that additional factors at the RNA level may contribute to its post transcriptional control, particularly at the level of RNA stability. In *B. subtilis*, as in most bacteria, mRNAs are generally short-lived, with an average half-life of approximately 2-5 min (51). In contrast, the *epeX* mRNA as one of the most stable transcripts in the cell (50,52).

We wondered what might be the cause of the exceptional stability of the *epeX* transcript. One explanation could be that global RNA binding proteins bind and thereby shield specific transcripts from RNA decay pathways (53). One possible candidate is SpoVG. The *spoVG* gene was first identified in *B. subtilis* as playing a role in sporulation (54). However, it has been shown to be widely conserved in low-GC Gram-positive bacteria and in Spirochetes such as *Borrelia burgdorferi* (55). SpoVG has been demonstrated to bind both DNA and RNA *in vitro*, so far. In *Listeria monocytogenes* and *Staphylococcus aureus*, it contributes to regulation of diverse processes such as biofilm formation, swarming motility and virulence, which poses the likelihood that it is a global regulator (56,57). These observations raise the question whether SpoVG is important in *B. subtilis* beyond its role in sporulation, where the mechanism of regulation has yet to be defined. Therefore, we decided to test whether SpoVG is able to regulate the *epeX* transcript at the post-transcriptional level.

Recombinant SpoVG with a C-terminal His_6_-tag was expressed and purified using IMAC followed by size exclusion chromatography. SDS-PAGE revealed a distinct band, which corresponded to the monomeric form of approx. 11 kDa (Figure S7a). Subsequently, we assessed the oligomeric state of SpoVG by native PAGE analysis. The protein predominantly migrated as a band with an apparent molecular weight of >21 kDa, consistent with the formation of a homodimer under native conditions (Figure S7c). This result is in agreement with the released crystallographic data (Protein Data Bank (PDB): 2IA9), which revealed that SpoVG forms a stable symmetric dimer mediated by *β*-sheet interactions and hydrophobic contacts (58). Structurally, SpoVG forms a *α*/*β* complex, consisting of six antiparallel *β*-strands that form a twisted *β*-sheet core, along with a single C-terminal *α*-helix. The *α*-helix aligns along one edge of the *β*-sheet, possibly contributing to dimer stabilisation and the formation of an RNA-binding interface. The dimerization results in a broad, continuous surface with conserved, positively charged residues, supporting a potential role in nucleic acid binding (Figure S7b). Overall, the structural features of SpoVG are consistent with its proposed function as an RNA binding protein and post-transcriptional regulator.

The RNA-binding activity of SpoVG was initially evaluated by electrophoretic mobility shift assays (EMSA) using the full-length *epeX* transcript ranging from the +1 region of *epeX* till the 3’-end of the intergenic region (IGR) between *epeX* and *epeE* (232 bp) (Figure S7d). A constant amount of the *epeX* RNA (5 pmol) was incubated with increasing concentrations of SpoVG ranging from 15 till 640 pmol. EMSA revealed complete RNA shifting with no detectable free RNA, indicating strong and saturated binding. Strikingly, the RNA-protein complexes exhibited both upward and downward shifts in mobility (Figure S7d). The up-shifted bands likely correspond to conventional RNA-protein complexes, where mobility decreases proportionally with increasing protein stoichiometry and the number of occupied binding sites, consistent with high-affinity, saturable interactions (59). In contrast, the downward shifts, where complexes migrate faster than free RNA, may indicate that SpoVG binding induces structural rearrangements within the *epeX* RNA that reduce its effective hydrodynamic radius or alter its charge distribution. Because native gel mobility is determined not only by molecular weight but also by net charge, shape, and conformation, binding of the small protein SpoVG (10.75 kDa) to the 232 nt *epeX* RNA, could plausibly lead to increased mobility rather than the typically slower migration (60). Similar downshifts have been reported previously, supporting this interpretation (61).

The binding behaviour of SpoVG to distinct *epeX* RNA fragments was subsequently quantitatively analysed by SPR spectroscopy analysis. Purified His_6_-tagged SpoVG protein was captured on the sensor chip that was previously immobilised with anti-His antibodies. Increasing concentrations of the *epeX* RNA, ranging from 10 nM to 5000 nM, were injected over the chip surface. The sensorgrams were recorded similarly as described above and binding heterogeneity was further resolved by IM analysis, which decomposed the sensorgrams into distinct SpoVG binding populations based on their *k*_a_ and *k*_d_ values. This allows the overall sensorgram to be deconvoluted into its contributing binding modes (Figure 6 &Table 2).

**Figure 6:**
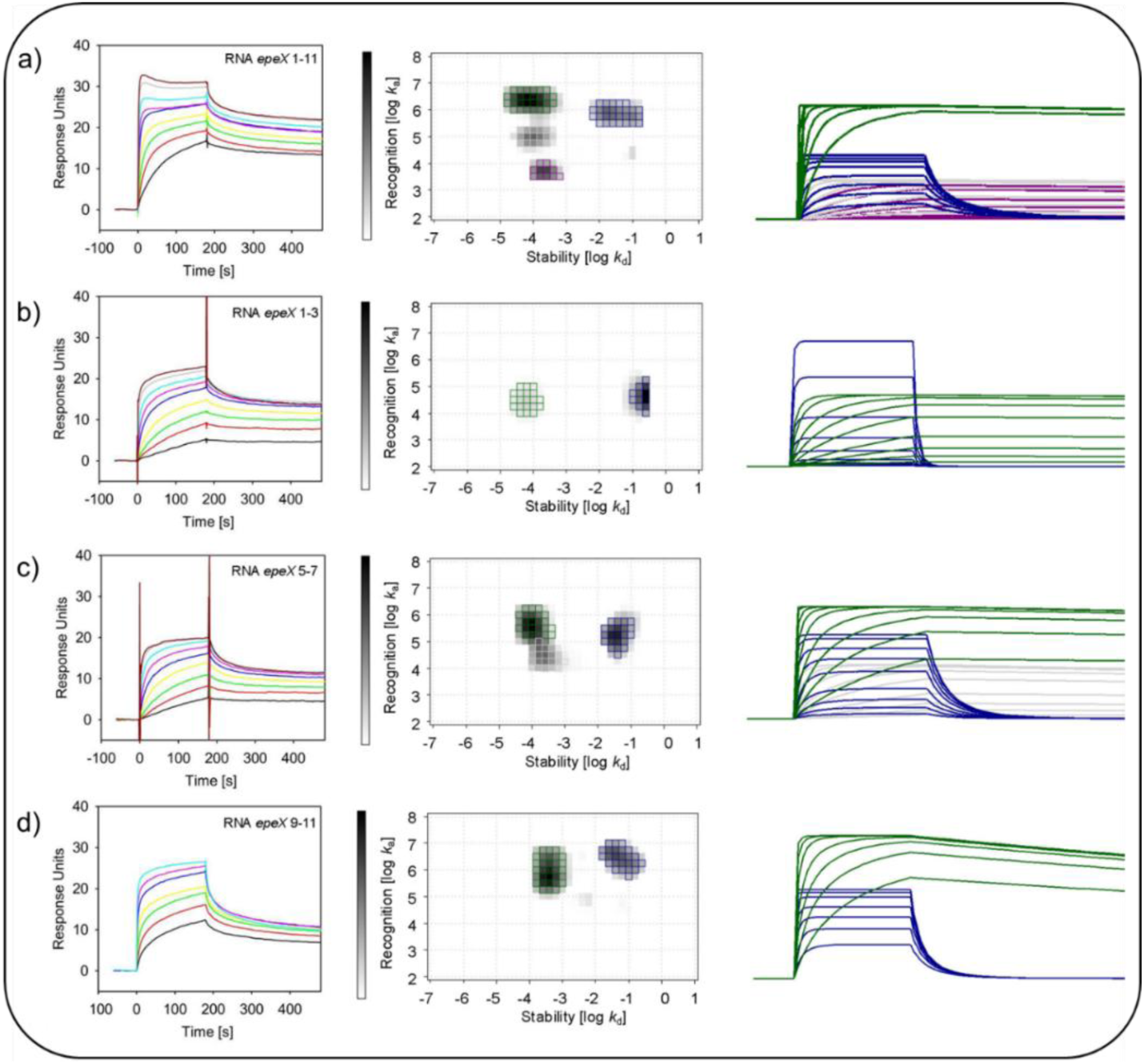
SPR analysis of SpoVG binding to full-length and truncated *epeX* RNA transcripts. Sensorgrams (left panels) depict response units of SpoVG to increasing concentrations of *epeX* RNA fragments (black: 10 nM; red: 25 nM; light green: 50 nM; yellow: 100 nM; blue: 250 nM; turquoise: 500 nM; light blue: 1000 nM; grey: 2500 nM; brown: 5000 nM). Interaction Maps (central panels) show the distribution of association and dissociation rate constants as stability (dissociation rate; log *k*_d_) and recognition (association rate, log *k*_a_). The corresponding calculated sensorgrams from the IM peaks (right panels) are presented on the right, the colours of the curved correspond to the respective peaks from the IM analyses. The data were quantified and the respective overall affinities (K_D_) calculated from the respective association (*k_a_*) and dissociation rates (*k_d_*) are shown, as well as the peak weights (PW) showing the overall contribution of the respective interaction towards the total sensorgrams given in (%). (see Table 2 for details). Since bulk binding are not included in the calculations, 100% were not reached in total. The full-length *epeX* RNA **a)** *epeX* 1-11, as well as truncated versions of the *epeX* RNA **b)** *epeX* 1-3, **c)** *epeX* 5-7, and **d)** *epeX* 9-11, were applied for SPR analysis.

**Table 2:** Kinetic values derived from IM analysis for SpoVG binding to the full-length transcript *epeX* and truncated versions of *epeX* RNA. The table presents distinct kinetic populations along with their corresponding peak weights (PW), overall affinity (K_D_), dissociation rates (*k*_d_), and association rates (*k*_a_). Note that due to exclusion of nonspecific and bulk binding events for the analysis, the total population does not reach 100%.

| RNA fragment | Binding population | Peak weights [%] | $K_D$ | $k_d [\text{s}^{-1}]$ | $k_a [\text{M}^{-1} \text{s}^{-1}]$ |
| --- | --- | --- | --- | --- | --- |
| *epeX* 1-11 | Green | 40 | 35 pM | $7.9 \times 10^{-5}$ | $2.3 \times 10^6$ |
| | Blue | 23 | 43 nM | $3.1 \times 10^{-2}$ | $7.2 \times 10^5$ |
| | Grey | 14 | 1 nM | $1.0 \times 10^{-4}$ | $9.9 \times 10^4$ |
| | Purple | 12 | 45 nM | $2.1 \times 10^{-4}$ | $4.7 \times 10^3$ |
| *epeX* 1-3 | Green | 19 | 2 nM | $5.9 \times 10^{-5}$ | $2.9 \times 10^4$ |
| | Blue | 63 | 5 $\mu\text{M}$ | $2.0 \times 10^{-1}$ | $4.3 \times 10^4$ |
| *epeX* 5-7 | Green | 38 | 0.3 nM | $1.1 \times 10^{-4}$ | $4.1 \times 10^5$ |
| | Blue | 30 | 0.2 $\mu\text{M}$ | $3.8 \times 10^{-2}$ | $1.7 \times 10^5$ |
| | Grey | 18 | 7 nM | $2.3 \times 10^{-4}$ | $3.3 \times 10^4$ |
| *epeX* 9-11 | Green | 54 | 0.4 nM | $3.9 \times 10^{-4}$ | $8.9 \times 10^5$ |
| | Blue | 35 | 20 nM | $5.6 \times 10^{-2}$ | $2.8 \times 10^6$ |

SpoVG exhibited strong binding to the full-length *epeX* RNA (*epeX* 1-11; Figure 5a), reaching approximately 35 response units (RU). IM analysis identified four kinetically distinct binding populations. The dominant population (green) accounted for 40% of the total response and displayed strong affinity (K_D_=35 pmol), with a high association rate (*k*_a_=2.3 × 10^6^ M^-1^ s^-1^) and an extremely slow dissociation rate (*k*_d_=7.9 × 10^-5^ s^-1^), indicating exceptional binding stability. Additional populations were detected with K_D_ values of 43 nM (blue; 23%), 1 nM (grey; 14%), and 45 nM (purple; 12%). These populations exhibited reduced association (*k*_a_=7.2 × 10^5^ M^-1^ s^-1^, *k*_a_=9.9 × 10^4^ M^-1^ s^-1^, and *k*_a_=4.7 × 10^3^ M^-1^ s^-1^) and increased dissociation rates (*k*_d_=3.1 × 10^-2^ s^-1^, *k*_d_=1.0 × 10^-4^ s^-1^, and *k*_d_=2.1 × 10^-4^ s^-1^), suggesting fewer stable interactions. The presence of multiple SpoVG binding populations suggests multivalent binding or non-specific interactions with structured regions of the RNA (Figure 6a & Table 2).

To localise physiological relevant interactions sites, truncated version of the *epeX* RNA were analysed. Binding of SpoVG to the 5’-fragment (*epeX* 1-3; Figure 5a) resulted in a simplified kinetic profile with two distinct populations. The dominant population (blue) represented 63% of the overall response and exhibited lower affinity (K_D_=5 µM), characterised by a moderate association rate (*k*_a_=4.3 × 10^4^ M^-1^ s^-1^), and rapid dissociation (*k*_d_=2.0 × 10^-1^ s^-1^). In contrast, a minor but kinetically stable population (green; 19%) displayed high-affinity binding (K_D_=2 nM), driven by a high association rate (*k*_a_=2.9 × 10^4^ M^-1^ s^-1^) and a particular slow off-rate (*k*_d_=5.9 × 10^-5^ s^-^1) (Figure 6b & Table 2). These kinetic parameters suggest the presence of a highly stable binding site within the 5’-region of the *epeX* RNA (Figure 6b & Table 2).

SpoVG binding to the central fragment *epeX* 5-7 (Figure 5a) resulted in an intermediate overall response and increased kinetic complexity, with three distinct populations identified. The most stable population (green; 38%) exhibited a K_D_ of 0.3 nM, with a fast association (*k*_a_=4.1 × 10^4^ M^-1^ s^-1^) and slow dissociation rate (*k*_d_=1.1 × 10^-4^ s^-1^). However, this interaction was less stable than that observed for the *epeX* 1-3 fragment. The remaining two populations included a moderate (blue; 30%), and a weaker interaction (grey; 18%) with K_D_= 0.2 µM, and K_D_=7 nM, respectively (Figure 6c & Table 2).

The 3’-fragment *epeX* 9-11 (Figure 5a), which includes the terminator structure within the IGR*_epeXE_*, exhibited two major binding populations. The dominant population (green; 54%) exhibited high affinity (K_D_=0.4 nM), resulting from a high association rate (*k*_a_=8.9 × 10^5^ M^-1^ s^-1^), and a relatively fast dissociation rate (*k*_d_=3.9 × 10^-4^ s^-1^). The second population (blue; 35%) displayed lower affinity (K_D_=20 nM), due to a very rapid association (*k*_a_=2.8 × 10^6^ M^-1^ s^-1^), and high dissociation rate (*k*_d_=5.6 × 10^-2^ s^-1^). Although, SpoVG associates rapidly with this RNA region, the instability of these binding events suggest they may arise from structural feature rather than specific regulatory interactions (Figure 6d & Table 2).

These binding profiles indicate that SpoVG is a strong RNA-binding protein, capable of interaction with multiple RNA regions across the *epeX* RNA. However, the high affinity across all fragments, including the 5s RNA as control (Figure S8), implies that not all binding events are biologically meaningful. Instead, some interactions may reflect general RNA binding affinity or structure-driven binding. Among all fragments tested, *epeX* 1-3 displayed the slowest dissociation rate, suggesting that this RNA region is most likely physiologically relevant. In contrast, binding to *epeX* 5-7, and *epeX* 9-11 occurred with fast association and reduced stability, which potentially reflect supportive interactions that stabilise SpoVG binding to the 5’-region and promotes overall *epeX* transcript stability. Additional analysis using shorter *epeX* truncations (*epeX* 1 to 11) confirmed SpoVG binding near the extreme 5’-end of the transcript, potentially enhancing transcript stability by blocking 5’-exonucleolytic degradation (Figure S8a). Collectively, the data indicate that although SpoVG can bind multiple regions of the *epeX* transcript, *epeX* 1-3 showed the strongest evidence for a biologically relevant, sequence-specific interaction.

### Interdependency of ComK, Kre, and SpoVG in EPE production

After establishing the direct binding of Kre and SpoVG to the *epeX* transcript, we next addressed the regulatory role of SpoVG in the context of the Kre-ComK switch. The effects of single and combinatorial deletions of *kre*, *comK*, and *spoVG* were assessed using the P*_liaI_*-*lux* reporter system as a readout for the EPE-mediated stress response to evaluate the physiological relevance of ComK, Kre and SpoVG in regulating EPE biosynthesis (13). For reasons of reference the data of the *comK, kre* as well as *comK/kre* double mutants, that were already described in detail above, are again reproduced in Figure 7a, demonstrating the strong effect of ComK on EPE production, which is exerted indirectly, via Kre (Figure 7a).

**Figure 7:**
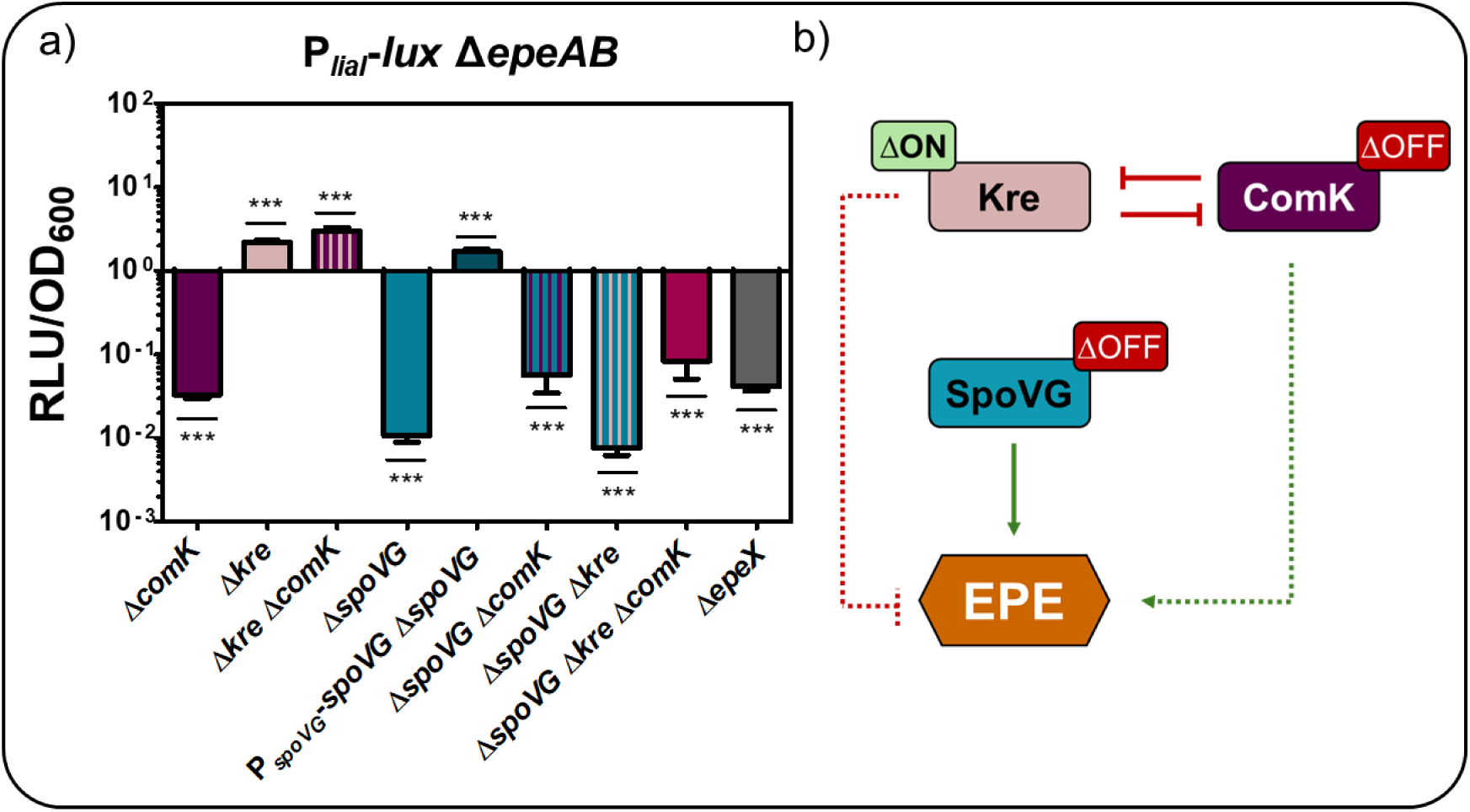
Effect of *comK*, *kre* and *spoVG* single and combinatorial mutants on EPE-dependent stress response. **a)** The impact of gene deletion strains in comparison to the *epeX* mutant on EPE stress response was shown as fold change of maximum RLU/OD_600_ values normalized to the reporter background strain P*_liaI_*-*lux epeAB*::*spec*. Statistical significance was assessed using a one-way ANOVA followed by Dunnett’s post-hoc test comparing each mutant to the wild type. Significance is indicated as follows: ns = not significant; * = p<0.05; ** = p<0.01; *** = p<0.001). For simplification, gene deletions are indicated with the delta symbol (Δ) throughout the figure, although this does not necessarily imply clean deletion strains. The full genotypes and respective resistance cassettes are provided in Table S1 **b)** Potential regulation scheme resolved from a.

This genetic relationship supports a regulatory model in which ComK downregulates Kre, thereby relieving post-transcriptional regulation at the *epeXEP* operon and enabling toxin synthesis (Figure 7b).

In contrast, deletion of *spoVG* led to a drastic reduction in EPE-dependent P*_liaI_* activity, which was reduced approximately 100-fold compared to the wild type reporter strain, demonstrating a critical importance of SpoVG in activating EPE production (Figure 7a). A complementation strain was constructed in which *spoVG*, driven by its native promoter, was integrated into the ectopic *amyE* locus (Figure 7a). This strain fully restored EPE production to wild type levels, confirming the direct regulatory role of SpoVG. The seemingly lower P*_liaI_* activity in the *spoVG* mutant compared to that observed in the *epeX* deletion strain is an artifact caused by the severe growth defects in the *spoVG* mutant, particularly during the transition to stationary phase (Figure S9), since the luminescence values are normalised by the optical density of the culture. A marked decline in optical density was detected around 5 hours after inoculation, followed by delayed and attenuated growth persisting into late stationary phase. These growth impairments are consistent with a loss of sporulation capacity and broader disruptions in regulatory processes. Partial restoration of growth in the complementation strain, which reached wild type levels approximately 10 hours after inoculation (Figure S9), further supported the direct involvement of SpoVG in maintaining physiological fitness.

Next, the impact of a *spoVG* deletion on P*_epeX_*-*lux*, P*_epeX_*_*epeX*-*lux*, and P*_epeX_*_*epeX*_IGR-*lux* was investigated to assess its influence on *epeXEP* expression (Figure S10). Across all constructs, an approximately two- to five-fold reduction in promoter activity was observed compared to the wild type. However, the pronounced growth defects associated with *spoVG* deletion likely contribute to reduced reporter activity, making it difficult to distinguish between direct regulatory effects and indirect consequences of impaired cellular physiology. Notably, even the P*_epeX_*-*lux*, which contained the *epeX* promoter only, showed reduced activity, despite the absence of downstream *epeX* sequence (Figure S10). Combined with the demonstrated binding of SpoVG to the *epeX* mRNA, this suggests that reduced reporter activity is potentially caused by growth-related effects rather than direct transcriptional regulation. These findings support a model in which SpoVG exerts regulatory control of *epeXEP* expression at the post-transcriptional level, presumably by stabilising the *epeX* transcript.

We further dissected the regulatory interdependencies of Kre, ComK, and SpoVG, by constructing combinatorial mutants. The *spoVG*/*kre* as well as *spoVG*/*comK* double mutants both showed drastically reduced EPE production, with levels comparable to the *spoVG* single mutant (Figure 7a). These results suggest that SpoVG is epistatic to both Kre and ComK, and that neither deletion of the repressor *kre* nor the upstream regulator *comK* can compensate for the loss of *spoVG*. This indicates that SpoVG is the primary activator in the regulatory cascade controlling EPE expression. The triple deletion mutant *spoVG*/*kre*/*comK* was unable to produce EPE, confirming that SpoVG is indispensable for EPE biosynthesis (Figure 7a). However, the double and triple mutants also showed severe physiological defects, including impaired growth due to inability of sporulation and development of natural competence (Figure S9).

Altogether, these data demonstrate that SpoVG is the central regulator required for EPE production, while Kre acts as a negative regulator and ComK as an indirect activator via repression of Kre. The inability of *kre* or *comK* deletion to restore EPE production in the absence of SpoVG highlights its dominant and essential role in this regulatory network (Figure 7b).

### Time Resolved Abundance of Proteins Impacting EPE production

To assess whether this regulation translates into changes at the protein level, the abundance of cytoplasmic EpeX pre-pro-peptide, and the EpeE epimerase was quantified using targeted mass spectrometry (MS) in wild type, as well as *kre*, and *spoVG* mutants. Samples were taken after 3, 4, 5, and 6 hours of cultivation in DSM medium. In the wild type, EpeX levels increased steadily over time, as shown by a positive log_2_ fold change relative to the 4 h timepoint of the corresponding strain (Figure 8a). This increase correlates with a strong *epeX* promoter activity and supports active toxin production during entry into stationary phase. In the *kre* mutant, EpeX abundance decreased nearly two-fold by 6 hours compared to wild type, despite the repressive effect on EPE production (Figure 8a). This suggests that Kre may contribute to transcript stability, possibly by preventing the mRNA from degradation or maintaining a favourable secondary structure for translation. Notably, across all measured time points, EpeX levels in the *spoVG* mutant were consistently lower than the wild type, with an approximately four-fold decrease observed at the 6 h time point (Figure 8a).

**Figure 8:**
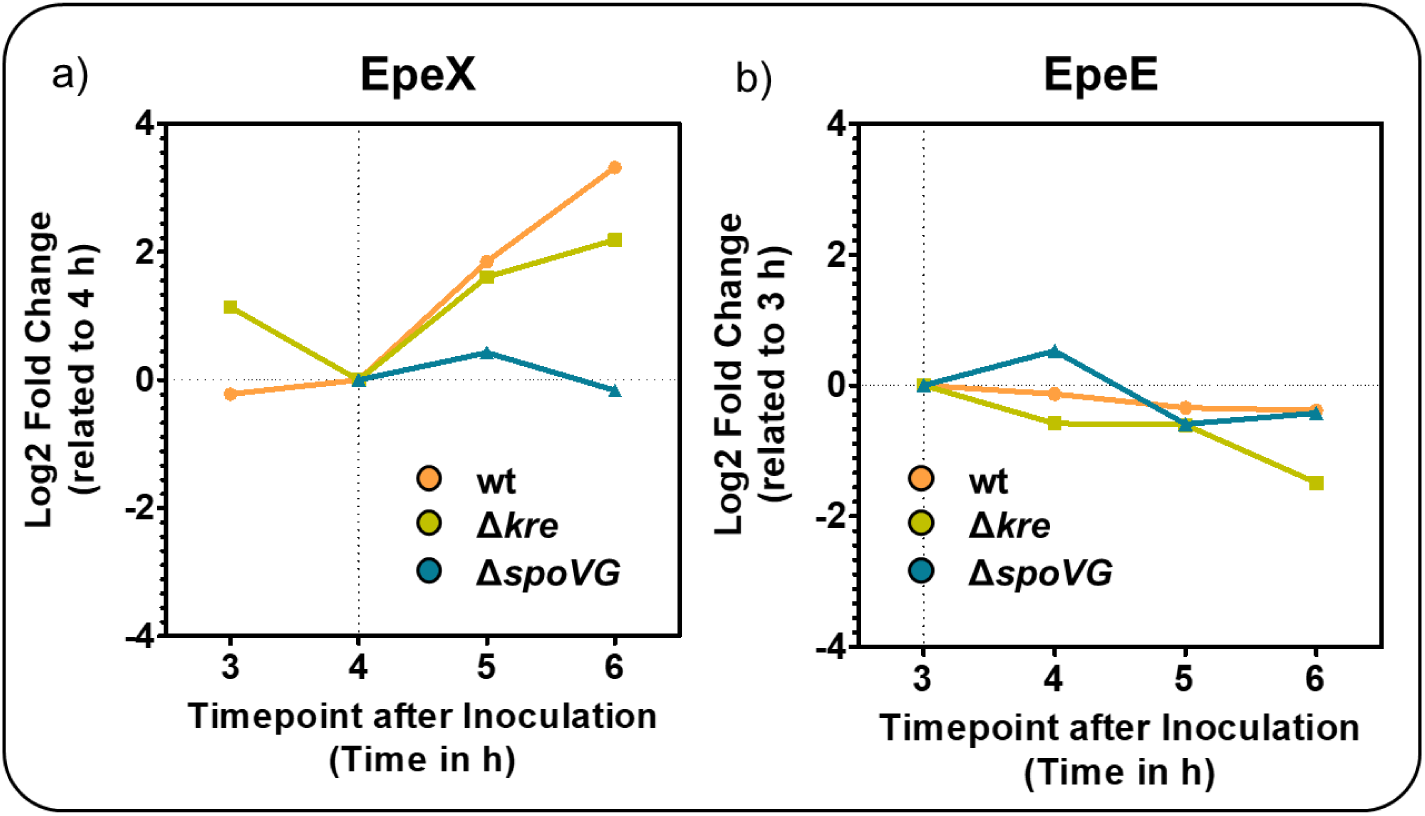
Temporal changes in EpeX and EpeE abundance during cultivation in DSM medium. **a)** The log_2_ fold change of EpeX abundance at 3, 4, 5, and 6 h relative to the 4 h time point, calculated for each strain: wild type, *kre* and *spoVG* mutant. **b)** Log_2_ fold change of EpeE abundance over time relative to the 3 h time point, calculated separately for each strain. Mass spectrometry was performed with biological triplicates. Note that fold changes represent relative changes within each strain. The absolute EpeX and EpeE abundances at the respective reference time points are not depicted.

This strongly supports the conclusion that SpoVG is essential for maintaining *epeX* at the post-transcriptional level, potentially by protecting the mRNA from RNases or by structural rearrangements. The complete loss of EpeX accumulation in this mutant reinforces its central activating role. MS analysis under the same conditions verified the presence of SpoVG and Kre in the wild type, indicating that the observed regulatory effects on *epeX* expression arise from their functional activity (Figure S11).

For EpeE, protein levels in the wild type remained relatively stable over time. In the *spoVG* mutant, EpeE abundance was slightly increased at 4 hours, but otherwise comparable to the wild type (Figure 8b). In contrast, the *kre* mutant displayed a gradual decrease in EpeE abundance, resulting in only ∼ 50% of EpeE after 6 hours compared to the wild type. This indicates that Kre may also facilitate *epeE* translation, however, to a lesser extent than SpoVG (Figure 8b). Collectively, these results demonstrate that SpoVG and Kre are involved in post-transcriptional regulation of EPE biosynthesis. SpoVG emerges as an essential activator, presumably stabilising the *epeX* transcript and ensuring robust protein accumulation. Kre, though primarily acting as a repressor, may support basal transcript integrity and/or fine-tune the translation process. These findings highlight the precision of coordinated toxin production with developmental states, environmental cues, and regulatory hierarchies in *B. subtilis*.

## DISCUSSION

Bacterial populations often face the challenge of balancing cooperative community behaviours with competition for limited resources. In *B. subtilis*, this balance is achieved through differentiation into subpopulations such as matrix producers, competent cells and cannibals (62,63). The production of the cannibalism toxins - SDP, SKF, and EPE - has been viewed as a last exit strategy for delaying or even preventing the highly energy-demanding and irreversible process of endospore formation (3,2). However, this study revealed a direct regulatory connection between competence and cannibalism raising new questions about the broader ecological role of the epipeptide EPE in adaptative stress response.

While toxin-mediated cell lysis provides access to nutrients for uptake by the toxin-producing cells, it is also accompanied by the release of extracellular DNA (2,3). Competent cells can subsequently take up this free DNA, which offers significant advantage by maintaining genetic diversity and providing a metabolically valuable source of nucleotides (39). Thus, the physiological interdependence of cannibalism and competence goes beyond the regulatory connection, suggesting a tightly coordinated and physiologically most meaningful developmental strategy (Figure 9a).

**Figure 9:**
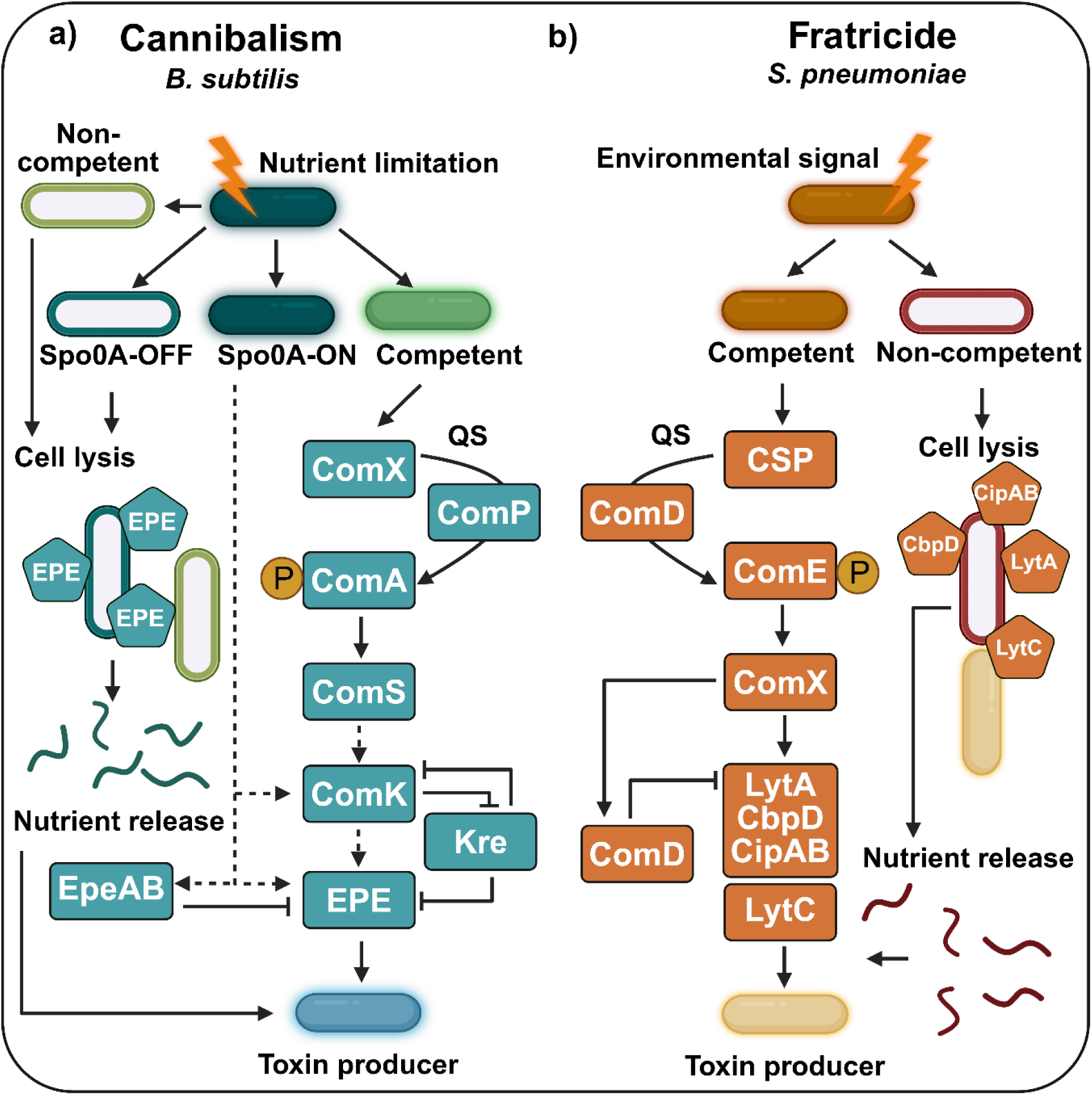
Cannibalism in *B. subtilis* and fratricide in *S. pneumoniae*. **a)** Cannibalism in *B. subtilis*. Under nutrient stress, cells either initiate the Spo0A-activating phosphorelay (Spo0A-ON) or remain in a non-sporulating state (Spo0A-OFF). Further nutrient limitation can induce competence, generating a heterologous population of competent and non-competent cells. Competent cells accumulate the quorum sensing (QS) signal peptide ComX, which activates ComP to phosphorylate ComA (ComA∼P). ComA∼P induces comS expression, which stabilises ComK. ComK, in turn, activates EPE production by repressing *kre*. Further Spo0A indirectly activates EPE production and expression of the autoimmunity EpeAB, as well as ComK by repressing the transition state regulator AbrB. Subsequently, EPE producing cells lyse Spo0A-OFF and non-competent cells. Nutrients released from the lysed cells support the growth and survival of the toxin producers. **b)** Fratricide in *S. pneumonia*. Favourable conditions and specific environmental signals trigger the development of competence in a subpopulation of cells, leading to the accumulation of competence-stimulating peptide (CSP). CSP activates ComD, which phosphorylates ComE (ComE∼P). ComE∼P leads to expression of ComX, which drives the production of fratricide factors, including CbpD, LytA, and CibAB. ComM provides immunity to competent cells against these toxins. LytC, which requires direct cell-cell contact but is independent of CSP regulation, further contributes to lysis. Together, these toxins selectively lyse non-competent cells, releasing nutrients that can be utilised by the competent, toxin producing population.

This kind of coupling between competence and cell lysis is not unique to *B. subtilis* (64). Originally identified in *Streptococcus pneumoniae*, a comparable strategy termed fratricide, is deeply integrated into the competence developmental program of this organism (65,66). In *S. pneumoniae*, competent cells lyse non-competent siblings to assess extracellular DNA (Figure 9b). While *B. subtilis* initiates cannibalism in response to nutrient limitation, mediated by both the Spo0A-activating phosphorelay and competence development (Figure 9a)*, S. pneumoniae* enters competence during exponential growth under favourable conditions, governed by quorum sensing alone (Figure 9b) (64). Competence is triggered by the accumulation of the 17-aa competence-stimulating peptide (CSP), encoded by *comC* (67–69). CSP is synthesized as a precursor peptide and further processed, as well as exported by the ComAB secretion system (67). At high cell densities, CSP reaches a threshold concentration and activates the membrane-bound histidine kinase ComD, which auto-phosphorylates and transfers the phosphate to its response regulator ComE (Figure 9b) (67,70).

Activated ComE induces transcription of ∼20 early competence genes, including those of the *comAB* and *comCDE* operons (71). Among these early genes is *comX*, encoding an alternative sigma factor that controls expression of over 60 late competence genes involved in DNA uptake and recombination (72). Importantly, ComX also initiates expression of the murein hydrolase CbpD and the autolysin LytA, as well as the bacteriocin peptides CibAB (65,73). The lysozyme LytC, although not regulated by CSP, also contributes to this fratricidal lysis, which requires cell-to-cell contact (Figure 9b). Fratricide in *S. pneumonia* is thought to enhance horizontal gene transfer, promote genetic diversity, and possibly contribute to pathogenicity by releasing virulence factors.

The findings presented in this work analysed in this context, cast a new light on the physiological role of the EPE toxin in *B. subtilis*. Instead of acting as a third cannibalism factor responsible for delaying sporulation, it might rather function as a fratricide toxin that is tightly linked to competence development. This hypothesis is supported by the similarities of the regulatory mechanisms with the canonical fratricide mechanism of *S. pneumoniae*, including quorum sensing and control by early competence genes (Figure 9). Complementing these pathways, *B. subtilis* also produces autolysins, encoded by the *lytABC* operon, which actively promote cell lysis and DNA release (74).

Beyond transcriptional control, post-transcriptional regulation constitutes another critical layer of gene expression control, encompassing mechanisms that govern RNA stability, processing, translation efficiency, and degradation (75–77). RNA binding proteins (RBPs) are key to these processes. By interacting with specific RNA sequences or structural motifs to modulate transcript fate, RBPs facilitate sRNA-mRNA interactions, remodel RNA secondary structures, and coordinate access to ribonucleases, thereby playing essential roles in shaping transcriptome dynamics (78,79). *B. subtilis* employs multiple RBPs with distinct regulatory functions. Among these, this work has revealed that SpoVG and Kre have emerged as key players in post-transcriptional control of *epeX* (Figure 5 & Figure 6). We demonstrated that SpoVG and Kre directly bind to the *epeX* mRNA, coordinating opposing regulatory effects. SpoVG is absolutely required for EPE production, likely stabilising the *epeX* transcript, while Kre appears to function antagonistically, potentially by modulating *epeX* and *epeE* expression (Figure 5, Figure 6, and Figure 7).

SpoVG has emerged as a pleiotropic regulator involved in virulence, stress adaptation, and gene expression in species such as *B. burgdorferi*, *S. aureus*, *L. monocytogenes*, and *B. anthracis* (80,81,57,82). The literature currently reports SpoVG to act as a dual DNA- and RNA-binding protein that appears to serve as a multifunctional regulator across diverse bacterial species (83). But, to date this has only been tested in vitro. However, work presented by Saylor et al. suggests its preference for RNA (83).

Here, SpoVG has been identified as a critical post-transcriptional regulator for EPE biosynthesis. SpoVG binds with high affinity to the 5’-end of the *epeX* transcript (Figure 6). Deletion of *spoVG* resulted in a complete loss of EPE-mediated stress response and a marked reduction in EpeX protein levels during growth, implicating SpoVG as essential for *epeX* transcript stability and subsequent toxin expression (Figure 7 & Figure 8). Direct binding of SpoVG to the *epeX* RNA was revealed by EMSA assays (Figure S7d).

Additionally, SPR analysis identified multiple binding sites across the complete length of the *epeX* transcript (Figure 6 & Figure S8). These interactions were characterised by varying association and dissociation kinetics, indicating a general affinity of SpoVG for the *epeX* mRNA. But the exceptionally high RNA-binding affinity of SpoVG complicates the distinction between specific and unspecific interactions as it engages with diverse RNA structures and sequences. Future competition assays with competitor RNAs and proteins will help delineate target specificity.

Notably, the most stable interaction, reflected by the lowest dissociation rate, was observed at the 5’-end of the transcript, likely holding the highest biological relevance (Figure 6 & Figure S8). These findings suggest that SpoVG might protect the *epeX* mRNA from 5’-3’-exonucleolytic degradation, e. g. by RNase J1, or following endoribonucleolytic cleavage by RNases such as RNase Y and RNase III (84,85). Sequence specificity is supported by conserved residues across SpoVG homologues (55), reinforcing the hypothesis that SpoVG selectively stabilises the *epeX* mRNA through defined interactions. At the same time, overall transcript stabilisation potentially also results from multiple SpoVG binding events distributed along the full-length *epeX* transcript (Figure 6).

Both *spoVG* and *epeX* are transcriptionally repressed by AbrB and indirectly activated by Spo0A∼P (86,6). This coordinated expression ensures that *epeX* mRNA accumulates during the transition phase, facilitating EPE biosynthesis in response to environmental signals. Recent evidence further underscores the relevance of this post-transcriptional control in *B. subtilis* and has demonstrated a functional decoupling of transcription and translation (87). This decoupling increases the necessity for precise regulation of nascent RNA, predominantly fulfilled by RBPs such as SpoVG and Kre.

In contrast to the EPE-promoting function of SpoVG, Kre negatively influences *epeX* expression. In this study, Kre was identified for the first time in *B. subtilis* as an RBP that associates with the *epeX* transcript, most likely at the terminator region (Figure S6). This interaction appears to promote enhanced transcription termination, thereby supressing EPE production. This repression is relieved during the competence state, as *kre* is transcriptionally downregulated by ComK (45). Notably, Kre homologues are restricted to species that also encode ComK, including *B. licheniformis* and *B. anthracis* (45). This distribution suggests that Kre plays an important and potentially conserved role in the control of competence.

Kre is highly positively charged, which presumably promotes strong electrostatic interactions with negatively charged nucleic acids. Its structure includes a potential helix-turn-helix motif, commonly associated with nucleic acid binding (Figure S5) (49). While Kre lacks detectable nuclease activity, it influences transcript stabilities (45,48). Based on its strong affinity for binding at the *epeX* terminator region, Kre is proposed to function as a post-transcriptional regulator that enhances rho-independent termination and modulates mRNA stability. Kre potentially stabilises the terminator hairpin of *epeX*, promoting efficient transcriptional termination and suppressing readthrough into the downstream *epeEP* locus, thereby preventing EPE production (Figure 5). A modular, cooperative binding mechanism is conceivable, in which Kre initially recognises a stable RNA secondary structure at the terminator, followed by additional contacts along the transcript that stabilise the protein-RNA complex. The multiplicity of kinetic populations observed in SPR analysis supports this model, in which rapid, reversible binding at lower affinity sites might be followed by high-affinity, stable interactions at the IGR*_epeXE_*, ultimately interfering with transcript elongation or stability (Figure 5).

These findings uncover a previously unrecognised regulatory interdependency between SpoVG, ComK, and Kre for controlling EPE production. This multilayered control ensures the precise timing and fine-tuning of *epeXEP* expression, allowing EPE biosynthesis to rapidly respond to environmental stressors. During exponential growth, EPE production is actively repressed through both transcriptional and post-transcriptional control that tightly control expression. At the transcriptional level, the global regulator AbrB is expressed and inhibits transcription of the *epeXEPAB* operon at P*_epeX_* (Figure 10) (88). Simultaneously, repression is reinforced post-transcriptionally: The RNA-binding protein Kre is present due to AbrB-mediated repression of *comK*, encoding the master regulator of competence (Figure 10) (45). Kre binds within the IGR*_epeXE_*, thereby stabilising the terminator hairpin structure (Figure 5). This likely promotes premature transcription termination and hence preventing the full-length transcription of the *epe* operon. Translation of *epeX* is further inhibited by ribosome stalling at its final codon (CAT), obstructing the synthesis of the pre-pro peptide EpeX (89,13). Moreover, RNA stability is compromised as the 5’-untranslated region of *epeX* remains unprotected in this phase due to *spoVG* repression by AbrB (Figure 10) (36). Consequently, the *epeX* transcript is more accessible for RNases and hence degradation.

**Figure 10:**
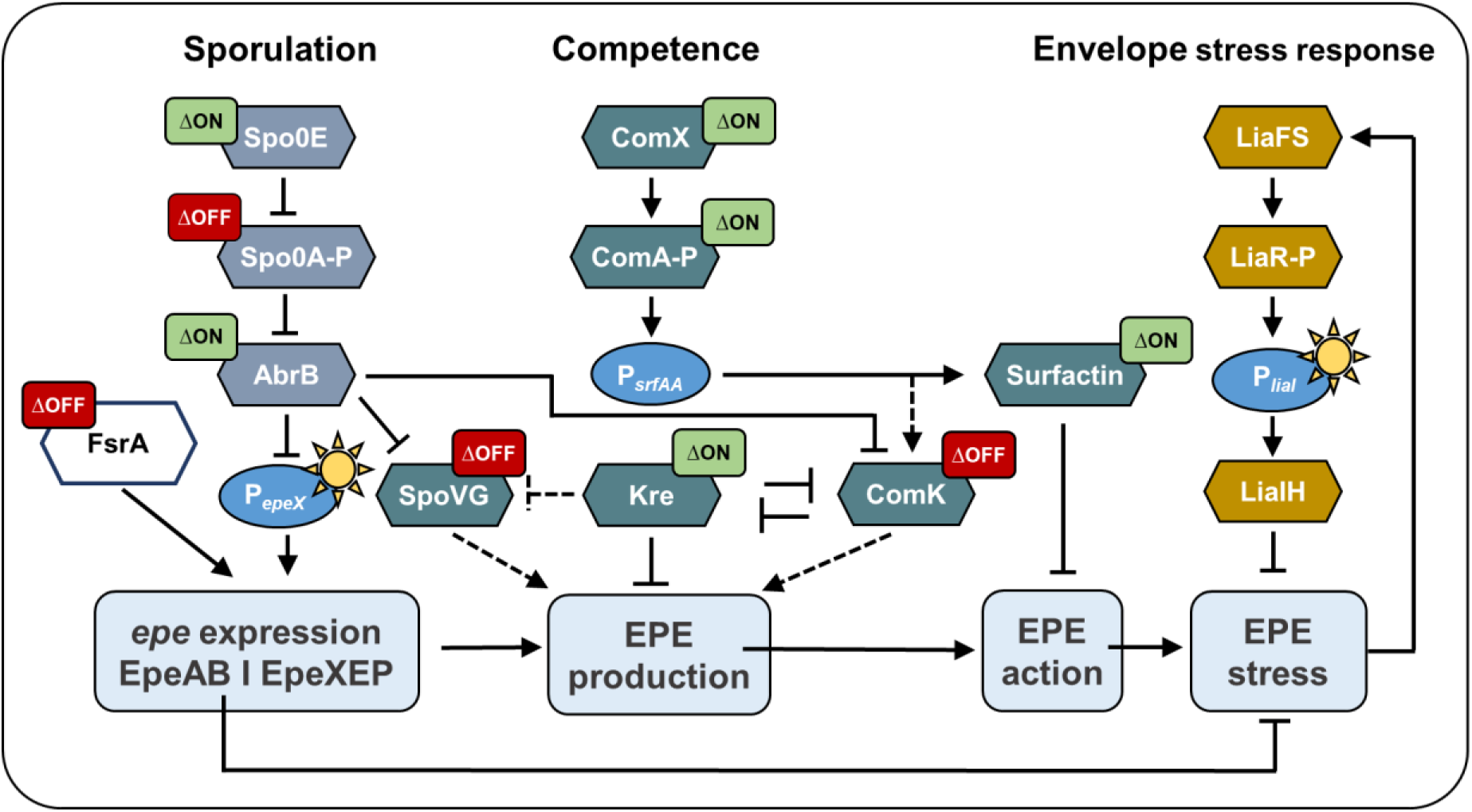
Regulatory pathways controlling EPE activation. The impact of specific gene deletion on P*_liaI_*-*lux* activity is shown using colour coding: Green (“ON”) indicating comparable or up to four times higher activity than the wild type. Red (“OFF”) imply a complete loss of P*_liaI_*-*lux* activity, similar to the effect of an *epeX* deletion. P*_srfAA_* both leads to ComK activation (via expression of *comS*) and surfactin production. The two *lux*-reporter fusions relevant to build up the regulatory scheme are indicated by the sun symbols.

As cells transition into stationary phase and encounter nutrient limitation, global changes in gene expression activate alternative sets of regulators. ComK becomes active and represses *kre* expression (Figure 10) (90,45), thereby removing the inhibitory influence of Kre on the *epeX* transcript. This change likely allows full-length transcription of the *epeXEPAB* operon to proceed. RNA stability increases further due to enhanced SpoVG levels, which was previously repressed by AbrB. SpoVG binds across and thereby protecting the *epeX* transcript, particularly at its 5’-end to prevent degradation (Figure S8 & Figure 6). Binding of SpoVG could also lead to structural rearrangement, which further stabilises the transcript and increases translation efficiency.

This carefully orchestrated regulation is temporally confined to a window just before the accumulation of Spo0A∼P reaches levels that commit cells to sporulation. During this transition phase, intermediate Spo0A∼P levels fine-tune its activity, creating a transient state in which competence development, toxin production, and early sporulation signals can coexist (Figure 10) (91–93). Once Spo0A∼P surpasses a critical threshold, cells irreversibly commit to sporulation and the opportunity for EPE production closes (18). Once synthesised, EPE executes its biological effect by perturbing the cytoplasmic membrane and ultimately provoking cell lysis, which releases nutrients and extracellular DNA (Figure 10) (6,3). This not only enhances population-level survival under stress but also contributes to genetic diversification and ecological competitiveness (3).

Taken together, EPE biosynthesis represents an example for how complex developmental decision processes are mediated by tightly coordinating transcriptional and post-transcriptional regulation. Rather than relying on a single global RNA chaperone, *B. subtilis* employs specialised RBPs such as SpoVG and Kre to precisely gate toxin production within a narrow physiological window. This multilayered control ensures that EPE is produced only when its benefits outweigh its risks and costs, thereby aligning individual cell fates with population-level survival strategies. More broadly, these findings highlight post-transcriptional regulation as a central determinant of bacterial developmental plasticity and underscore the importance of RNA-centred regulatory networks in shaping adaptive responses to environmental stress. Beyond expanding our understanding of bacterial gene regulation, our findings highlight EPE production as a prime example for how a multilayered control is employed for coordinating multicellular differentiation, endospore formation, and competence for genetic transformation with a bacterial form of programmed cell death. Future studies need to aim at dissecting the temporal and molecular dynamics of this network to mechanistically unravel the regulatory logic underlying microbial development and toxin-mediated cooperation.

## ACKNOWLEDGEMENTS

SPR analyses were performed in the Bioanalytics core facility of the JGU Biocenter. Jürgen Bartel is acknowledged for his technical support. We further acknowledge *BioRender*, which was applied to generate schematic figures [Licence: 2155-6232].

## AUTHOR CONTRIBUTIONS

<u>Sarah Miercke</u>: Conceptualisation, Experimental Work, Data analysis and interpretation, Writing - Original Draft. <u>Kirsten Schaubruch</u>: Experimental Work. <u>Sandra Maaß</u>: Experimental Work, Data analysis and interpretation, Writing - Review & Editing. <u>Anne-Katrin Rußeck</u>: Experimental Work. <u>Andreas Lawaetz</u>: Experimental Work. <u>Emma L. Denham</u>: Supervision, Writing - Review & Editing. <u>Ralf Heermann</u>: Data analysis and interpretation, Writing - Review & Editing. <u>Thorsten Mascher</u>: Conceptualisation, Supervision, Funding, Writing - Review & Editing.

## SUPPLEMENTARY DATA

Supplementary Data are available at NAR online.

## DATA AVAILABILITY

The MS data and Skyline files are available on Panorama Public (permanent link: https://panoramaweb.org/W5CMfT.url) and are associated with the ProteomeXchange ID PXD075352. Reviewer access is available via the permanent link using the following credentials, Password: aG4KSRW7rl8uV!

## REFERENCES

1. Earl, A.M., Losick, R. and Kolter, R. (2008) Ecology and Genomics of *Bacillus subtilis*, Trends Microbiol., 16, 269–275. First published on May 28, 2008.

2. González-Pastor, J.E., Hobbs, E.C. and Losick, R. (2003) Cannibalism by Sporulating Bacteria, Science, 301, 510–513.

3. González-Pastor, J.E. (2011) Cannibalism: A Social Behavior in Sporulating *Bacillus subtilis*, FEMS Microbiol. Rev., 35, 415–424.

4. Liu, W.-T., Yang, Y.-L. and Xu, Y. et al. (2010) Imaging Mass Spectrometry of Intraspecies Metabolic Exchange Revealed the Cannibalistic Factors of *Bacillus subtilis*, Proc. Natl. Acad. Sci. U.S.A., 107, 16286–16290.

5. Popp, P.F., Benjdia, A. and Strahl, H. et al. (2020) The Epipeptide YydF Intrinsically Triggers the Cell Envelope Stress Response of *Bacillus subtilis* and Causes Severe Membrane Perturbations, Front. Microbiol., 11.

6. Popp, P.F., Friebel, L. and Benjdia, A. et al. (2021) The Epipeptide Biosynthesis Locus *epeXEPAB* Is Widely Distributed in Firmicutes and Triggers Intrinsic Cell Envelope Stress, Microb. physiol., 31, 306–318. First published on Jun 11, 2021.

7. Benjdia, A., Guillot, A. and Ruffié, P. et al. (2017) Post-translational Modification of Ribosomally Synthesized Peptides by a Radical SAM Epimerase in *Bacillus subtilis*, Nat. Chem., 9, 698–707.

8. Kubiak, X., Polsinelli, I. and Chavas, L.M.G. et al. (2024) Structural and Mechanistic Basis for RiPP Epimerization by a Radical SAM Enzyme, Nat. Chem. Biol., 20, 382–391. First published on Dec 29, 2023.

9. Kalamara, M., Abbott, J. and Sukhodub, T. et al. (2023) The Putative Role of the Epipeptide EpeX in *Bacillus subtilis* Intra-Species Competition, Microbiology, 169.

10. Nicolas, P., Mäder, U. and Dervyn, E. et al. (2012) Condition-Dependent Transcriptome Reveals High-Level Regulatory Architecture in *Bacillus subtilis*, Science, 335, 1103–1106.

11. Dérozier, S., Nicolas, P. and Mäder, U. et al. (2021) Genoscapist: Online Exploration of Quantitative Profiles Along Genomes via Interactively Customized Graphical Representations, Bioinformatics, 37, 2747–2749.

12. Mandell, Z.F., Vishwakarma, R.K. and Yakhnin, H. et al. (2022) Comprehensive Transcription Terminator Atlas for *Bacillus subtilis*, Nat. Microbiol., 7, 1918–1931.

13. Miercke, S., Ghandour, R. and Papenfort, K. et al. (2025) The FsrA-Mediated Iron-Sparing Response Regulates the Biosynthesis of the Epipeptide EPE in Bacillus subtilis, Mol. Micro. First published on Dec 17, 2025.

14. Vlamakis, H., Aguilar, C. and Losick, R. et al. (2008) Control of Cell Fate by the Formation of an Architecturally Complex Bacterial Community, Genes Dev., 22, 945–953.

15. Altenbuchner, J. (2016) Editing of the *Bacillus subtilis* Genome by the CRISPR-Cas9 System, Appl. Environ. Microbiol., 82, 5421–5427.

16. Trach, K.A. and Hoch, J.A. (1993) Multisensory Activation of the Phosphorelay Initiating Sporulation in *Bacillus subtilis*: Identification and Sequence of the Protein Kinase of the Alternate Pathway, Mol. Micro., 8, 69–79.

17. Hoch, J.A. (1993) The Phosphorelay Signal Transduction Pathway in the Initiation of *Bacillus subtilis* Sporulation, J. Cell. Biochem., 51, 55–61.

18. Burbulys, D., Trach, K.A. and Hoch, J.A. (1991) Initiation of Sporulation in *B. subtilis* is Controlled by a Multicomponent Phosphorelay, Cell, 64, 545–552.

19. Perego, M., Spiegelman, G.B. and Hoch, J.A. (1988) Structure of the Gene for the Transition State Regulator, AbrB: Regulator Synthesis is Controlled by the *spo0A* Sporulation Gene in *Bacillus subtilis*, Mol. Micro., 2, 689–699.

20. Schultz, D., Wolynes, P.G. and Ben Jacob, E. et al. (2009) Deciding Fate in Adverse Times: Sporulation and Competence in *Bacillus subtilis*, Proc. Natl. Acad. Sci. U.S.A., 106, 21027–21034. First published on Dec 7, 2009.

21. Piggot, P.J. and Coote, J.G. (1976) Genetic Aspects of Bacterial Endospore Formation, Bact. Rev., 40, 908–962.

22. Stragier, P. and Losick, R. (1996) Molecular Genetics of Sporulation in *Bacillus subtilis*, Annu. Rev. Genet., 30, 297–41.

23. McKenney, P.T., Driks, A. and Eichenberger, P. (2013) The *Bacillus subtilis* Endospore: Assembly and Functions of the Multilayered Coat, Nat. Rev. Microbiol., 11, 33–44. First published on Dec 3, 2012.

24. Checinska, A., Paszczynski, A. and Burbank, M. (2015) *Bacillus* and Other Spore-Forming Genera: Variations in Sesponses and Mechanisms for Survival, Annu. Rev. Food Sci. Technol., 6, 351–369. First published on Feb 20, 2015.

25. Setlow, P. (2014) Spore Resistance Properties, Microbiology spectrum, 2.

26. Parker, G.F., Daniel, R.A. and Errington, J. (1996) Timing and Genetic Regulation of Commitment to Sporulation in *Bacillus subtilis*, Microbiology, 142, 3445–3452.

27. Shank, E.A. and Kolter, R. (2011) Extracellular Signaling and Multicellularity in *Bacillus subtilis*, Curr. Opin. Microbiol., 14, 741–747. First published on Oct 23, 2011.

28. Sambrook, J. and Russell, D.W. (2001) Molecular Cloning. A laboratory manual. Cold Spring Harbor Laboratory Press, Cold Spring Harbor, N.Y.

29. Harwood, C.R. (ed.) (1990). Molecular Biological Methods for Bacillus. Wiley, Chichester.

30. Popp, P.F., Dotzler, M. and Radeck, J. et al. (2017) The *Bacillus* BioBrick Box 2.0: Expanding the Genetic Toolbox for the Standardized Work With *Bacillus subtilis*, Sci. Rep., 7, 15058. First published on Nov 8, 2017.

31. Radeck, J., Kraft, K. and Bartels, J. et al. (2013) The *Bacillus* BioBrick Box: Generation and Evaluation of Essential Genetic Building Blocks for Standardized Work with *Bacillus subtilis*, J. Biol. Eng., 7.

32. Fietze, T., Wilk, P. and Kabinger, F. et al. (2022) HUG Domain Is Responsible for Active Dimer Stabilization in an NrdJd Ribonucleotide Reductase, Biochemistry, 61, 1633–1641.

33. Bradford, M.M. (1976) A Rapid and Sensitive Method for the Quantitation of Microgram Quantities of Protein Utilizing the Principle of Protein-Dye Binding, Anal. Biochem., 72, 248–254.

34. Altschuh, D., Björkelund, H. and Strandgård, J. et al. (2012) Deciphering Complex Protein Interaction Kinetics Using Interaction Map, Biochem. Biophys. Res. Commun., 428, 74–79. First published on Oct 9, 2012.

35. Pino, L.K., Searle, B.C. and Bollinger, J.G. et al. (2020) The Skyline ecosystem: Informatics for quantitative mass spectrometry proteomics, Mass Spectrometry Rev., 39, 229–244. First published on Jul 9, 2017.

36. Chumsakul, O., Takahashi, H. and Oshima, T. et al. (2011) Genome-Wide Binding Profiles of the *Bacillus subtilis* Transition State Regulator AbrB and its Homolog Abh Reveals Their Interactive Role in Transcriptional Regulation, Nucleic Acids Res., 39, 414–428. First published on Sep 3, 2010.

37. van Sinderen, D., Luttinger, A. and Kong, L., et al. (1995) *comK* Encodes the Competence Transcription Factor, the Key Regulatory Protein for Competence Development in *Bacillus subtilis*, *Mol*. Micro., 15, 455–462.

38. Berka, R.M., Hahn, J. and Albano, M. et al. (2002) Microarray Analysis of the *Bacillus subtilis* K-state: Genome-Wide Expression Changes Dependent on ComK, Mol. Micro., 43, 1331–1345.

39. Finkel, S.E. and Kolter, R. (2001) DNA as a Nutrient: Novel Role for Bacterial Competence Gene Homologs, J. Bacteriol., 183, 6288–6293.

40. Ansaldi, M., Marolt, D. and Stebe, T. et al. (2002) Specific Activation of the *Bacillus* Quorum-Sensing Systems by Isoprenylated Pheromone Variants, *Mol*. Micro., 44, 1561–1573.

41. Weinrauch, Y., Penchev, R. and Dubnau, E. et al. (1990) A *Bacillus subtilis* Regulatory Gene Product for Genetic Competence and Sporulation Resembles Sensor Protein Members of the Bacterial Two-Component Signal-Transduction Systems, Genes Dev., 4, 860–872.

42. Roggiani, M. and Dubnau, D. (1993) ComA, a Phosphorylated Response Regulator Protein of *Bacillus subtilis*, Binds to the Promoter Region of *srfA*, J. Bacteriol., 175, 3182–3187.

43. Cosmina, P., Rodriguez, F. and Ferra, F. de et al. (1993) Sequence and Analysis of the Genetic Locus Responsible for Surfactin Synthesis in *Bacillus subtilis*, Mol. Micro., 8, 821–831.

44. Comella, N. and Grossman, A.D. (2005) Conservation of Genes and Processes Controlled by the Quorum Response in Bacteria: Characterization of Genes Controlled by the Quorum-Sensing Transcription Factor ComA in *Bacillus subtilis*, Mol. Micro., 57, 1159–1174.

45. Gamba, P., Jonker, M.J. and Hamoen, L.W. (2015) A Novel Feedback Loop That Controls Bimodal Expression of Genetic Competence, PLoS Genet., 11, e1005047. First published on Jun 25, 2015.

46. van Sinderen, D. and Venema, G. (1994) *comK* Acts as an Autoregulatory Control Switch in the Signal Transduction Route to Competence in *Bacillus subtilis*, J. Bacteriol., 176, 5762–5770.

47. Ogura, M., Yamaguchi, H. and Kobayashi, K. et al. (2002) Whole-Genome Analysis of Genes Regulated by the *Bacillus subtilis* Competence Transcription Factor ComK, J. Bacteriol., 184, 2344–2351.

48. Pi, H., Weiss, A. and Laut, C.L. et al. (2022) An RNA-Binding Protein Acts as a Major Post-Transcriptional Modulator in *Bacillus anthracis*, Nat. Commun., 13, 1491. First published on Mar 21, 2022.

49. Aravind, L., Anantharaman, V. and Balaji, S. et al. (2005) The Many Faces of the Helix-Turn-Helix Domain: Transcription Regulation and Beyond, FEMS Microbiol. Rev., 29, 231–262.

50. Hambraeus, G., Wachenfeldt, C. von and Hederstedt, L. (2003) Genome-Wide Survey of mRNA Half-Lives in *Bacillus subtilis* Identifies Extremely Stable mRNAs, Mol. Gen. Genomics, 269, 706–714.

51. Jenniches, L., Michaux, C. and Popella, L. et al. (2024) Improved RNA stability estimation through Bayesian modeling reveals most Salmonella transcripts have subminute half-lives, Proc. Natl. Acad. Sci. U.S.A., 121, e2308814121. First published on Mar 25, 2024.

52. Lehnik-Habrink, M., Lewis, R.J. and Mäder, U. et al. (2012) RNA Degradation in *Bacillus subtilis*: An Interplay of Essential Endo- and Exoribonucleases, *Mol*. Micro., 84, 1005–1017. First published on May 8, 2012.

53. Ul Haq, I., Müller, P. and Brantl, S. (2020) Intermolecular Communication in *Bacillus subtilis*: RNA-RNA, RNA-Protein and Small Protein-Protein Interactions, Front. Mol. Biosci., 7, 178. First published on Aug 7, 2020.

54. Matsuno, K. and Sonenshein, A.L. (1999) Role of SpoVG in Asymmetric Septation in *Bacillus subtilis*, J. Bacteriol., 181, 3392–3401.

55. Jutras, B.L., Chenail, A.M. and Rowland, C.L. et al. (2013) Eubacterial SpoVG Homologs Constitute a New Family of Site-Specific DNA-Binding Proteins, PloS One, 8, e66683. First published on Jun 20, 2013.

56. Burke, T.P. and Portnoy, D.A. (2016) SpoVG Is a Conserved RNA-Binding Protein That Regulates *Listeria monocytogenes* Lysozyme Resistance, Virulence, and Swarming Motility, mBio, 7, e00240. First published on Apr 5, 2016.

57. Xu, L., Zhang, X. and Wang, W. et al. (2025) The Global Regulator SpoVG is Involved in Biofilm Formation and Stress Response in Foodborne *Staphylococcus aureus*, Int. J. Food Microbio., 428, 110997. First published on Nov 28, 2024.

58. Tan, X., Zheng, N. and Hayashi, K., et al. (2008) Worldwide Protein Data Bank.

59. Hellman, L.M. and Fried, M.G. (2007) Electrophoretic Mobility Shift Assay (EMSA) for Detecting Protein-Nucleic Acid Interactions, Nat. Protoc., 2, 1849–1861.

60. Woodson, S.A. and Koculi, E. (2009) Analysis of RNA Folding by Native Polyacrylamide Gel Electrophoresis, Methods Enzymol., 469, 189–208. First published on Nov 17, 2009.

61. Ryder, S.P., Recht, M.I. and Williamson, J.R. (2008) Quantitative Analysis of Protein-RNA Interactions by Gel Mobility Shift, Methods in molecular biology (Clifton, N.J.), 488, 99–115.

62. Veening, J.-W., Hamoen, L.W. and Kuipers, O.P. (2005) Phosphatases Modulate the Bistable Sporulation Gene Expression Pattern in *Bacillus subtilis*, *Mol*. Micro., 56, 1481–1494.

63. López, D., Vlamakis, H. and Losick, R. et al. (2009) Cannibalism Enhances Biofilm Development in *Bacillus subtilis*, *Mol*. Micro., 74, 609–618. First published on Sep 22, 2009.

64. Claverys, J.-P. and Håvarstein, L.S. (2007) Cannibalism and Fratricide: Mechanisms and *Raisons d’être*, Nat. Rev. Microbiol., 5, 219–229.

65. Guiral, S., Mitchell, T.J. and Martin, B. et al. (2005) Competence-Programmed Predation of Non-Competent Cells in the Human Pathogen *Streptococcus pneumoniae*: Genetic Requirements, Proc. Natl. Acad. Sci. U.S.A., 102, 8710–8715. First published on May 31, 2005.

66. Steinmoen, H., Knutsen, E. and Håvarstein, L.S. (2002) Induction of Natural Competence in *Streptococcus pneumoniae* Triggers Lysis and DNA Release From a Subfraction of the Cell Population, Proc. Natl. Acad. Sci. U.S.A., 99, 7681–7686.

67. Håvarstein, L.S., Coomaraswamy, G. and Morrison, D.A. (1995) An Unmodified Heptadecapeptide Pheromone Induces Competence for Genetic Transformation in *Streptococcus pneumoniae*, Proc. Natl. Acad. Sci. U.S.A., 92, 11140–11144.

68. Whatmore, A.M., Barcus, V.A. and Dowson, C.G. (1999) Genetic Diversity of the Streptococcal Competence (*com*) Gene Locus, J. Bacteriol., 181, 3144–3154.

69. Håvarstein, L.S., Hakenbeck, R. and Gaustad, P. (1997) Natural Competence in the Genus *Streptococcus*: Evidence That Streptococci can Change Pherotype by Interspecies Recombinational Exchanges, J. Bacteriol., 179, 6589–6594.

70. Håvarstein, L.S., Gaustad, P. and Nes, I.F. et al. (1996) Identification of the Streptococcal Competence-Pheromone Receptor, *Mol*. Micro., 21, 863–869.

71. Ween, O., Gaustad, P. and Håvarstein, L.S. (1999) Identification of DNA Binding Sites for ComE, a Key Regulator of Natural Competence in *Streptococcus pneumoniae*, Mol. Micro., 33, 817–827.

72. Claverys, J.-P., Prudhomme, M. and Martin, B. (2006) Induction of Competence Regulons as a General Response to Stress in Gram-Positive Bacteria, Annu. Rev. Microbiol., 60, 451–475.

73. Kausmally, L., Johnsborg, O. and Lunde, M. et al. (2005) Choline-Binding Protein D (CbpD) in *Streptococcus pneumoniae* is Essential for Competence-Induced Cell Lysis, J. Bacteriol., 187, 4338–4345.

74. Lazarevic, V., Margot, P. and Soldo, B. et al. (1992) Sequencing and Analysis of the *Bacillus subtilis lytRABC* Divergon: A Regulatory Unit Encompassing the Structural Genes of the N-Acetylmuramoyl-L-Alanine Amidase and Its Modifier, J. Gen. Microbiol., 138, 1949–1961.

75. Papenfort, K. and Melamed, S. (2023) Small RNAs, Large Networks: Posttranscriptional Regulons in Gram-Negative Bacteria, Annu. Rev. Microbiol., 77, 23–43. First published on Mar 21, 2023.

76. Trinquier, A., Durand, S. and Braun, F. et al. (2020) Regulation of RNA Processing and Degradation in Bacteria, BBA - Gene Regulatory Mechanisms, 1863, 194505.

77. Condon, C. and Bechhofer, D.H. (2011) Regulated RNA Stability in the Gram Positives, Curr. Opin. Microbiol., 14, 148–154. First published on Feb 19, 2011.

78. Christopoulou, N. and Granneman, S. (2022) The Role of RNA - Binding Proteins in Mediating Adaptive Responses in Gram-Positive Bacteria, FEBS J., 289, 1746–1764.

79. Sun, S., Li, X. and Zhai, J. et al. (2025) Orchestrating Nutrient Homeostasis: RNA-Binding Proteins as Molecular Conductors in Metabolic Disease Pathogenesis, Nutrients, 17. First published on Jul 19, 2025.

80. Chen, M., Lyu, Y. and Feng, E. et al. (2020) SpoVG is Necessary for Sporulation in *Bacillus anthracis*, Microorganisms, 8. First published on Apr 10, 2020.

81. Shi, C., Zheng, L. and Lu, Z. et al. (2023) The Global Regulator SpoVG Regulates *Listeria monocytogenes* Biofilm Formation, Microb. pathog., 180, 106144. First published on May 4, 2023.

82. Savage, C.R., Jutras, B.L. and Bestor, A. et al. (2018) *Borrelia burgdorferi* SpoVG DNA- and RNA-Binding Protein Modulates the Physiology of the Lyme Disease Spirochete, J. Bacteriol., 200. First published on May 24, 2018.

83. Saylor, T.C., Savage, C.R. and Krusenstjerna, A.C. et al. (2023) Quantitative Analyses of Interactions Between SpoVG and RNA/DNA, Biochem. Biophys. Res. Commun., 654, 40–46. First published on Feb 27, 2023.

84. Durand, S., Gilet, L. and Bessières, P. et al. (2012) Three Essential Ribonucleases-RNase Y, J1, and III-Control the Abundance of a Majority of *Bacillus subtilis* mRNAs, PLoS Genet., 8, e1002520. First published on Mar 8, 2012.

85. Mathy, N., Hébert, A. and Mervelet, P. et al. (2010) *Bacillus subtilis* Ribonucleases J1 and J2 Form a Complex With Altered Enzyme Behaviour, Mol. Micro., 75, 489–498. First published on Dec 16, 2009.

86. Zuber, P. and Losick, R. (1987) Role of AbrB in Spo0A- and Spo0B-Dependent Utilization of a Sporulation Promoter in *Bacillus subtilis*, J. Bacteriol., 169, 2223–2230.

87. Johnson, G.E., Lalanne, J.-B. and Peters, M.L. et al. (2020) Functionally Uncoupled Transcription-Translation in *Bacillus subtilis*, Nature, 585, 124–128. First published on Aug 26, 2020.

88. Butcher, B.G., Lin, Y.-P. and Helmann, J.D. (2007) The *yydFGHIJ* Operon of *Bacillus subtilis* Encodes a Peptide That Induces the LiaRS Two-Component System, J. Bacteriol., 189, 8616–8625.

89. Han, Y., Wang, B. and Agnolin, A. et al. (2025) Ribosome Pausing in Amylase Producing *Bacillus subtilis* During Long Fermentation, Microb. Cell Fact., 24.

90. Hamoen, L.W., Kausche, D. and Marahiel, M.A. et al. (2003) The *Bacillus subtilis* Transition State Regulator AbrB Binds to the -35 Promoter Region of *comK*, FEMS Microbiol. Lett., 218, 299–304.

91. Bettenworth, V., Steinfeld, B. and Duin, H. et al. (2019) Phenotypic Heterogeneity in Bacterial Quorum Sensing Systems, J. Mol. Biol., 431, 4530–4546. First published on Apr 30, 2019.

92. Kalamara, M., Spacapan, M. and Mandic-Mulec, I. et al. (2018) Social Behaviours by *Bacillus subtilis*: Quorum Sensing, Kin Discrimination and Beyond, *Mol*. Micro., 110, 863–878. First published on Nov 1, 2018.

93. Smits, W.K., Bongiorni, C. and Veening, J.-W. et al. (2007) Temporal Separation of Distinct Differentiation Pathways by a Dual Specificity Rap-Phr System in *Bacillus subtilis*, Mol. Micro., 65, 103–120.

