## Supplemental Information for "SpoVG and the Kre-ComK Regulatory Module Orchestrate Production of the EPE Toxin in *Bacillus subtilis*"

### Table of Contents

**Table S1:** Strains applied in this study.

**Table S2:** Plasmids used in this study.

**Table S3:** Oligonucleotides used in this study.

**Table S4:** Transition list of mass spectrometry analysis.

**Figure S1:** Timing of *epe* expression and EPE stress reponse.

**Figure S2:** Impact of sporulation related mutants on EPE production and stress response.

**Figure S3:** Quorum sensing networks and phosphorelay in *B. subtilis*.

**Figure S4:** Impact of competence related mutants on EPE production and stress response.

**Figure S5:** Purification and structural prediction of the RNA-binding protein Kre.

**Figure S6:** SPR analysis of Kre binding to truncated *epeX* RNA transcripts.

**Figure S7:** Structural characterisation of the RNA binding protein SpoVG and its interaction with the *epeX* RNA.

**Figure S8:** SPR analysis of SpoVG binding to truncated *epeX* RNA transcripts.

**Figure S9:** Effects of *comK*, *kre*, and *spoVG* single and combinatory mutants on EPE-mediated stress response.

**Figure S10:** Effect of *spoVG* deletion on *epeXEP* expression.

**Figure S11:** Temporal changes in Kre and SpoVG abundance during cultivation in DSM medium.

### Supplementary

**Table S1**: Strains applied in this study.

| Strain | Genotype* | Reference |
|---|---|---|
| DK1042 | *B. subtilis* NCIB3610 *comI* (Q12L) = DK1042 | Konkol et al. 2013 (1) |
| BL21 | *E. coli* overexpression strain | Laboratory stock |
| DH10*β* | *E. coli* laboratory wild type strain | Laboratory stock |
| DH5*α* | *E. coli* laboratory wild type strain | Laboratory stock |
| TMB6136 | DK1042 *sacA*::$P_{liaI}$-*luxABCDE_cat* | Laboratory stock |
| TMB6170 | DK1042 *sacA*::$P_{epeX}$-*luxABCDE_cat* | This study |
| TMB6217 | DK1042 *sacA*::$P_{liaI}$-*luxABCDE_cat*, *epeAB*::*spec* | This study |
| TMB6237 | DK1042 *sacA*::$P_{spo0A}$-*luxABCDE_cat* | This study |
| TMB6247 | DK1042 *sacA*::$P_{liaI}$-*luxABCDE_cat*, *epeAB*::*spec*, *epeX*::*mls* | This study |
| TMB6256 | DK1042 *sacA*::$P_{liaI}$-*luxABCDE_cat*, *epeAB*::*spec*, *srfAA*::*kan* | This study |
| TMB6257 | DK1042 *sacA*::$P_{liaI}$-*luxABCDE_cat*, *epeAB*::*spec*, *srfAB*::*kan* | This study |
| TMB6262 | DK1042 *sacA*::$P_{liaI}$-*luxABCDE_cat*, *epeAB*::*spec*, *comA*::*mls* | This study |
| TMB6274 | DK1042 *sacA*::$P_{liaI}$-*luxABCDE_cat*, *epeAB*::*spec*, comX::*kan* | This study |
| TMB6274 | DK1042 *sacA*::$P_{liaI}$-*luxABCDE_cat*, *epeAB*::*spec*, *comX*::*kan* | This study |
| TMB6284 | DK1042 *sacA*::$P_{liaI}$-*luxABCDE_cat*, *epeAB*::*spec*, *spo0A*::*kan* | This study |
| TMB6285 | DK1042 *sacA*::$P_{liaI}$-*luxABCDE_cat*, *epeAB*::*spec*, *spo0E*::*mls* | This study |
| TMB6286 | DK1042 *sacA*::$P_{liaI}$-*luxABCDE_cat*, *epeAB*::*spec*, *spo0F*::*mls* | This study |
| TMB6334 | DK1042 *sacA*::$P_{liaI}$-*luxABCDE_cat*, *epeAB*::*spec*, *comK*::*mls* | This study |
| TMB6350 | DK1042 *sacA*::$P_{empty}$-*luxABCDE_cat* | This study |
| TMB6382 | DK1042 *sacA*::$P_{srfAA}$-*luxABCDE_cat* | This study |
| TMB6687 | DK1042 *kre*::*mls* | This study |
| TMB6688 | DK1042 *spoVG*::*mls* | This study |
| TMB6692 | DK1042 *sacA*::$P_{liaI}$-*luxABCDE_cat*, *epeAB*::*spec*, *spoVG*::*mls* | This study |
| TMB6752 | DK1042 *sacA*::$P_{liaI}$-*luxABCDE_cat*, *epeAB*::*spec*, Δ126 | This study |
| TMB6755 | DK1042 *sacA*::$P_{liaI}$-*luxABCDE_cat*, *epeAB*::*spec*, *kre*::*kan* | This study |
| TMB6767 | DK1042 *sacA*::$P_{liaI}$-*luxABCDE_cat*, *epeAB*::*spec*, *abrB*::*kan* | This study |
| TMB6789 | DK1042 *sacA*::$P_{liaI}$-*luxABCDE_cat*, *epeAB*::*spec*, *comS*::*mls* | This study |
| TMB6831 | DK1042 *sacA*::$P_{epeX}$-*luxABCDE_cat*, *spoVG*::*mls* | This study |
| TMB6833 | DK1042 *sacA*::$P_{epeX}$-*luxABCDE_cat*, *kre*::*mls* | This study |
| TMB6839 | DK1042 *sacA*::$P_{abrB}$-*luxABCDE_cat* | This study |
| TMB6865 | DK1042 *sacA*::$P_{epeX\_}$*epeX*_IGR-*luxABCDE_cat*, | This study |
| TMB6892 | DK1042 *sacA*::$P_{epeX\_}$*epeX*_IGR-*luxABCDE_cat*, *spoVG*::*mls* | This study |
| TMB6893 | DK1042 *sacA*::$P_{epeX\_}$*epeX*_IGR-*luxABCDE_cat*, *comK*::*mls* | This study |
| TMB6894 | DK1042 *sacA*::$P_{epeX\_}$*epeX*_IGR-*luxABCDE_cat*, *kre*::*mls* | This study |
| TMB6905 | DK1042 *sacA*::$P_{epeX\_}$*epeX*_IGR-*luxABCDE_cat*, *abrB*::*mls* | This study |
| TMB6906 | DK1042 *sacA*::$P_{epeX\_}$*epeX*_IGR-*luxABCDE_cat*, *spo0A*::*kan* | This study |
| TMB6927 | DK1042 *sacA*::$P_{comK}$-*luxABCDE_cat* | This study |
| TMB6986 | DK1042 *sacA*::$P_{epeX}$-*luxABCDE_cat*, *spo0A*::*kan* | This study |

| | | |
|---|---|---|
| TMB6988 | DK1042 *sacA*::P$_{epeX}$_*epeX*_IGR-*luxABCDE_cat, spo0A*::*kan, abrB*::*mls* | This study |
| TMB6989 | DK1042 *sacA*::P$_{epeX}$-*luxABCDE_cat, abrB*::*mls* | This study |
| TMB6996 | DK1042 *sacA*::P$_{epeX}$-*luxABCDE_cat, spo0A*::*kan*, *abrB*::*mls* | This study |
| TMB7105 | DK1042 *sacA*::*epeX*_IGR-*luxABCDE_cat* | This study |
| TMB7118 | DK1042 *sacA*::P$_{spoIVA}$-*luxABCDE_cat* | This study |
| TMB7140 | DK1042 *sacA*::P$_{liaI}$-*luxABCDE_cat*, *epeXEPAB*::*spec* | This study |
| TMB7172 | DK1042 *sacA*::P$_{spoVG}$-*luxABCDE_cat* | This study |
| TMB7173 | DK1042 *sacA*::P$_{kre}$-*luxABCDE_cat* | This study |
| TMB7305 | DK1042 *sacA*::P$_{epeX}$-*luxABCDE_cat, comK*::*mls* | This study |
| TMB7306 | DK1042 *sacA*::P$_{liaI}$-*luxABCDE_cat*, *epeAB*::*spec, comK*::*mls, kre*::*kan* | This study |
| TMB7318 | DK1042 *sacA*::P$_{liaI}$-*luxABCDE_cat*, *epeAB*::*spec, amyE*::P$_{spoVG}$-*spoVG_kan, spoVG*::*mls* | This study |
| TMB7319 | DK1042 *sacA*::P$_{liaI}$-*luxABCDE_cat*, *epeAB*::*spec*, Δ*spoVG* | This study |
| TMB7331 | DK1042 *sacA*::P$_{liaI}$-*luxABCDE_cat*, *epeAB*::*spec, comK*::*mls,* Δ*spoVG* | This study |
| TMB7332 | DK1042 *sacA*::P$_{liaI}$-*luxABCDE_cat*, *epeAB*::*spec, kre*::*kan,* Δ*spoVG* | This study |
| TMB7333 | DK1042 *sacA*::P$_{liaI}$-*luxABCDE_cat*, *epeAB*::*spec, comK*::*mls, kre*::*kan,* Δ*spoVG* | This study |
| TME3984 | DH10*β* pBS3C*lux*_P$_{comK}$ | This study |
| TME4430 | DH10*β* pBS3C*lux*_P$_{spo0A}$ | This study |
| TME4510 | DH10*β* pBS3C*lux*_P$_{srfAA}$ | This study |
| TME4747 | DH10*β* pBS3C*αlux*_P$_{epeX}$_*epeX*_IGR | This study |
| TME4758 | DH10*β* pBS3C*lux*_P$_{abrB}$ | This study |
| TME4831 | DH10*β* pBS3C*lux*_P$_{spoIVA}$ | This study |
| TME4868 | DH10*β* pBS3C*αlux*_P$_{spoVG}$ | This study |
| TME4869 | DH10*β* pBS3C*αlux*_P$_{kre}$ | This study |
| TME4870 | DH10*β* pET28b_*spoVG*_C-His$_6$-tag | This study |
| TME4871 | DH10*β* pET28b_*kre*_C-His$_6$-tag | This study |
| TME4894 | BL21 pET28b_*spoVG*_C-His$_6$-tag | This study |
| TME4894 | BL21 pET28b_*kre*_C-His$_6$-tag | This study |
| TME4970 | DH10*β* pBS1K_P$_{spoVG}$-*spoVG* | This study |
| TME4972 | DH10*β* pMAD_*spoVG* clean deletion | This study |

* cat=chloramphenicol resistance; bla=ampicillin resistance; mls=macrolide-lincosamide-streptogramin B resistance; kan=kanamycin resistance

**Table S2**: Plasmids used in this study.

| Plasmids | Genotype* | Reference |
|---|---|---|
| pBS1K | *amyE' … 'amyE, bla, kan* | Popp et al. 2017 (2) |
| pBS1K_P$_{spoVG}$-*spoVG* | *amyE'* P$_{spoVG}$-*spoVG_kan 'amyE, bla,* | This study |
| pBS3C*lux* | *sacA'* … *'sacA, luxABCDE, bla, cat* | Radeck et al. 2013 (3) |
| pBS3C*lux*_P$_{abrB}$ | *sacA'* P$_{abrB}$-*luxABCDE_cat 'sacA* | This study |
| pBS3C*lux*_P$_{comK}$ | *sacA'* P$_{comK}$-*luxABCDE_cat 'sacA* | This study |
| pBS3C*lux*_P$_{empty}$ | *sacA' luxABCDE_cat 'sacA* | Laboratory stock |
| pBS3C*lux*_P$_{epeX}$ | *sacA'* P$_{epeX}$-*luxABCDE_cat 'sacA* | Popp et al. 2021 (4) |
| pBS3C*lux*_P$_{liaI}$ | *sacA'* P$_{liaI}$-*luxABCDE_cat 'sacA* | Radeck et al. 2016 (5) |
| pBS3C*lux*_P$_{spo0A}$ | *sacA'* P$_{liaI}$-*luxABCDE_cat 'sacA* | This study |
| pBS3C*αlux* | *sacA'* … *'sacA, luxABCDE, bla, cat* (exchangeable RBS-site of *luxA*), *lacZα* | Popp et al. 2017 (2) |
| pBS3C*αlux*_P$_{epeX}$-*epeX*-IGR | *sacA'* P$_{epeX}$-*epeX*-IGR-*luxABCDE_cat 'sacA* | This study |
| pBS3C*αlux*_P$_{kre}$ | *sacA'* P$_{kre}$-*luxABCDE_cat 'sacA* | This study |
| pBS3C*αlux*_P$_{spoVG}$ | *sacA'* P$_{spoVG}$-*luxABCDE_cat 'sacA* | This study |
| pET28b(+) | protein expression vector | Laboratory stock |
| pET28b(+)_*kre*_C-6x- His$_6$-tag | *kre*_C-6x- His$_6$-tag | This study |
| pET28b(+)_*spoVG*_C-His$_6$-tag | *spoVG*_C-His$_6$-tag | This study |
| pMAD | ori pE194-temperature sensitive, *bla, mls* | Arnaud et al. 2004 (6) |
| pMAD_*spoVG* clean deletion | ori pE194-temperature sensitive, *bla, mls, spoVG* flanking sites | This study |

* cat=chloramphenicol resistance; bla=ampicillin resistance; mls=macrolide-lincosamide-streptogramin B resistance; kan=kanamycin resistance

**Table S3**: Oligonucleotides used in this study.

| Primer | Sequence 5'-3' | Source/ Name* |
|---|---|---|
| TM0137 | CAGCGAACCATTTGAGGTGATAGG | *kan* fwd |
| TM0138 | CGATACAAATTCCTCGTAGGCGCTCGG | *kan* rev |
| TM0149 | CGTATGTATTCAAATATATCCTCCTCAC | *spec* check rev |
| TM0716 | CAGGAAAAGGCCATTTTACC | *mls* check fwd |
| TM0717 | AAATCGTCAATTCCTGCATGT | *mls* check rev |
| TM2262 | GAGCGTAGCGAAAAATCC | pBS3C*lux* check fwd |
| TM2263 | GAAATGATGCTCCAGTAACC | pBS3C*lux* check rev |
| TM2759 | GAGGGGCTAGAGGACTATAGG | *epeX* check fwd |

| | | |
|---|---|---|
| TM2778 | gatcgaattcgcggccgcttctagagTGCCATTGCATCTCCTGCGACTGTCG | $P_{abrB}$ fwd |
| TM2779 | gatcactagtaGAGATACTTATTTGTTTAAATTATATTTTTCTTCG | $P_{abrB}$ rev |
| TM6452 | GACTTCATGAGCTCAGTGTAC | *comK* check fwd |
| TM6805 | ACGAAAAACCTGCTGTCCTTT | *comX* check fwd |
| TM6806 | GTAATGACGCCGAAGCTAATG | *comX* check rev |
| TM7270 | gtcaggtctcactagtaTTCTAATATAAACCAATTATTCTATATCATTCAATACATGTC | $P_{epeX}$ rev |
| TM7367 | GAGGTCTAATGCAAGCTGTAATTC | *srfAA* check fwd |
| TM7368 | ATCTCATAGAGCGGCACGTAATC | *srfAA* check rev |
| TM7369 | AGATCACTTGCTTCTTGCCATTC | *srfAB* check fwd |
| TM7370 | TTCTGCTTGTATCCCTTGTAAGC | *srfAB* check rev |
| TM7373 | TAGATAAGCATGAAGTTGGACCG | *comA* check fwd |
| TM7374 | GGTGATGTTTACGCTGATCCTG | *comA* check rev |
| TM7405 | GTGTTCTTGTCGTCGGATTTC | *spo0A* check fwd |
| TM7406 | TCGAAATTCACGGTGCAAAC | *spo0A* check rev |
| TM7407 | GGCAACAATCAGAAGGTCTCA | *spo0B* check fwd |
| TM7408 | GATTGGCAACCTCAAACAGAA | *spo0B* check rev |
| TM7409 | GAAGTGTATCAGGATGAAGTGGTG | *spo0F* check fwd |
| TM7410 | GGTTTGAAGAACCAAATTCACG | *spo0F* check rev |
| TM7413 | GGACAGTCAAGAGGTTATTTATGAGC | *spo0E* check fwd |
| TM7414 | AGTGAGTGAGTTTCTCGTTCTCG | *spo0E* check rev |
| TM7415 | ATAGTATTTCAGAAGACGATCCGC | *abrB* check fwd |
| TM7416 | AATGTAAGGACAATAGCTGGTATGC | *abrB* check rev |
| TM7512 | agtcactagtGTGCAACGCATTTTCTCTTTC | $P_{srfAA}$ fwd |

| | | |
|---|---|---|
| TM7514 | agtcggtctcaaattTCTTGAAGCCATGTATGAGTG | $P_{srfAA}$ rev |
| TM7811 | cgcgaattcAGTATATGGATAACGGTCGA | $P_{comK}$ fwd |
| TM7812 | atgctagcGCCTCCATCCTTTTTCTGCA | $P_{comK}$ rev |
| TM7813 | gaattcCGTTTTTTTGTGCCAATGGGTC | $P_{liaI}$ fwd |
| TM7814 | gtcgacTCGTTTTCCTTGTCTTCATCTTATAC | $P_{liaI}$ rev |
| TM7854 | AGTTTGGTACAAGCCATTCTGC | *kre* check fwd |
| TM7855 | CACTGTTAGTTGGAAAGTCAATGG | *kre* check rev |
| TM7856 | AGCGCGAAATTGATGTTGTC | *spoVG* check fwd |
| TM7857 | AGGGCCAATAATCGTATCTTCTC | *spoVG* check rev |
| TM7977 | CACCTGATTCAAATGGACAGCCTG | *comS* check fwd |
| TM7978 | GCATATCAATGAGCAGCAGGTGG | *comS* check rev |
| TM7985 | acgaattcATGAAAAAGGAAATCACTAAC | $P_{epeX}$_*epeX*_IGR rev |
| TM8135 | agtcggtctcgaattGGACCTTTATATGATAGTACCGC | $P_{epeX}$ fwd |
| TM8147 | agtcggtctcgaattGGACCTTTATATGATAGTACCGC | $P_{epeX}$_*epeX*_IGR fwd |
| TM8186 | atcgctgcagCAATCAAACACAGGACACTA | $P_{spoIVA}$ fwd |
| TM8187 | agtcgaattcTCCCCGCGTTTTCTGTAAA | $P_{spoIVA}$ rev |
| TM8211 | agtcggtctcgaattATGAAAAAGGAAATCACTAACAATG | *epeX*_IGR fwd |
| TM8212 | agtcggtctcgtcgaGAAATATCCCCCAACAAATG | *epeX*_IGR rev |
| TM8290 | ctaggaattcGAAAAAGAAGGCAAGATCTGTATAATG | $P_{kre}$ fwd |
| TM8291 | ctaggctagcCTCTCCTTTGCTCTTATCAATTTCTATATG | $P_{kre}$ rev |
| TM8292 | ctaggaattcGAAAAGTGATTCTGGGAGAGC | $P_{spoVG}$ fwd |
| TM8293 | ctaggctagcGTTCACCACCTTTTCCCTATA | $P_{spoVG}$ rev |
| TM8296 | ctagggtctcgcatgGTGGAAGTTACTGACGTAAGATTACG | *spoVG* C-terminal fwd |
| TM8297 | ctagggtctcgaattAGAAGCTCCAGCTTCTTCGAAT | *spoVG* C-terminal rev |
| TM8300 | ctagggtctcgcatgATGGACGACCATGCATATACG | *kre* C-terminal fwd |
| TM8301 | ctagggtctcgaattAAAGTAACTCTCGCCAAGTTTTTTAAAG | *kre* C-terminal rev |

| | | |
|---|---|---|
| TM8412 | GATTTCCTTTTTCATATTATCCCTCCTCCTTTTCTAATAT | *in vitro epeX*_1 rev |
| TM8413 | gaaattaatacgactcactatagggagaATATTAGAAAAGGAGGAGGGATAATATGAAAAAGGAAATC | *in vitro epeX*_1 _T7 fwd |
| TM8414 | AGTTTTTCACAGTCTCATTGTTAGTGATTTCCTTTTTCAT | *in vitro epeX*_2 rev |
| TM8415 | gaaattaatacgactcactatagggagaATGAAAAAGGAAATCACTAACAATGAGACTGTGAAAAACT | *in vitro epeX*_2 _T7 fwd |
| TM8416 | TAATAGACCCTTAAATTCTAAGTTTTTCACAGTCTCATTG | *in vitro epeX*_3 rev |
| TM8417 | gaaattaatacgactcactatagggagaCAATGAGACTGTGAAAAACTTAGAATTTAAGGGTCTATTA | *in vitro epeX*_3 _T7 fwd |
| TM8418 | GCTAACTTTTGTGATTCATCTAATAGACCCTTAAATTCTA | *in vitro epeX*_4 rev |
| TM8419 | gaaattaatacgactcactatagggagaTAGAATTTAAGGGTCTATTAGATGAATCACAAAAGTTAGC | *in vitro epeX*_4 _T7 fwd |
| TM8420 | ACCAAAGATCATTCACTTTTGCTAACTTTTGTGATTCATC | *in vitro epeX*_5 rev |
| TM8421 | gaaattaatacgactcactatagggagaGATGAATCACAAAAGTTAGCAAAAGTGAATGATCTTTGGT | *in vitro epeX*_5_T7 fwd |
| TM8422 | TTCTTTTGATTTTACAAAATACCAAAGATCATTCACTTTT | *in vitro epeX*_6 rev |
| TM8423 | gaaattaatacgactcactatagggagaAAAAGTGAATGATCTTTGGTATTTTGTAAAATCAAAAGAA | *in vitro epeX*_6_T7 fwd |
| TM8424 | CTTCCAAGAATCCAGCGATTTTCTTTTGATTTTACAAAAT | *in vitro epeX*_7 rev |
| TM8425 | gaaattaatacgactcactatagggagaATTTTGTAAAATCAAAAGAAAATCGCTGGATTCTTGGAAG | *in vitro epeX*_7_T7 fwd |
| TM8426 | ACCCCTCTAATTAATGACCACTTCCAAGAATCCAGCGATT | *in vitro epeX*_8 rev |
| TM8427 | gaaattaatacgactcactatagggagaAATCGCTGGATTCTTGGAAGTGGTCATTAATTAGAGGGGT | *in vitro epeX*_8_T7 fwd |
| TM8428 | AGAGTTAGACTCTCCTTTGTACCCCTCTAATTAATGACCA | *in vitro epeX*_9 rev |

| | | |
|---|---|---|
| TM8429 | gaaattaatacgactcactatagggagaTGGTCATTAATTAGAGGGGTACAAAGGAGAGTCTAACTCT | *in vitro* *epeX*_9_T7 fwd |
| TM8430 | CCCCCAACAAATGGTAAAGGAGAGTTAGACTCTCCTTTGT | *in vitro* *epeX*_10 rev |
| TM8431 | gaaattaatacgactcactatagggagaACAAAGGAGAGTCTAACTCTCCTTTACCATTTGTTGGGGG | *in vitro* *epeX*_10_T7 fwd |
| TM8432 | TTTTATTATACATGAAATATCCCCCAACAAATGGTAAAGG | *in vitro* *epeX*_11 rev |
| TM8433 | gaaattaatacgactcactatagggagaCCTTTACCATTTGTTGGGGGATATTTCATGTATAATAAAA | *in vitro* *epeX*_11_T7 fwd |
| TM8442 | agtcggtctcaaattGGATGCGTTAGTCGAGATCG | *spoVG* complement fwd |
| TM8443 | agtcggtctcactagGGAGACTTCATGAAGTGCAAG | *spoVG* complement rev |
| TM8446 | agtcggtctcgtcgaGATGTTGTCATGACCGTTGCC | *spoVG* upstream fwd |
| TM8447 | agtcggtctcgacctAGTAGTTCACCACCTTTTCCC | *spoVG* upstream rev |
| TM8448 | agtcggtctcgaggtCCAAAAAGCAAGGACTGCTGAAAG | *spoVG* downstream fwd |
| TM8449 | agtcggtctcgaattCCACTTTACTGTGATTGACTACCG | *spoVG* downstream rev |
| TM8470 | CCCCGACTACCATCGGCGCTGA | *in vitro* 5s rev |
| TM8471 | gaaattaatacgactcactatagggagaGGTGGCGATAGCGAAGAGGTCAC | *in vitro* 5s_T7 fwd |

* mls=macrolide-lincosamide-streptogramin B resistance; kan=kanamycin resistance, spec=spectinomycin resistance; fwd=forward; rev=reverse

**Table S4**: Transition list of mass spectrometry analysis.

| Sample | Protein Name | Peptide Sequence | Fragment Ion | m/z Precursor Ion (Q1) | m/z Fragment Ion (Q3) | Collision Energy |
|---|---|---|---|---|---|---|
| LysC | EpeE | TIQVISFTGGEVFLDYK | y4 | 639.67 | 538.29 | 26.6 |
| LysC | EpeE | TIQVISFTGGEVFLDYK | y3 | 639.67 | 425.20 | 26.6 |
| LysC | EpeE | TIQVISFTGGEVFLDYK | y9 | 639.67 | 514.26 | 26.6 |
| LysC | EpeE | TIQVISFTGGEVFLDYK | y7 | 639.67 | 457.24 | 26.6 |
| LysC | EpeE | TIQVISFTGGEVFLDYK | y5 | 639.67 | 343.18 | 26.6 |
| LysC | EpeE | QITLISNGFWGLSK | y4 | 521.96 | 404.25 | 22.1 |
| LysC | EpeE | QITLISNGFWGLSK | y3 | 521.96 | 347.23 | 22.1 |
| LysC | EpeE | QITLISNGFWGLSK | y9 | 521.96 | 498.25 | 22.1 |
| LysC | EpeE | QITLISNGFWGLSK | y7 | 521.96 | 397.71 | 22.1 |
| LysC | EpeE | QITLISNGFWGLSK | y6 | 521.96 | 369.20 | 22.1 |
| LysC | EpeX | LAKVNDLWYFVK | y6 | 499.28 | 855.48 | 21.3 |
| LysC | EpeX | LAKVNDLWYFVK | y4 | 499.28 | 556.31 | 21.3 |
| LysC | EpeX | LAKVNDLWYFVK | y3 | 499.28 | 393.25 | 21.3 |
| LysC | EpeX | LAKVNDLWYFVK | y6 | 499.28 | 428.24 | 21.3 |
| LysC | EpeX | VNDLWYFVK | y7 | 592.31 | 970.50 | 20.7 |
| LysC | EpeX | VNDLWYFVK | y3 | 592.31 | 393.25 | 20.7 |
| LysC | EpeX | VNDLWYFVK | y7 | 592.31 | 485.76 | 20.7 |
| LysC | EpeX | VNDLWYFVK | y6 | 592.31 | 428.24 | 20.7 |
| LysC | EpeX | VNDLWYFVK | y5 | 592.31 | 371.70 | 20.7 |
| LysC | EpeX | SKENRWILGSGH | y6 | 692.36 | 583.32 | 23.7 |
| LysC | EpeX | SKENRWILGSGH | y5 | 692.36 | 470.24 | 23.7 |
| LysC | EpeX | SKENRWILGSGH | y4 | 692.36 | 357.15 | 23.7 |
| LysC | EpeX | SKENRWILGSGH | y3 | 692.36 | 300.13 | 23.7 |
| LysC | EpeX | SKENRWILGSGH | y7 | 692.36 | 385.20 | 23.7 |
| LysC | EpeX | SKENRWILGSGH | y7 | 461.91 | 769.40 | 19.8 |
| LysC | EpeX | SKENRWILGSGH | y3 | 461.91 | 300.13 | 19.8 |
| LysC | EpeX | SKENRWILGSGH | y11 | 461.91 | 648.84 | 19.8 |
| LysC | EpeX | SKENRWILGSGH | y9 | 461.91 | 520.28 | 19.8 |
| Trypsin | Kre | MDDHAYTK | y6 | 490.71 | 734.35 | 17.6 |
| Trypsin | Kre | MDDHAYTK | y4 | 490.71 | 482.26 | 17.6 |
| Trypsin | Kre | MDDHAYTK | y3 | 490.71 | 411.22 | 17.6 |
| Trypsin | Kre | MDDHAYTK | y7 | 490.71 | 425.19 | 17.6 |
| Trypsin | Kre | AKHLLQEYVGMK | y8 | 708.89 | 967.49 | 24.2 |
| Trypsin | Kre | AKHLLQEYVGMK | y7 | 708.89 | 854.41 | 24.2 |
| Trypsin | Kre | AKHLLQEYVGMK | y6 | 708.89 | 726.35 | 24.2 |
| Trypsin | Kre | AKHLLQEYVGMK | y5 | 708.89 | 597.31 | 24.2 |
| Trypsin | Kre | AKHLLQEYVGMK | y8 | 708.89 | 484.25 | 24.2 |
| Trypsin | Kre | EKPLVPNR | y7 | 476.78 | 823.51 | 17.2 |
| Trypsin | Kre | EKPLVPNR | y6 | 476.78 | 695.42 | 17.2 |
| Trypsin | Kre | EKPLVPNR | y3 | 476.78 | 386.21 | 17.2 |

| | | | | | | |
|---|---|---|---|---|---|---|
| Trypsin | Kre | EKPLVPNR | y7 | 476.78 | 412.26 | 17.2 |
| Trypsin | Kre | EKPLVPNR | y6 | 476.78 | 348.21 | 17.2 |
| Trypsin | Kre | QQPAYHKPVFK | y5 | 448.25 | 618.40 | 19.3 |
| Trypsin | Kre | QQPAYHKPVFK | y4 | 448.25 | 490.30 | 19.3 |
| Trypsin | Kre | QQPAYHKPVFK | y8 | 448.25 | 495.28 | 19.3 |
| Trypsin | Kre | QQPAYHKPVFK | y7 | 448.25 | 459.76 | 19.3 |
| Trypsin | Kre | QQPAYHKPVFK | y6 | 448.25 | 378.23 | 19.3 |
| Trypsin | EpeE | EYIRELVTEFAK | y6 | 499.94 | 694.38 | 21.3 |
| Trypsin | EpeE | EYIRELVTEFAK | y5 | 499.94 | 595.31 | 21.3 |
| Trypsin | EpeE | EYIRELVTEFAK | y10 | 499.94 | 603.35 | 21.3 |
| Trypsin | EpeE | EYIRELVTEFAK | y7 | 499.94 | 404.23 | 21.3 |
| Trypsin | EpeE | EYIRELVTEFAK | y6 | 499.94 | 347.69 | 21.3 |
| Trypsin | EpeE | ELVTEFAK | y6 | 468.76 | 694.38 | 17.0 |
| Trypsin | EpeE | ELVTEFAK | y5 | 468.76 | 595.31 | 17.0 |
| Trypsin | EpeE | ELVTEFAK | y4 | 468.76 | 494.26 | 17.0 |
| Trypsin | EpeE | ELVTEFAK | y3 | 468.76 | 365.22 | 17.0 |
| Trypsin | EpeE | ELVTEFAK | y6 | 468.76 | 347.69 | 17.0 |
| Trypsin | EpeE | TIQVISFTGGEVFLDYK | y7 | 959.00 | 913.47 | 31.7 |
| Trypsin | EpeE | TIQVISFTGGEVFLDYK | y6 | 959.00 | 784.42 | 31.7 |
| Trypsin | EpeE | TIQVISFTGGEVFLDYK | y11 | 959.00 | 638.32 | 31.7 |
| Trypsin | EpeE | TIQVISFTGGEVFLDYK | y7 | 959.00 | 457.24 | 31.7 |
| Trypsin | EpeE | TIQVISFTGGEVFLDYK | y5 | 959.00 | 343.18 | 31.7 |
| Trypsin | EpeE | QITLISNGFWGLSK | y4 | 521.96 | 404.25 | 22.1 |
| Trypsin | EpeE | QITLISNGFWGLSK | y3 | 521.96 | 347.23 | 22.1 |
| Trypsin | EpeE | QITLISNGFWGLSK | y9 | 521.96 | 498.25 | 22.1 |
| Trypsin | EpeE | QITLISNGFWGLSK | y7 | 521.96 | 397.71 | 22.1 |
| Trypsin | EpeE | QITLISNGFWGLSK | y6 | 521.96 | 369.20 | 22.1 |
| Trypsin | EpeE | LNSNLLLFILRK | y5 | 722.46 | 676.45 | 24.6 |
| Trypsin | EpeE | LNSNLLLFILRK | y4 | 722.46 | 529.38 | 24.6 |
| Trypsin | EpeE | LNSNLLLFILRK | y10 | 722.46 | 608.89 | 24.6 |
| Trypsin | EpeE | LNSNLLLFILRK | y8 | 722.46 | 508.35 | 24.6 |
| Trypsin | EpeE | LNSNLLLFILRK | y6 | 722.46 | 395.27 | 24.6 |
| Trypsin | EpeE | EGFKWFLNILK | y5 | 465.60 | 600.41 | 20.0 |
| Trypsin | EpeE | EGFKWFLNILK | y4 | 465.60 | 487.32 | 20.0 |
| Trypsin | EpeE | EGFKWFLNILK | y3 | 465.60 | 373.28 | 20.0 |
| Trypsin | EpeE | EGFKWFLNILK | y8 | 465.60 | 531.33 | 20.0 |
| Trypsin | EpeE | EGFKWFLNILK | y6 | 465.60 | 374.24 | 20.0 |
| Trypsin | EpeX | WILGSGH | y6 | 385.20 | 583.32 | 14.5 |
| Trypsin | EpeX | WILGSGH | y5 | 385.20 | 470.24 | 14.5 |
| Trypsin | EpeX | WILGSGH | y4 | 385.20 | 357.15 | 14.5 |
| Trypsin | EpeX | WILGSGH | y3 | 385.20 | 300.13 | 14.5 |
| Trypsin | SpoVG | VEVTDVR | y6 | 409.22 | 718.37 | 15.2 |
| Trypsin | SpoVG | VEVTDVR | y5 | 409.22 | 589.33 | 15.2 |

| | | | | | | |
|---|---|---|---|---|---|---|
| Trypsin | SpoVG | VEVTDVR | y4 | 409.22 | 490.26 | 15.2 |
| Trypsin | SpoVG | VEVTDVR | y3 | 409.22 | 389.21 | 15.2 |
| Trypsin | SpoVG | VEVTDVR | y6 | 409.22 | 359.69 | 15.2 |
| Trypsin | SpoVG | TPDGEFR | y6 | 411.19 | 720.33 | 15.2 |
| Trypsin | SpoVG | TPDGEFR | y5 | 411.19 | 623.28 | 15.2 |
| Trypsin | SpoVG | TPDGEFR | y4 | 411.19 | 508.25 | 15.2 |
| Trypsin | SpoVG | TPDGEFR | y3 | 411.19 | 451.23 | 15.2 |
| Trypsin | SpoVG | TPDGEFR | y6 | 411.19 | 360.67 | 15.2 |
| Trypsin | SpoVG | TPDGEFRDITHPINSSTR | y5 | 681.67 | 564.27 | 28.2 |
| Trypsin | SpoVG | TPDGEFRDITHPINSSTR | y4 | 681.67 | 450.23 | 28.2 |
| Trypsin | SpoVG | TPDGEFRDITHPINSSTR | y3 | 681.67 | 363.20 | 28.2 |
| Trypsin | SpoVG | TPDGEFRDITHPINSSTR | y8 | 681.67 | 456.24 | 28.2 |
| Trypsin | SpoVG | TPDGEFRDITHPINSSTR | y6 | 681.67 | 339.18 | 28.2 |
| Trypsin | SpoVG | LGDTEALEFEEAGAS | y8 | 769.85 | 839.34 | 26.0 |
| Trypsin | SpoVG | LGDTEALEFEEAGAS | y7 | 769.85 | 710.30 | 26.0 |
| Trypsin | SpoVG | LGDTEALEFEEAGAS | y5 | 769.85 | 434.19 | 26.0 |
| Trypsin | SpoVG | LGDTEALEFEEAGAS | y4 | 769.85 | 305.15 | 26.0 |
| Trypsin | SpoVG | LGDTEALEFEEAGAS | y8 | 769.85 | 420.17 | 26.0 |
| Trypsin | SpoVG | IQDAVLNEYHR | y6 | 453.23 | 831.41 | 19.5 |
| Trypsin | SpoVG | IQDAVLNEYHR | y5 | 453.23 | 718.33 | 19.5 |
| Trypsin | SpoVG | IQDAVLNEYHR | y4 | 453.23 | 604.28 | 19.5 |
| Trypsin | SpoVG | IQDAVLNEYHR | y3 | 453.23 | 475.24 | 19.5 |
| Trypsin | SpoVG | IQDAVLNEYHR | y9 | 453.23 | 558.78 | 19.5 |
| Trypsin | SpoVG | IQDAVLNEYHR | y6 | 453.23 | 416.21 | 19.5 |

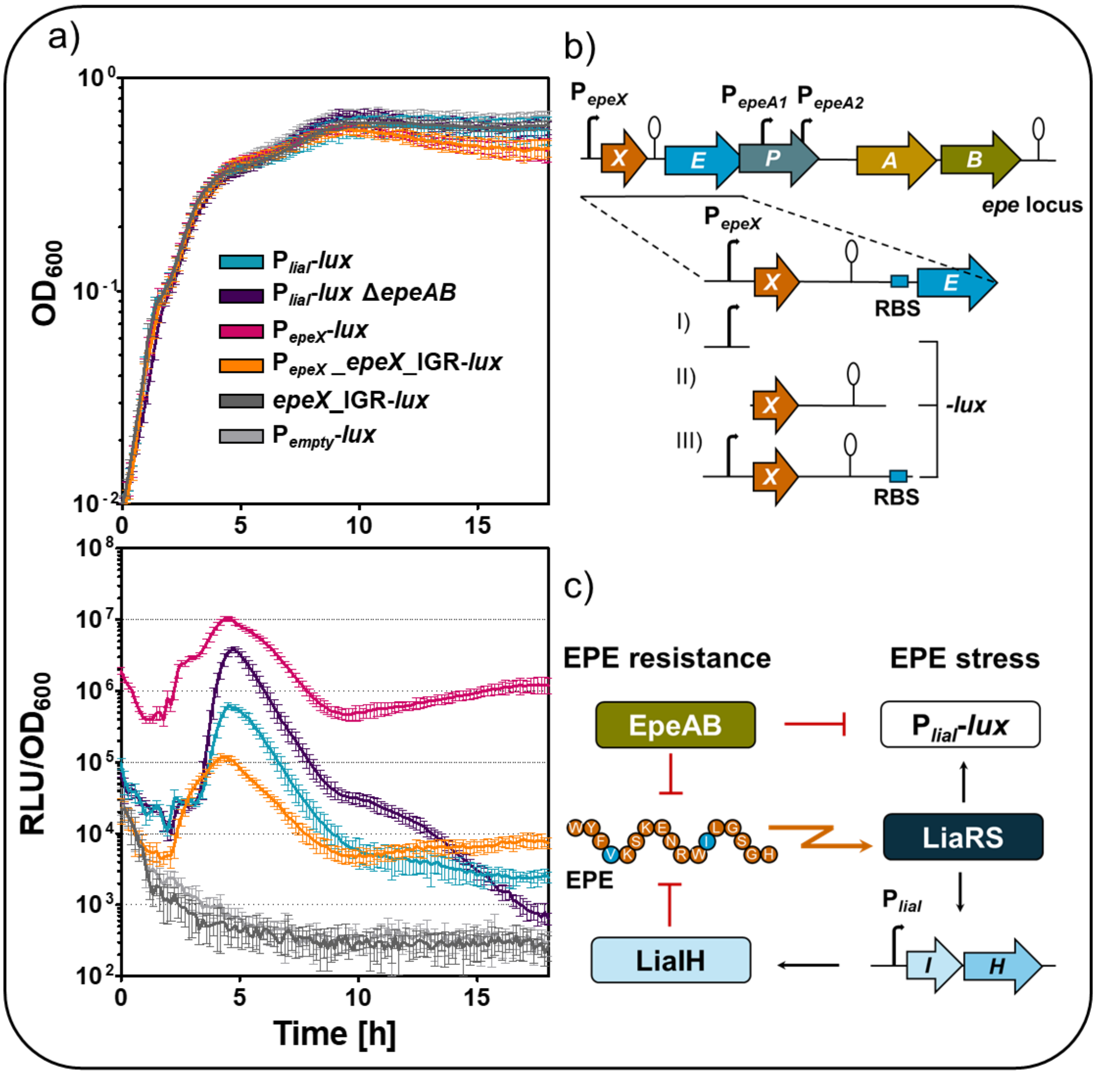


**Figure S1**: Timing of *epe* expression and EPE stress reponse.

**a)** The upper graph depicts the growth curve of decicated reporter strains as function of optical density at a wavelenght of 600 nm ($OD_{600}$) over time, and the lower graph shows the corresponding promoter activities as relative luminescence units (RLU) normalised to the $OD_{600}$. While EPE expression was monitored by applying $P_{epeX}$-*lux* (pink) and $P_{epeX}$-*epeX*_IGR-*lux* (orange) reporter fusion, EPE-mediated stress reponse was obtained by using the $P_{liaI}$-*lux* reporter (blue), which sensitivity can be increased by *epeAB* deletion (dark purple). $P_{empty}$-*lux* (light grey) and *epeX*_IGR-*lux* (dark grey) served as negative control, displaying the background luminescence activity. The standard error of the mean (SEM) was included at each point of measurement as indicated by error bars. **b)** Schematic overview of reporter constructs used to analyse dynamic of EPE expression. **I)** Trascriptional $P_{epeX}$-*lux* fusion was constructed to monitor *epeX* expression. **II)** The *epeX*-IGR-*lux* reporter does not contain the *epeX* promoter, thus the strains served as negative control. **III)** Further, the translational $P_{epeX}$-*epeX*_IGR-*lux* reporter fusion, was generated to monitor the dynamic of full *epeXEP* expression, necessary for EPE production. **c)** EPE specifically induces the *lia* system, whereby LiaS senses the EPE stress, leading to induction of $P_{liaI}$ in a LiaR-dependent manner. Induction of $P_{liaI}$, results in expression of LiaIH, which in turn confers, next to the intrinsic autoimmunity EpeAB, resistance against external EPE. The analysis was performed in DSM medium.

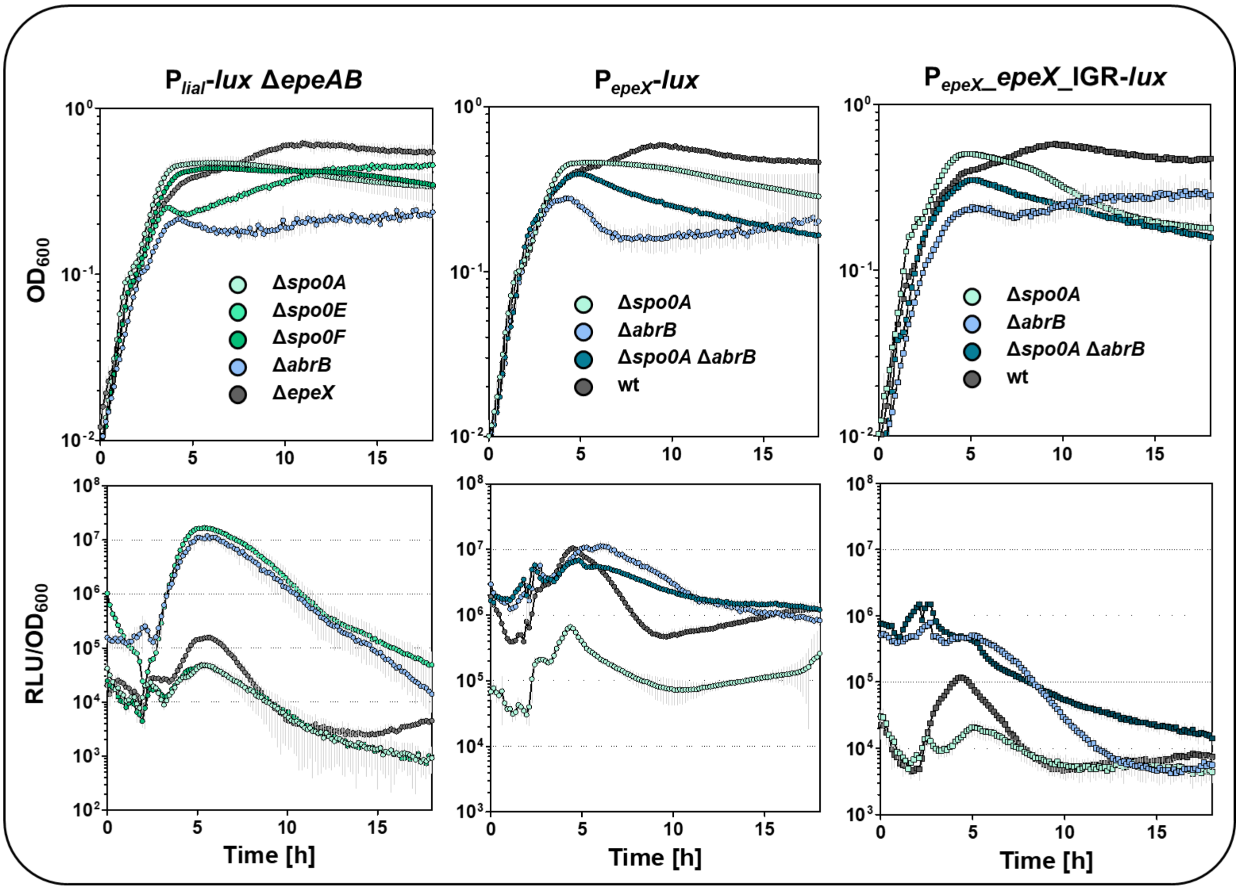


**Figure S2**: Impact of sporulation related mutants on EPE production and stress response.

While the upper graphs depict the growth curve as function of $OD_{600}$ over time, the lower graphs show the RLU values normalised to the corresponding $OD_{600}$ over time. The dynamic of the $P_{\mathit{liaI}}$-*lux epeAB*, $P_{\mathit{epeX}}$-*lux*, and $P_{\mathit{epeX}}$_*epeX*_IGR-*lux* reporter activity in absence and presence of sporulation associated gene deletions were displayed. The standard derivation of biological and technical triplicates was included as error bars to each time point of measurement. For simplification, gene deletions are indicated with the delta symbol (Δ) throughout the figure, although this does not necessarily imply clean deletion strains. The full genotypes and respective resistance cassettes are provided in Table S1.

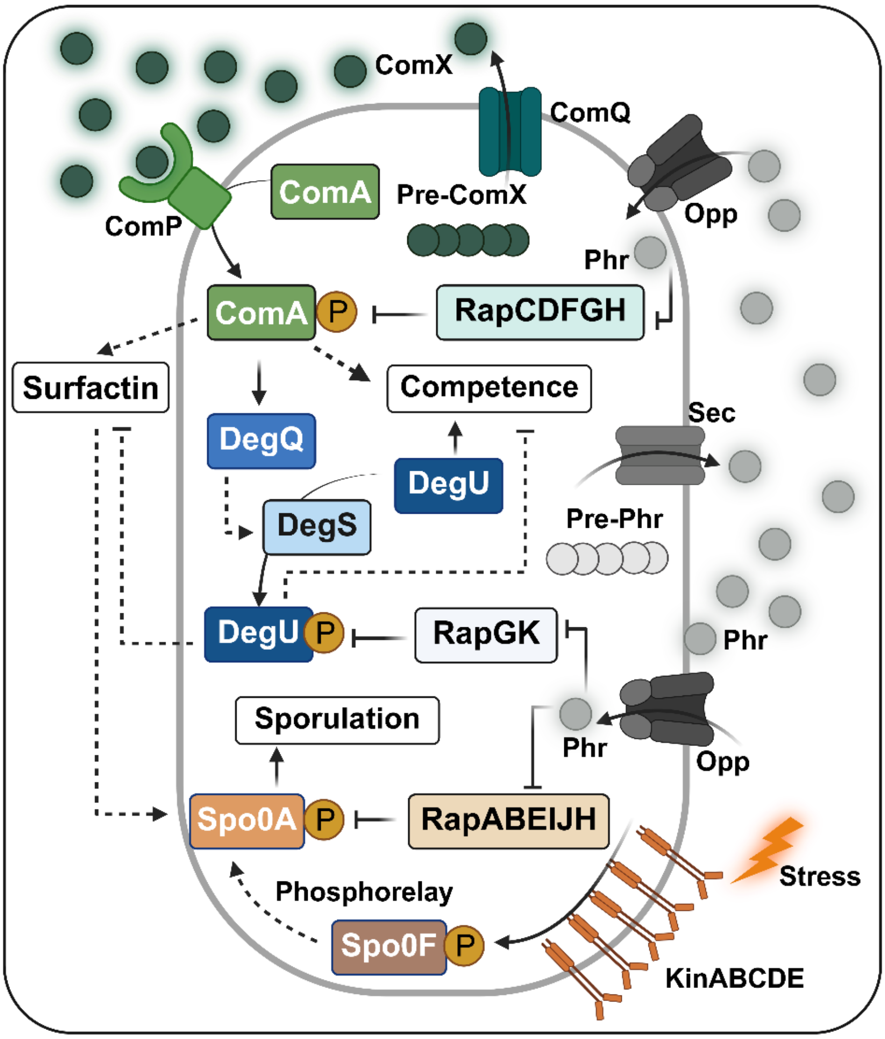


**Figure S3**: Quorum sensing networks and phosphorelay in *B. subtilis*.

Schematic representation of the regulatory pathways integrating quorum sensing signals, environmental stress, and phosphorelay to control among other competence and sporulation. Pre-ComX is synthesised intracellularly and exported by ComQ, where it is processed into mature ComX. Extracellular ComX accumulates in a density-dependent manner and is sensed by the membrane kinase ComP, which phosphorylates the response regulator ComA (ComA~P). ComA~P induces *srfAABCD* (*srf*) expression, leading to surfactin production and competence development. ComA~P also activates degQ, which stimulates DegS kinase activity and promotes DegU phosphorylation. While unphosphorylated DegU enhances *comK* transcription and competence, phosphorylated DegU represses *srf* expression, antagonising surfactin synthesis and competence. ComA~P activity is inhibited by RapC, D, F, G, and H, whereas DegU phosphorylation is counteracted by RapG and RapK. Rap phosphatase activity is relieved by cognate Phr peptides, which are secreted via the secretion (Sec) pathway, accumulate extracellularly, and are re-imported through the oligopeptide permease (Opp) once threshold concentrations are reached. Moreover, environmental stress signals are perceived by the sensor kinases KinA–E, which initiate the Spo0F–Spo0B–Spo0A phosphorelay, leading to Spo0A phosphorylation and ultimately sporulation. Surfactin further feeds into this pathway by stimulating Spo0A~P accumulation.

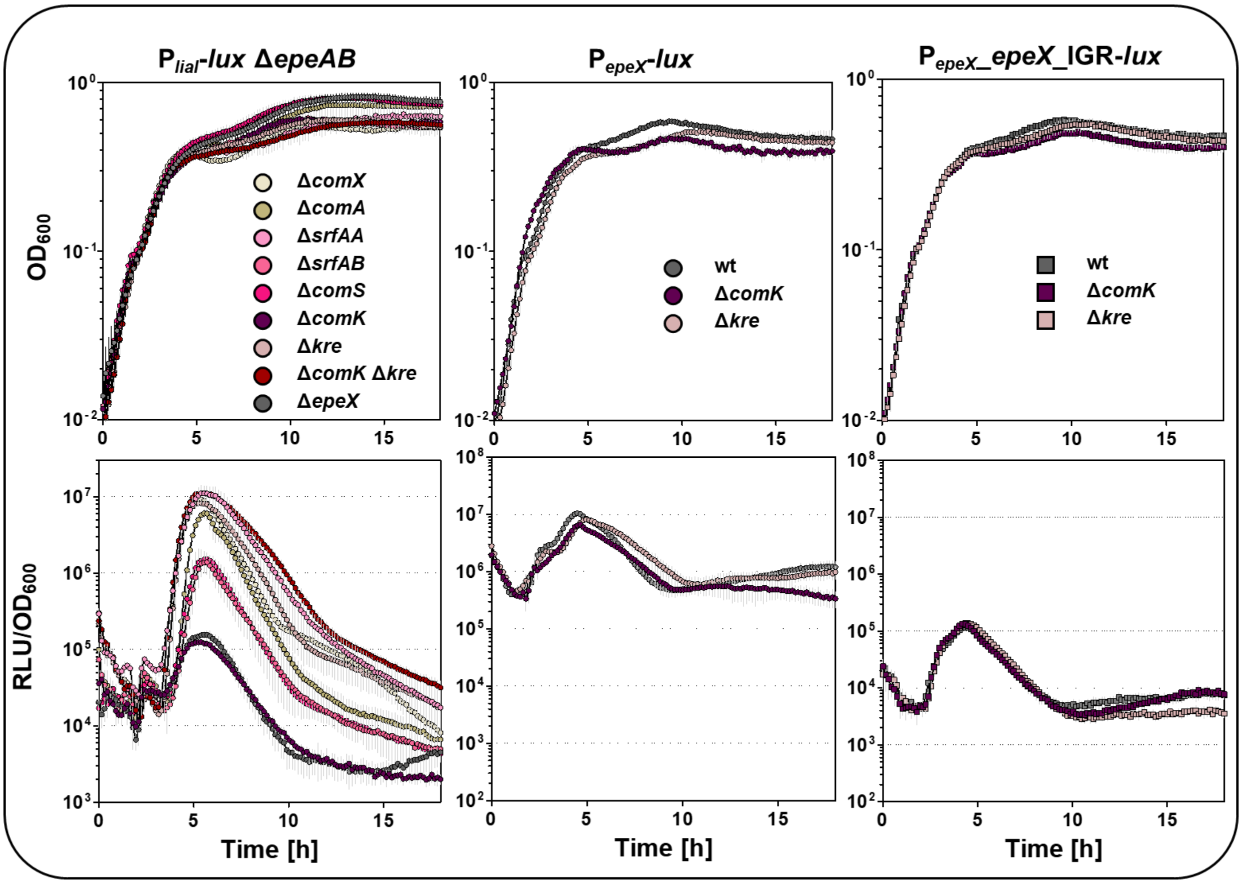


**Figure S4**: Impact of competence related mutants on EPE production and stress response.

While the upper graphs depict the growth curve as function of $OD_{600}$ over time, the lower graphs show the RLU values normalised to the corresponding $OD_{600}$ over time. The dynamic of the $P_{\textit{liaI}}$-*lux epeAB*, $P_{\textit{epeX}}$-*lux*, and $P_{\textit{epeX}}$_*epeX*_IGR-*lux* reporter activity in absence and presence of competence associated gene deletions were displayed. The standard derivation of biological and technical triplicates was included as error bars to each time point of measurement. For simplification, gene deletions are indicated with the delta symbol (Δ) throughout the figure, although this does not necessarily imply clean deletion strains. The full genotypes and respective resistance cassettes are provided in Table S1.

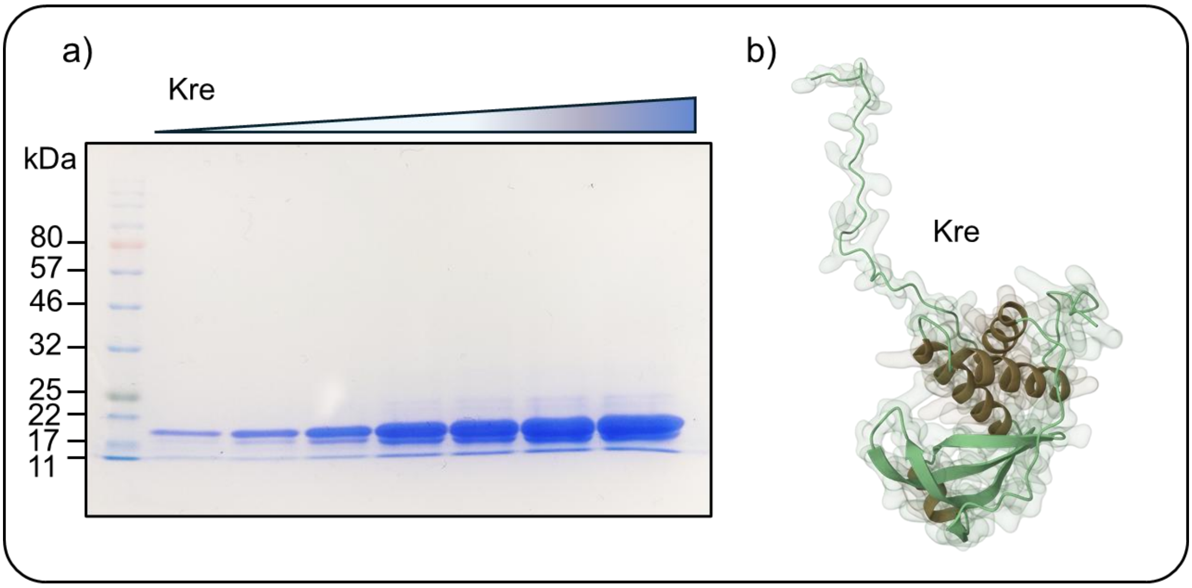


**Figure S5**: Purification and structural prediction of the RNA-binding protein Kre.

**a)** SDS-PAGE analysis of purified Kre protein after immobilised metal affinity chromatography (IMAC) using C-terminal His-tag, followed by size exclusion chromatography. Several dilutions in increasing amounts were loaded on the denaturation gel. The protein concentration was determined as 1.56 mg $mL^{-1}$ in 19% glycerol by Bradford reaction. These correspond to 10, 20, 35, 70, 90, 175, and 350 pmol purified Kre per lane, respectively. The molecular mass of Kre is 17.75 kDa. **b)** Predicted three-dimensional structure of Kre generated with AlphaFold.

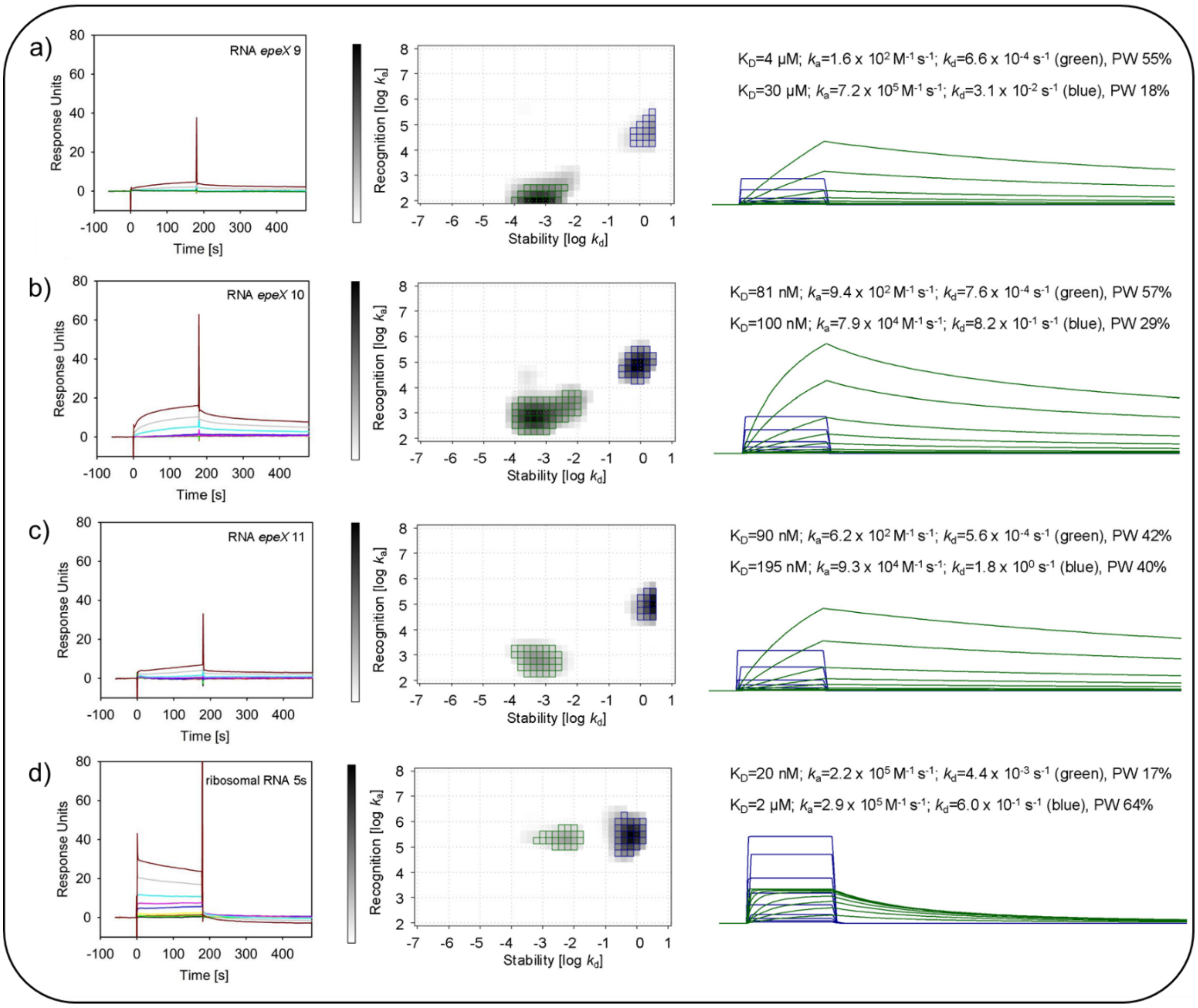


**Figure S6**: SPR analysis of Kre binding to truncated *epeX* RNA transcripts.

Sensorgrams (left panels) depict response units of Kre to increasing concentrations of *epeX* RNA fragments (black: 10 nM; red: 25 nM; light green: 50 nM; yellow: 100 nM; blue: 250 nM; turquoise: 500 nM; light blue: 1000 nM; grey: 2500 nM; brown: 5000 nM). InteractionMaps (central panels) show the distribution of association and dissociation rate constants as stability (dissociation rate; log $k_d$) and recognition (association rate, log $k_a$). The corresponding calculated sensorgrams from the IM peaks (right panels) are presented on the right, the colours of the curved correspond to the repectivs peaks from the IM analyses. The data were quantified and the respective overall affinities ($K_D$) calculated from the respective association ($k_a$) and dissociation rates ($k_d$) are shown, as well as the peak weights (PW) showing the overall contribution of the respective interaction towards the total sensorgrams given in (%). Since bulk binding are not included in the calculations, 100% were not reached in total. Short truncated versions of the epeX RNA **a)** epeX 9, **b)** epeX10, **c)** epeX 11, and **d)** the 5s (80 bp) RNA as control, were applied for SPR analysis.

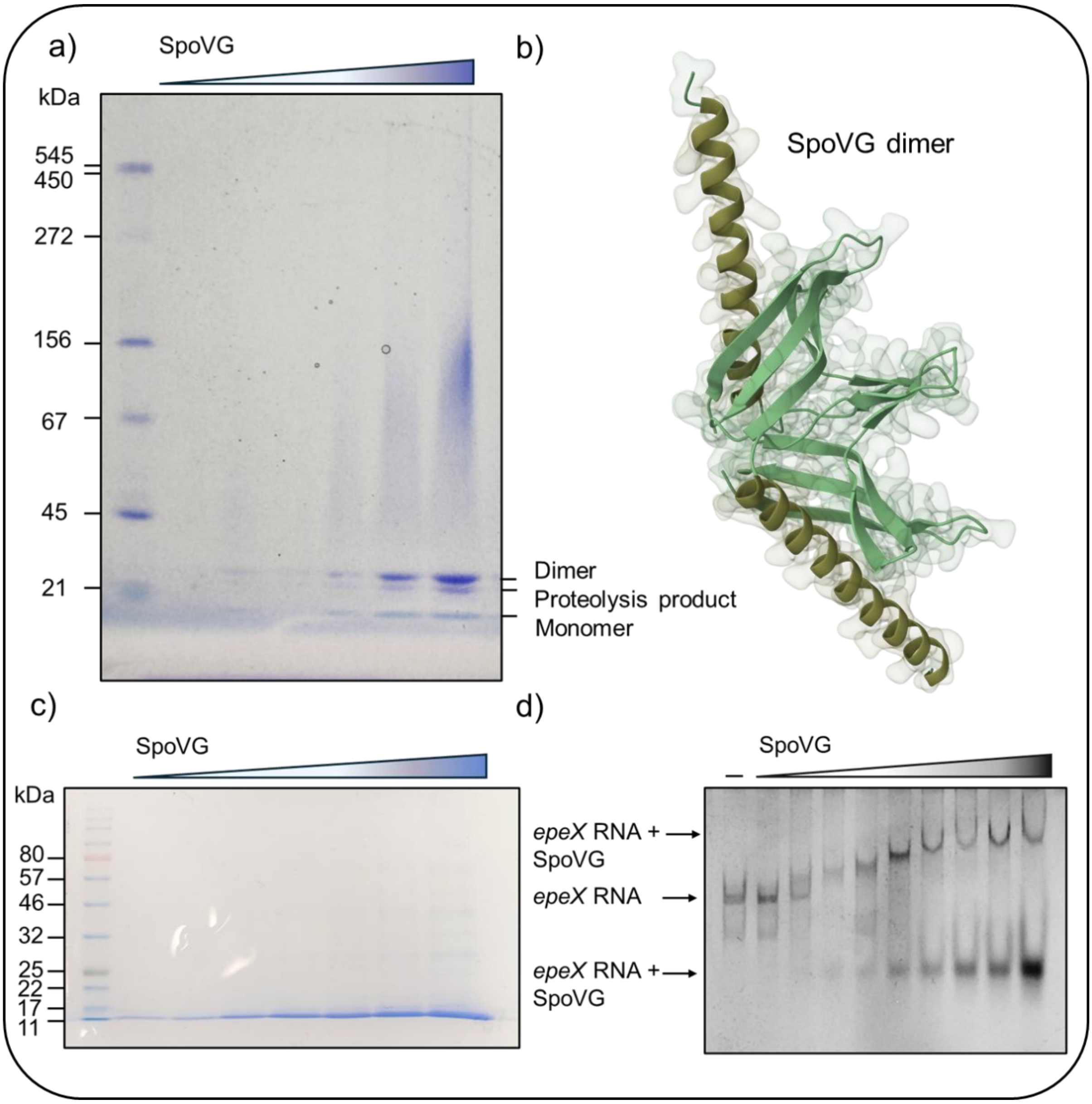


**Figure S7**: Structural characterisation of the RNA binding protein SpoVG and its interaction with the *epeX* RNA.

**a)** Native PAGE of SpoVG showing oligomeric states. Protein bands corresponding to the monomer (10.75 kDa), dimer, and a proteolytic fragment are visible. Protein concentration was calculated as 1.71 µg $mL^{-1}$ in 19% glycerol by Bradford reaction and SpoVG protein was loaded on the gel in increasing concentrations reaching from 15, 30, 60, 130, 160, and 320 pmol, respectively. **b)** Predicted dimeric structure of SpoVG generated using AlphaFold. **c)** SDS-PAGE analysis of purified Kre protein after immobilised metal affinity chromatography (IMAC) using C-terminal His-tag, followed by size exclusion chromatography. Increasing amounts of purified SpoVG were loaded on the denaturation gel (15, 30, 60, 130, 160, 320, and 640 pmol, respectively). **d)** Electrophoretic mobility shift assay (EMSA) showing binding of SpoVG to *epeX* RNA. Two distinct shifts are visible, indicating the formation of at least two RNA–protein complexes, one with slower and one with faster mobility relative to the free RNA as indicated by arrows.

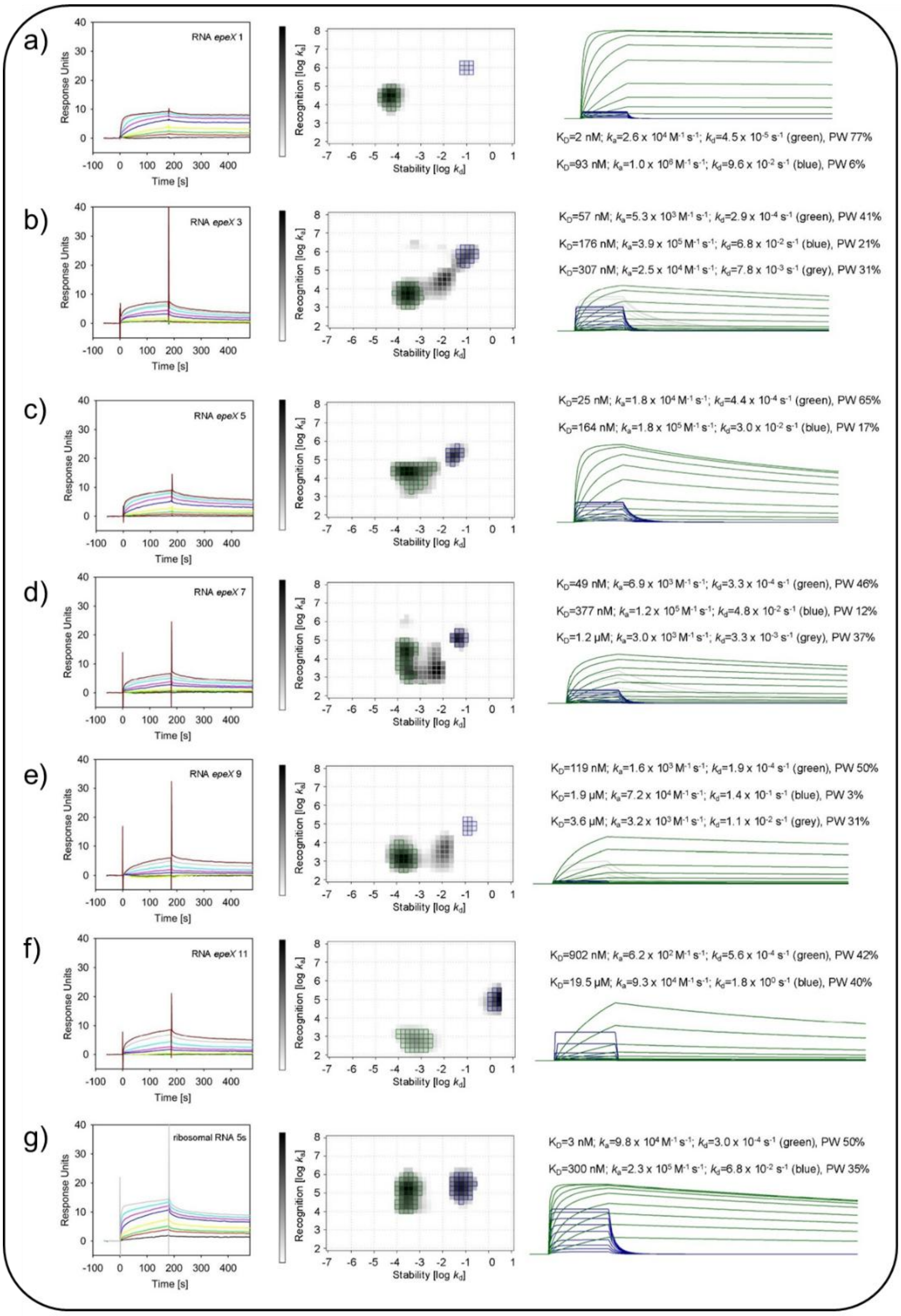


**Figure S8**: SPR analysis of SpoVG binding to truncated *epeX* RNA transcripts.

Sensorgrams (left panels) depict response units of SpoVG to increasing concentrations of *epeX* RNA fragments (black: 10 nM; red: 25 nM; light green: 50 nM; yellow: 100 nM; blue: 250 nM; turquoise: 500 nM; light blue: 1000 nM; grey: 2500 nM; brown: 5000 nM). InteractionMaps (central panels) show the distribution of association and dissociation rate constants as stability (dissociation rate; log $k_d$) and recognition (association rate, log $k_a$). The corresponding calculated sensorgrams from the IM peaks (right panels) are presented on the right, the colours of the curved correspond to the respective peaks from the IM analyses. The data were quantified and the respective overall affinities ($K_D$) calculated from the respective association ($k_a$) and dissociation rates ($k_d$) are shown, as well as the peak weights (PW) showing the overall contribution of the respective interaction towards the total sensorgrams given in (%). Since bulk binding is not included in the calculations, 100% were not reached in total. Short truncated versions of the *epeX* RNA **a)** *epeX* 1, **b)** *epeX* 3, **c)** *epeX* 5, **d)** *epeX* 7, **e)** *epeX* 9, **f)** *epeX* 11, and **g)** the 5s (80 bp) RNA as control, were applied for SPR analysis.

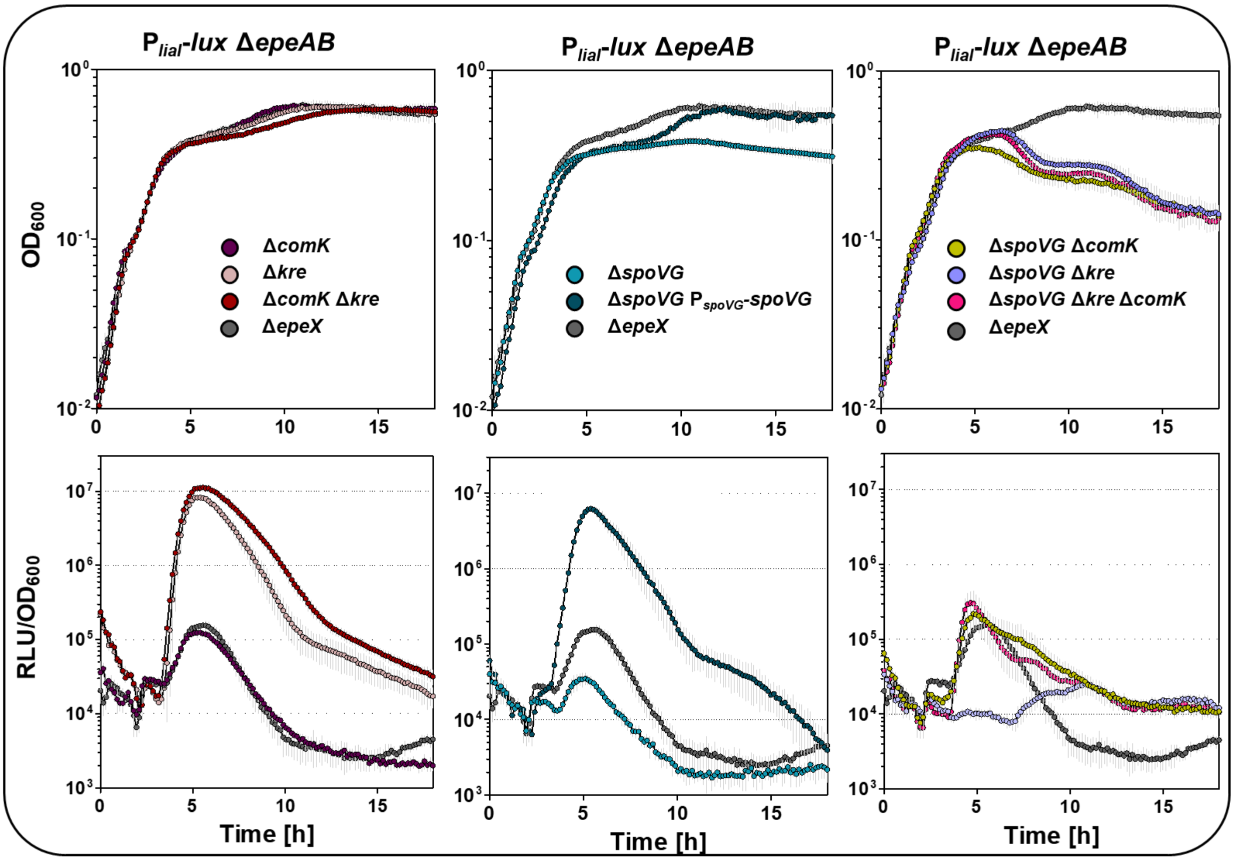


**Figure S9**: Effects of *comK*, *kre*, and *spoVG* single and combinatory mutants on EPE-mediated stress response.

While the upper graphs depict the growth curve as function of $OD_{600}$ over time, the lower graphs show the RLU values normalised to the corresponding $OD_{600}$ over time. The dynamic of the $P_{liaI}$-*lux epeAB*, reporter activity presence of *comK, kre,* and *spoVG* gene deletions and corresponding combinatorics was displayed. The standard derivation of biological and technical triplicates was included as error bars to each time point of measurement. For simplification, gene deletions are indicated with the delta symbol (Δ) throughout the figure, although this does not necessarily imply clean deletion strains. The full genotypes and respective resistance cassettes are provided in Table S1.

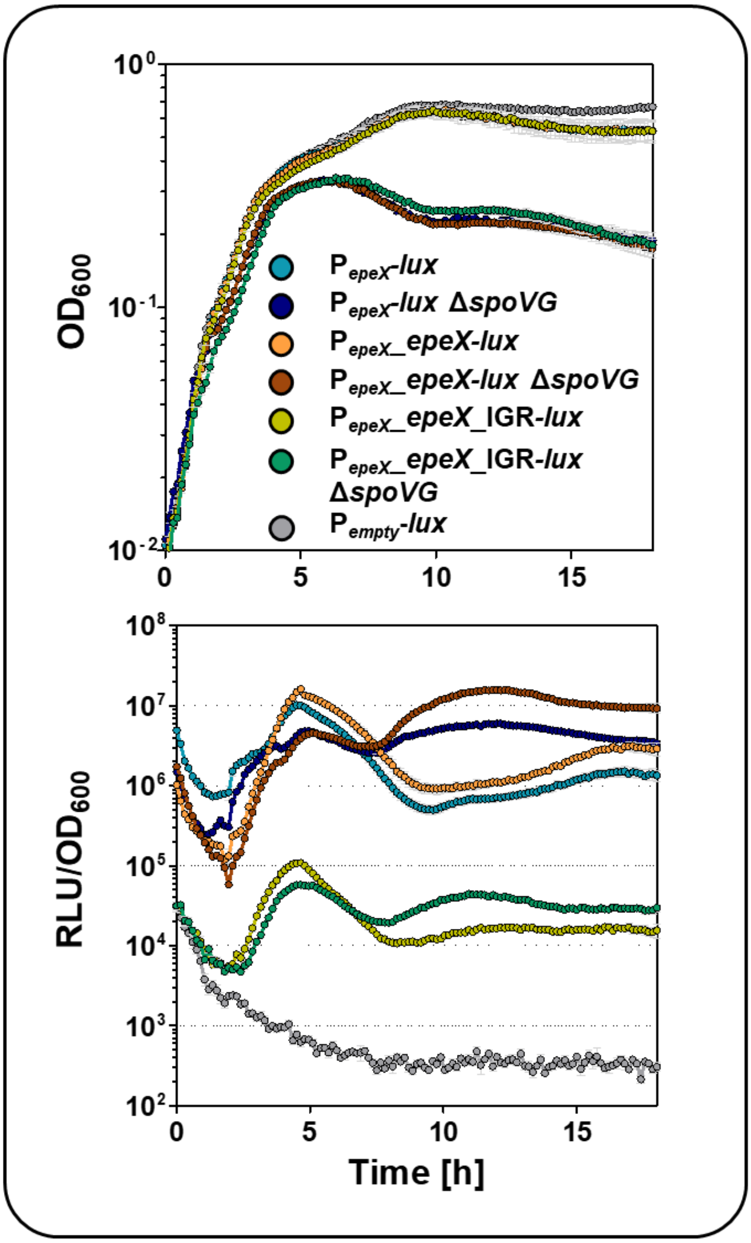


**Figure S10**: Effect of *spoVG* deletion on *epeXEP* expression.

While the upper graphs depict the growth curve as function of $OD_{600}$ over time, the lower graphs show the RLU values normalised to the corresponding $OD_{600}$ over time. The dynamic of the $P_{epeX}$-*lux*, $P_{epeX}$_*epeX-lux*, and $P_{epeX}$_*epeX*_IGR-*lux* reporter activity in absence and presence and *spoVG* gene deletionwas displayed. The standard derivation of biological and technical triplicates was included as error bars to each time point of measurement. For simplification, gene deletions are indicated with the delta symbol (Δ) throughout the figure, although this does not necessarily imply clean deletion strains. The full genotypes and respective resistance cassettes are provided in Table S1.

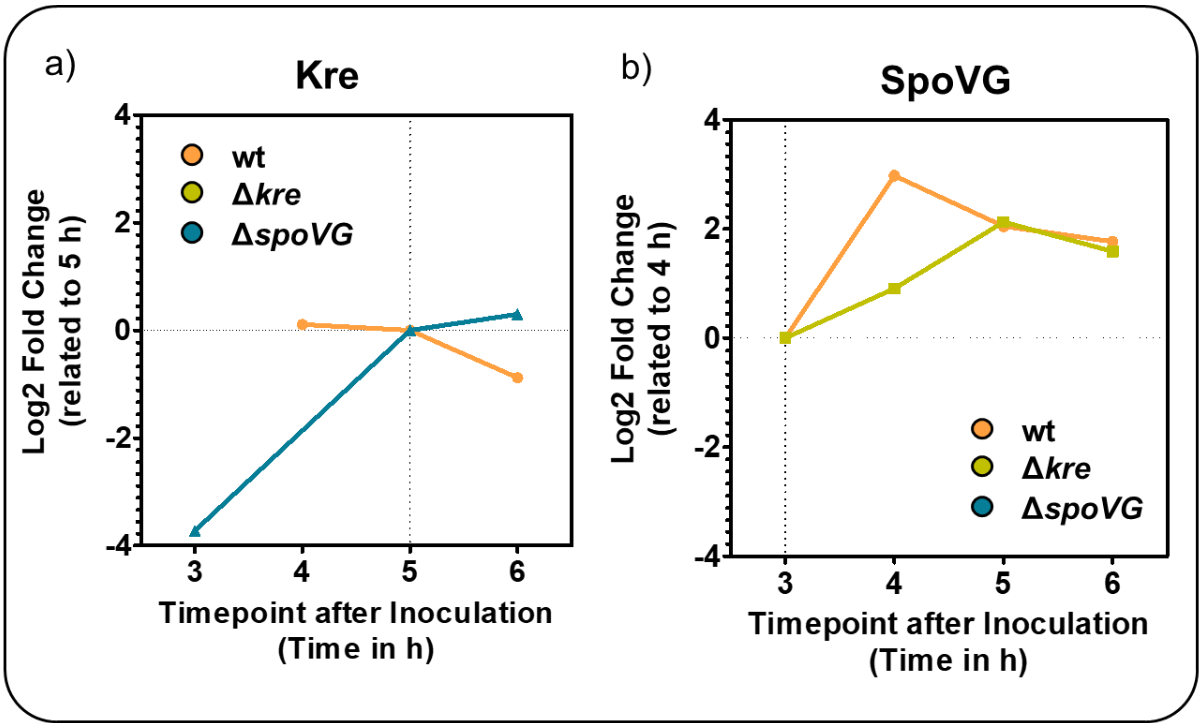


**Figure S11**: Temporal changes in Kre and SpoVG abundance during cultivation in DSM medium.

**a)** The $\log_2$ fold change of Kre abundance at 3, 4, 5, and 6 h relative to the 5 h time point, calculated for each strain: wild type, *kre* and *spoVG* mutant. **b)** $\text{Log}_2$ fold change of SpoVG abundance over time relative to the 3 h time point, calculated separately for each strain. Mass spectrometry was performed with biological triplicates. Note that fold changes represent relative changes within each strain. The absolute EpeX and EpeE abundances at the respective reference time points are not depicted.

#### References


1. Konkol, M.A., Blair, K.M. and Kearns, D.B. (2013) Plasmid-Encoded ComI Inhibits Competence in the Ancestral 3610 Strain of *Bacillus subtilis*, *J. Bacteriol.,* **195,** 4085–4093.

2. Popp, P.F., Dotzler, M. and Radeck, J. *et al.* (2017) The *Bacillus* BioBrick Box 2.0: Expanding the Genetic Toolbox for the Standardized Work With *Bacillus subtilis*, *Sci. Rep.,* **7,** 15058. First published on Nov 8, 2017.

3. Radeck, J., Kraft, K. and Bartels, J. *et al.* (2013) The *Bacillus* BioBrick Box: Generation and Evaluation of Essential Genetic Building Blocks for Standardized Work with *Bacillus subtilis*, *J. Biol. Eng.,* **7**.

4. Popp, P.F., Friebel, L. and Benjdia, A. *et al.* (2021) The Epipeptide Biosynthesis Locus *epeXEPAB* Is Widely Distributed in Firmicutes and Triggers Intrinsic Cell Envelope Stress, *Microb. physiol.,* **31,** 306–318. First published on Jun 11, 2021.

5. Radeck, J., Gebhard, S. and Orchard, P.S. *et al.* (2016) Anatomy of the Bacitracin Resistance Network in *Bacillus subtilis*, *Mol. Micro.,* **100,** 607–620.

6. Arnaud, M., Chastanet, A. and Débarbouillé, M. (2004) New Vector for Efficient Allelic Replacement in Naturally Nontransformable, Low-GC-Content, Gram-Positive Bacteria, *Appl. Environ. Microbiol.,* **70,** 6887–6891.
